# Detecting and quantifying natural selection at two linked loci from time series data of allele frequencies with forward-in-time simulations

**DOI:** 10.1101/562967

**Authors:** Zhangyi He, Xiaoyang Dai, Mark Beaumont, Feng Yu

**Author notes:** Corresponding author. Email addresses (Zhangyi He), (Feng Yu). MRC Toxicology Unit, University of Cambridge, Leicester LE1 7HB, United Kingdom &.

## Abstract

Recent advances in DNA sequencing techniques have made it possible to monitor genomes in great detail over time. This improvement provides an opportunity for us to study natural selection based on time serial samples of genomes while accounting for genetic recombination effect and local linkage information. Such genomic time series data allow for more accurate estimation of population genetic parameters and hypothesis testing on the recent action of natural selection. In this work, we develop a novel Bayesian statistical framework for inferring natural selection at a pair of linked loci by capitalising on the temporal aspect of DNA data with the additional flexibility of modelling the sampled chromosomes that contain unknown alleles. Our approach is based on a hidden Markov model where the underlying process is a two-locus Wright-Fisher diffusion with selection, which enables us to explicitly model genetic recombination and local linkage. The posterior probability distribution for the selection coefficients is obtained by using the particle marginal Metropolis-Hastings algorithm, which allows us to efficiently calculate the likelihood. We evaluate the performance of our Bayesian inference procedure through extensive simulations, showing that our method can deliver accurate estimates of selection coefficients, and the addition of genetic recombination and local linkage brings about significant improvement in the inference of natural selection. We illustrate the utility of our approach on real data with an application to ancient DNA data associated with white spotting patterns in horses.

## 1. Introduction

Natural selection is a fundamental evolutionary process that maintains function and drives adaptation, thereby altering patterns of diversity at the genetic level. Methods for detecting and quantifying natural selection have important applications such as identifying the genetic basis of diseases and understanding the molecular basis of adaptation. There has been a long line of theoretical and experimental research devoted to modelling and inferring natural selection, and the vast majority of earlier analyses are based on allele frequency data obtained at a single time point that requires unrealistic assumptions of ancestral demography and selective regimes (see Bank et al., 2014, for a review). With the advances in DNA sequencing technologies, an ever-increasing amount of allele frequency data sampled at multiple time points are becoming available. For example, such time series data arise from experimental evolution (*e*.*g*., Burke et al., 2010; Orozco-terWengel et al., 2012; Lang et al., 2013; Wiser et al., 2013), viral/phage populations (*e*.*g*., Wichman et al., 1999, 2005; Holder & Bull, 2001; Bollback & Huelsenbeck, 2007), or ancient DNA (aDNA) (*e*.*g*., Hummel et al., 2005; Ludwig et al., 2009; Orlando et al., 2013; Mathieson et al., 2015). Temporally spaced samples provide much more valuable information regarding natural selection since expected changes in allele frequencies over time are closely related to the strength of natural selection acting on the population. One can therefore expect allele frequency time series data to improve our ability to estimate selection coefficients and test hypotheses regarding natural selection.

There has been a growing literature on the statistical inference of natural selection from time series data of allele frequencies over the past decade (*e*.*g*., Bollback et al., 2008; Malaspinas et al., 2012; Mathieson & McVean, 2013; Steinrücken et al., 2014; Lacerda & Seoighe, 2014; Feder et al., 2014; Foll et al., 2014, 2015; Terhorst et al., 2015; Schraiber et al., 2016; Shim et al., 2016; Ferrer-Admetlla et al., 2016; Paris et al., 2019; He et al., 2019), reviewed in Bank et al. (2014) and Malaspinas (2016). A common approach to analysing time series data of allele frequencies is based upon the hidden Markov model (HMM) framework of Williamson & Slatkin (1999), where the underlying population is assumed to evolve according to the Wright-Fisher model introduced by Fisher (1922) and Wright (1931), and the observations are modelled through independent binomial sampling from the underlying population at each given time point (see Tataru et al., 2017, for a detailed review of the statistical inference in the Wright-Fisher model using allele frequency data). However, such approaches can become computationally infeasible for large populations because they require a prohibitively large amount of computation and storage for the calculation of the likelihood. Most existing HMM-based methods are therefore built on either the diffusion approximation of the Wright-Fisher model (*e*.*g*., Bollback et al., 2008; Malaspinas et al., 2012; Steinrücken et al., 2014; Schraiber et al., 2016; Ferrer-Admetlla et al., 2016; He et al., 2019) or the moment-based approximation of the Wright-Fisher model (*e*.*g*., Lacerda & Seoighe, 2014; Feder et al., 2014; Terhorst et al., 2015; Paris et al., 2019). Such approximations enable efficient integration over all possible allele frequency trajectories of the underlying population, thereby allowing the likelihood computation to be completed in a reasonable amount of time.

The recent advent of high-throughput sequencing technologies has made it possible to monitor genomes in great detail over time. This provides an opportunity for detecting and estimating natural selection at multiple linked loci from time series data of allele frequencies while taking the process of genetic recombination and the information of local linkage into account. Properly accounting for genetic recombination and local linkage can be expected to provide more precise estimates for the selection coefficient and more accurate hypothesis testing on the recent action of natural selection since genetic recombination may either reinforce or oppose changes in allele frequencies caused by natural selection according to the levels of linkage disequilibrium (He et al., 2020). However, with the exception of Terhorst et al. (2015), all existing methods built on the Wright-Fisher model for inferring natural selection from allele frequency time series data are limited to either a single locus (*e*.*g*., Bollback et al., 2008; Malaspinas et al., 2012; Steinrücken et al., 2014; Schraiber et al., 2016; Paris et al., 2019; He et al., 2019) or multiple independent loci (*e*.*g*., Foll et al., 2014, 2015; Shim et al., 2016; Ferrer-Admetlla et al., 2016), *i*.*e*., genetic recombination effect and local linkage information are ignored in these methods. The exception amongst these approaches, Terhorst et al. (2015), extended a moment-based approximation of the Wright-Fisher model introduced by Feder et al. (2014) to multiple linked loci with an application to the pooled sequencing (Pool-Seq) data from evolve-and-resequence (E&R) experiments, where the allele frequency transition between two consecutive sampling time points is modelled deterministically, with added Gaussian noise.

In the present work, we propose a novel HMM-based method for Bayesian inference of natural selection at two linked loci from time series data of allele frequencies while accounting for the process of genetic recombination, thereby incorporating the information on local linkage. The key innovation is that a diffusion approximation to the Wright-Fisher model of the stochastic evolutionary dynamics under natural selection at two linked loci is used as the hidden Markov process to characterise the changes in the haplotype frequencies of the underlying population over time, which enables us to explicitly model genetic recombination and local linkage. The diffusion approximation we use in our method allows us to avoid the restriction that the allele frequency trajectory of the underlying population remains far away from allele fixation or loss, which was imposed by the Gaussian approximation used in Terhorst et al. (2015). Our posterior computation is carried out with the particle marginal Metropolis-Hastings (PMMH) algorithm developed by Andrieu et al. (2010), which enables us to efficiently calculate the likelihood. Also, our method can handle sampled chromosomes with unknown alleles, which is common in aDNA data due to postmortem damage. In addition, our approach can be readily extended to model a range of complex evolutionary scenarios like non-constant demographic histories.

We evaluate the performance of our Bayesian inference procedure through extensive simulations. We illustrate that our approach allows for efficient and accurate estimation of selection coefficients from allele frequency time series data, regardless of whether sampled chromosomes contain unknown alleles or not. We present two scenarios where existing single-locus methods fail to deliver precise estimates for selection coefficients whereas our approach still works well, especially when the loci are tightly linked. This shows the efficacy of our method in modelling genetic recombination and local linkage. Finally, we apply our Bayesian inference procedure to analyse the aDNA data associated with white spotting patterns in horses from Wutke et al. (2016) and find that in horses there is no evidence showing that the tobiano pattern is positively selected but strong evidence of the sabino pattern being negatively selected.

## 2. Materials and Methods

In this section, we begin with a short review of the Wright-Fisher diffusion for two linked loci evolving under natural selection over time, and then describe our Bayesian procedure for the inference of natural selection at the two linked loci from temporally spaced samples, *e*.*g*., how to set up the HMM framework and how to compute the posterior probability distribution for the population genetic quantities of interest with Markov chain Monte Carlo (MCMC) techniques.

### 2.1. Wright-Fisher diffusion

Consider a diploid population of randomly mating individuals at two linked loci 𝒜 and ℬ evolving under natural selection according to the two-locus Wright-Fisher model with selection (see, *e.g*., He et al., 2017), for which we assume discrete time and nonoverlapping generations. At each locus, there are two possible allele types, labelled 𝒜_1_, 𝒜_2_ and ℬ_1_, ℬ_2_, respectively, resulting in four possible haplotypes 𝒜_1_ℬ_1_, 𝒜_1_ℬ_2_, 𝒜_2_ℬ_1_ and 𝒜_2_ℬ_2_, labelled haplotypes 1, 2, 3 and 4, respectively. We attach symbols 𝒜_1_ and ℬ_1_ to the mutant alleles, which are assumed to arise only once in the population and be positively selected once it exists, and we attach symbols 𝒜_2_ and ℬ_2_ to the ancestral alleles, which are assumed to originally exist in the population.

We incorporate viability selection into the population dynamics and assume that the viability is fixed from the time that the mutant allele arises in the population and is only determined by the genotype at each single locus. More specifically, we assume that the relative viabilities of the sixteen possible (ordered) genotypes at the two loci are determined multiplicatively from the relative viabilities at individual loci, and the relative viabilities of the three possible genotypes at each single locus, *e.g*., genotypes 𝒜_1_𝒜_1_, 𝒜_1_𝒜_2_ and 𝒜_2_𝒜_2_ at a given locus 𝒜, are taken to be 1, 1 − *h*_𝒜_*s*_𝒜_ and 1 − *s*_𝒜_, respectively, where *s*_𝒜_ ∈ [0, 1] is the selection coefficient and *h*_𝒜_ ∈ [0, 1] is the dominance parameter. For example, the relative viability of the 𝒜_1_ℬ_2_*/*𝒜_2_ℬ_2_ genotype is (1 − *h*_𝒜_*s*_𝒜_)(1 − *s*_ℬ_). We let *r* ∈ [0, 0.5] be the recombination rate of the two loci on the same chromosome (*i.e*., the fraction of recombinant offspring showing a crossover between the two loci). We assume that the population size is fixed to be *N* diploid individuals for all time.

#### 2.1.1. Two-locus Wright-Fisher diffusion with selection

We consider a scaling limit of the Wright-Fisher model, where the unit of time is rescaled by 2*N*. The scaled selection coefficients *α*_𝒜_ = 2*Ns*_𝒜_ and *α*_ℬ_ = 2*Ns*_ℬ_, and the scaled recombination rate *ρ* = 4*Nr* are kept constant while the population size *N* is taken to infinity. As the population size approaches infinity, the haplotype frequency trajectories follow a standard diffusion limit of the two-locus Wright-Fisher model with selection (see, *e.g*., He et al., 2020), called the two-locus Wright-Fisher diffusion with selection. The Wright-Fisher diffusion has already been successfully applied in the inference of natural selection from allele frequency time series data. The partial differential equation (PDE) satisfied by the transition probability density function of the Wright-Fisher diffusion was used in *e.g*., Bollback et al. (2008); Gutenkunst et al. (2009); Steinrücken et al. (2014); He et al. (2019). In this work, as used in *e.g*., Schraiber et al. (2016), we characterise the Wright-Fisher diffusion as the solution of the stochastic differential equation (SDE) instead.

We let *X*_*i*_(*t*) be the frequency of haplotype *i* in the population at time *t* for *i* = 1, 2, 3, 4, and denote the haplotype frequencies of the four possible types in the population by ***X***(*t*), which evolves in the state space (*i.e*., a three-simplex)

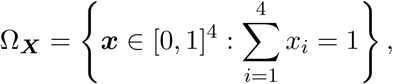

and satisfies the SDE in the following form

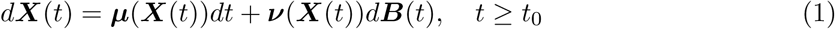

with initial condition ***X***(*t*_0_) = ***x***_0_. In Eq. (1), the drift term ***µ***(***x***) is a four-dimensional vector being

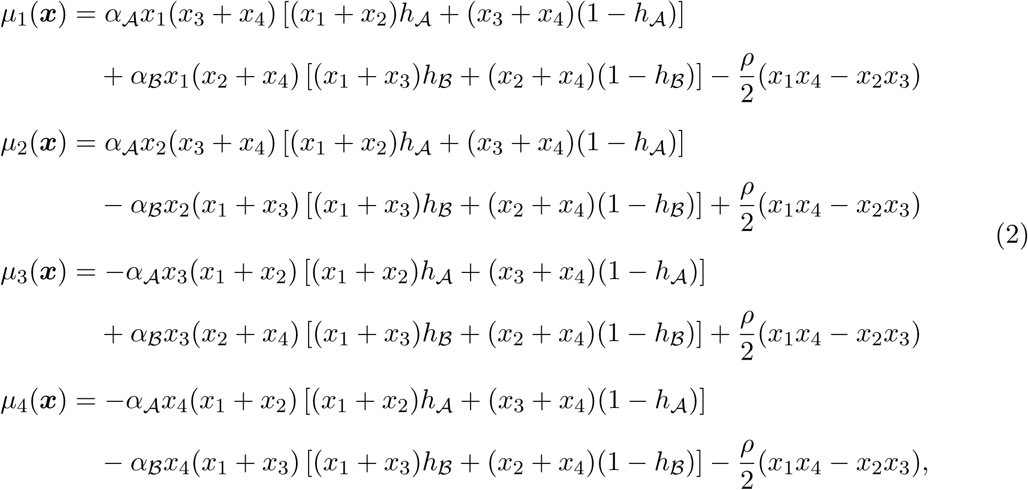

the diffusion term ***ν***(***x***) is a four by three matrix satisfying

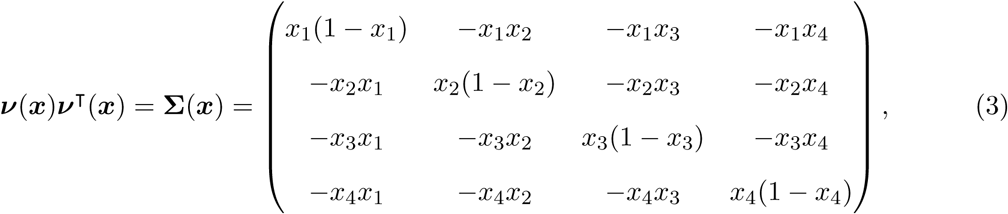

and ***B***(*t*) is a three-dimensional standard Brownian motion. The term *x*_1_*x*_4_ − *x*_2_*x*_3_ in Eq. (2) is a measure of the linkage disequilibrium between loci 𝒜 and ℬ, which quantifies the non-random association of the alleles at the two loci.

#### 2.1.2. Forward-in-time simulation of the Wright-Fisher diffusion

To obtain a numerical solution of the Wright-Fisher SDE in Eq. (1), we need to compute the diffusion term ***ν***(***x***), which we have to perform at each time step in most existing numerical simulation schemes. The diffusion term ***ν***(***x***) can be analytically derived with the Cholesky decomposition (Sato, 1976), which however, explodes at the boundaries. There exist other matrix decompositions capable of computing the diffusion term ***ν***(***x***) such as spectral decomposition, which are valid for positive semi-definite matrices, typically at the expense of either additional numerical errors and computational costs, or limitations in applicability to the infinitesimal covariance matrix **Σ**(***x***) of the form in Eq. (3).

Following He et al. (2020), we reformulate the Wright-Fisher SDE in the following form

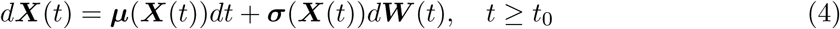

with initial condition ***X***(*t*_0_) = ***x***_0_, where the diffusion term ***σ***(***x***) can be explicitly written down as

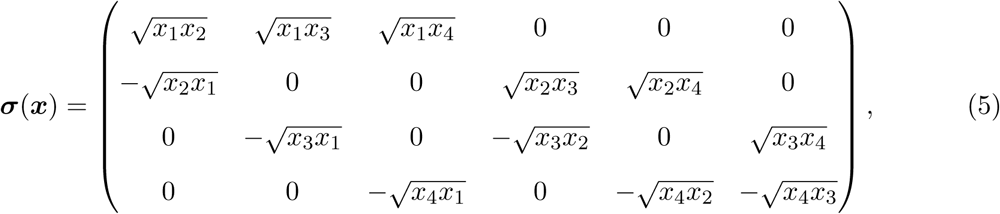

and ***W*** (*t*) is a six-dimensional standard Brownian motion. Combining Eqs. (3) and (5), we have

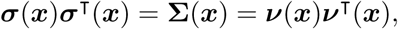

which implies that the two Wright-Fisher SDEs have the same infinitesimal generator

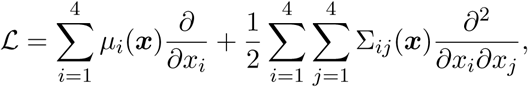

thus having the same weak solution. This guarantees that we can achieve the numerical solution of the Wright-Fisher SDE of the form in Eq. (1) by numerically solving the Wright-Fisher SDE of the form in Eq. (4), which enables us to avoid boundary issues and reduce computational costs.

There exist a number of numerical simulation schemes for SDEs (see Kloeden & Platen, 1992, for an excellent introduction). The numerical approach we adopt in this work is the commonly used Euler-Maruyama scheme, one of the most popular numerical methods for SDEs in practice due to its high efficiency and low complexity. More specifically, we divide each generation into *L* subintervals by setting Δ*t* = 1*/*(2*NL*), and then the Euler-Maruyama approximation of the Wright-Fisher diffusion can be formulated as

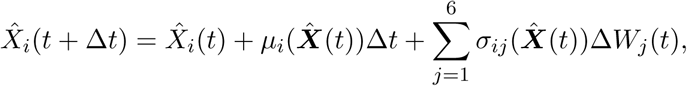

for *i* = 1, 2, 3, 4, where Δ*W*_*j*_(*t*) = *W*_*j*_(*t* + Δ*t*) − *W*_*j*_(*t*) are independent and normally distributed with mean 0 and variance Δ*t* for *j* = 1, 2, …, 6. The Euler-Maruyama scheme is numerically stable and strongly consistent (see, *e.g*., Kloeden & Platen, 1992), and the convergence of the Euler-Maruyama approximation of the Wright-Fisher diffusion is guaranteed by Zhang (2006).

### 2.2. Bayesian inference of natural selection

Suppose that the available data are always sampled from the underlying population at a finite number of distinct time points, say *t*_1_ < *t*_2_ < … < *t*_*K*_, where the time is measured in units of 2*N* generations to be consistent with the Wright-Fisher diffusion time scale. At the *k*-th sampling time point, we let 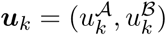 and 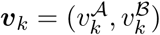 denote the counts of mutant alleles and ancestral alleles observed at loci 𝒜 and ℬ in the sample of *n*_*k*_ chromosomes drawn from the underlying population, respectively. The population genetic quantities of interest in the present work are the scaled selection coefficients *α*_𝒜_ and *α*_ℬ_, the dominance parameters *h*_𝒜_ and *h*_ℬ_, and the scaled recombination rate *ρ*, which are denoted by ***ϑ*** = (*α*_𝒜_, *h*_𝒜_, *α*_ℬ_, *h*_ℬ_, *ρ*).

#### 2.2.1. Hidden Markov model

Similar to Bollback et al. (2008), the underlying population is assumed to evolve according to the two-locus Wright-Fisher diffusion with selection in our HMM framework, and the observations are modelled as independent samples drawn from the underlying population at each sampling time point (see Figure 1 for the graphical representation of our HMM framework). To compute the posterior probability distribution *p*(***ϑ*** | ***u***_1:*K*_, ***v***_1:*K*_), we condition and integrate over all possible haplotype frequency trajectories of the underlying population at each sampling time point. More specifically, we let ***x***_1:*K*_ = {***x***_1_, ***x***_2_, …, ***x***_*K*_} denote the haplotype frequency trajectories of the underlying population at the sampling time points ***t***_1:*K*_. The posterior probability distribution for the population genetic quantities of interest can then be written as

**Figure 1:**
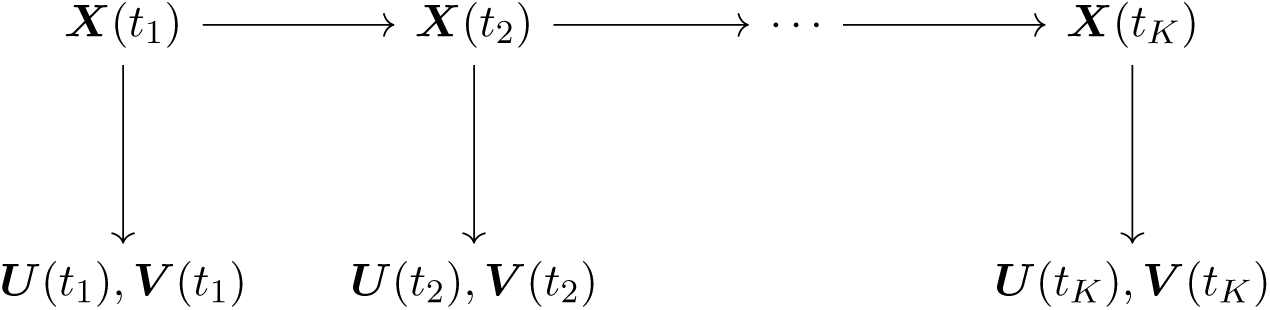
Graphical representation of the HMM framework for time series data of allele frequencies.

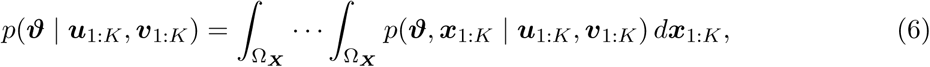

where

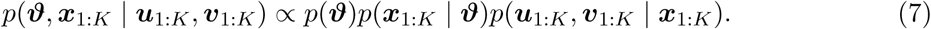

In Eq. (7), *p*(***ϑ***) is the prior probability distribution for the population genetic quantities of interest and can be taken to be a uniform prior over the parameter space if prior knowledge is poor, *p*(***x***_1:*K*_ | ***ϑ***) is the probability distribution for the haplotype frequency trajectories of the underlying population at the sampling time points ***t***_1:*K*_, and *p*(***u***_1:*K*_, ***v***_1:*K*_ | ***x***_1:*K*_) is the conditional probability for the observations at the sampling time points ***t***_1:*K*_ given the haplotype frequency trajectories of the underlying population.

Since the Wright-Fisher diffusion is a Markov process, the probability distribution for the haplotype frequency trajectories of the underlying population at the sampling time points ***t***_1:*K*_ can be decomposed as

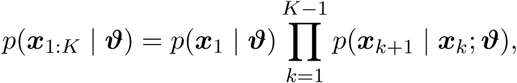

where *p*(***x***_1_ | ***ϑ***) is the prior probability distribution for the haplotype frequencies of the underlying population at the initial sampling time point and can be taken to be a uniform prior over the state space Ω_***X***_, known as the flat Dirichlet distribution, if prior knowledge is poor. The term in the product above, *p*(***x***_*k*+1_ | ***x***_*k*_; ***ϑ***), is the transition probability density of the Wright-Fisher diffusion between two consecutive sampling time points for *k* = 1, 2, …, *K* − 1, which can be obtained by numerically solving the Kolmogorov backward equation (or its adjoint) associated with the Wright-Fisher diffusion. However, this requires a fine enough discretisation of the state space Ω_***X***_, if a finite difference method is used, and strongly depends on the underlying population genetic parameters (Ragsdale & Gutenkunst, 2017). In addition, numerically solving such a PDE in three dimensions for our posterior computation is computationally challenging and prohibitively expensive. We therefore resort to an ‘exact-approximate’ Monte Carlo procedure (Andrieu & Vihola, 2016) in this work that involves simulating the Wright-Fisher SDE in the form of Eq. (4), as a tractable alternative.

Given the haplotype frequency trajectories of the underlying population, the observations at each sampling time point are independent of one another, which means that

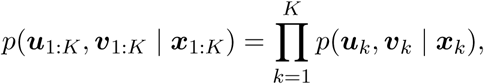

where *p*(***u***_*k*_, ***v***_*k*_ | ***x***_*k*_) is the conditional probability for the observations at the *k*-th sampling time point given the haplotype frequency trajectories of the underlying population for *k* = 1, 2, …, *K*. To calculate the emission probability *p*(***u***_*k*_, ***v***_*k*_ | ***x***_*k*_), we let ***z***_*k*_ = (*z*_1,*k*_, *z*_2,*k*_, *z*_3,*k*_, *z*_4,*k*_) denote the counts of the 𝒜_1_ℬ_1_, 𝒜_1_ℬ_2_, 𝒜_2_ℬ_1_ and 𝒜_2_ℬ_2_ haplotypes in the sample at the *k*-th sampling time point, which are usually unobserved (see Figure 2 for the graphical representation of our HMM framework incorporating the additional level of sampling noise). We then have

**Figure 2:**
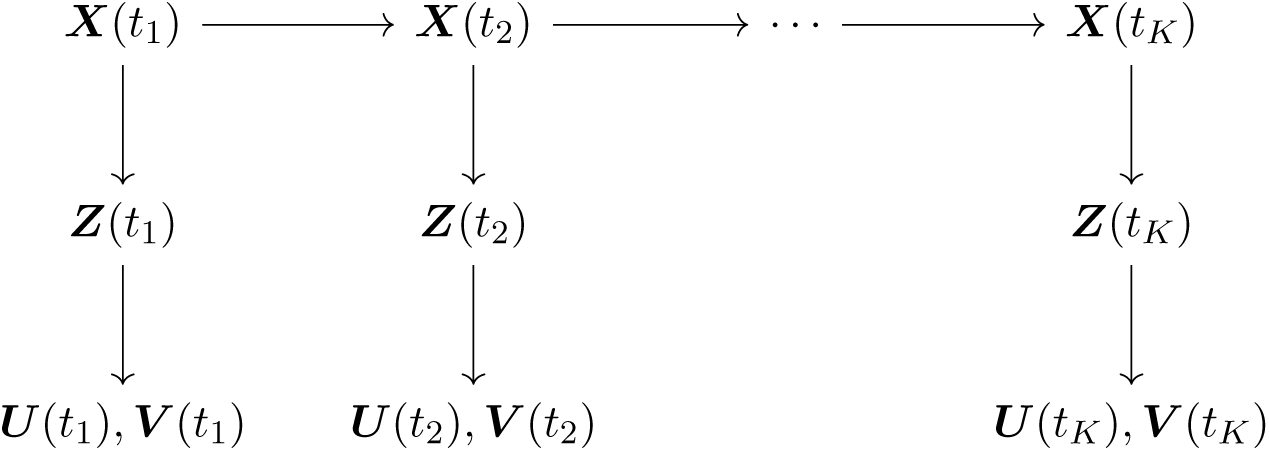
Graphical representation of the HMM framework for time series data of allele frequencies incorporating the additional level of sampling noise caused by the unobserved haplotype counts of the sample.

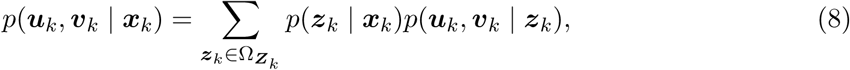

where

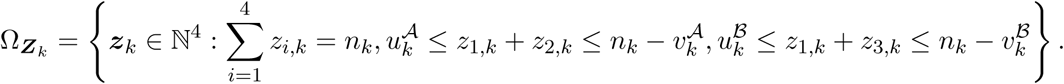

Conditional on the haplotype frequency trajectories of the underlying population at the *k*-th sampling time point, the haplotype counts of the sample can be modelled through multinomial sampling from the underlying population with sample size *n*_*k*_. We can then formulate the first term in the summation of Eq. (8) as

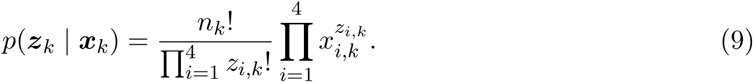

The second term in the summation of Eq. (8) can be decomposed as

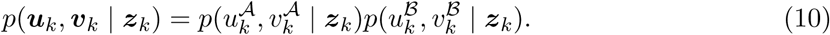

Let *ϕ* denote the probability that a sampled chromosome at a single locus is of unknown type, which we assume to be identical for all loci. We therefore have

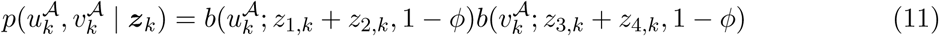

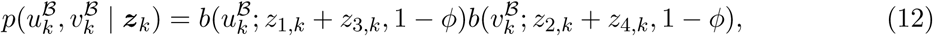

where

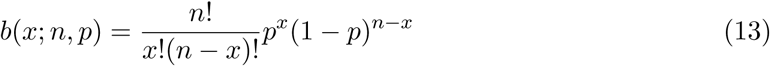

is the formula for the binomial distribution. The probability that the sampled chromosome at a single locus is of unknown type is usually unavailable, but we can estimate it with

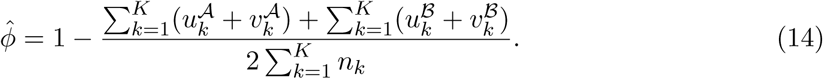

#### 2.2.2. Particle marginal Metropolis-Hastings

To compute the marginal posterior *p*(***ϑ*** | ***u***_1:*K*_, ***v***_1:*K*_), we resort to MCMC techniques since the posterior probability distribution in Eq. (6) is unavailable in a closed form. A Metropolis-Hastings (MH) scheme can be devised to explore the population genetic quantities of interest with a fairly arbitrary proposal probability distribution, *e.g*., a random walk proposal, where a sample of new candidates of the parameters ***ϑ***^*\**^ is drawn from the proposal *q*(***ϑ***^*\**^ | ***ϑ***) and is accepted with the Metropolis-Hastings ratio

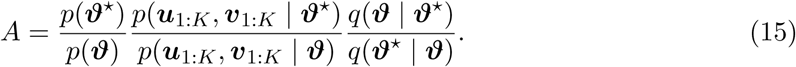

Our problem reduces to the calculation of the intractable marginal likelihood *p*(***u***_1:*K*_, ***v***_1:*K*_ | ***ϑ***) in Eq. (15), which can be formulated as

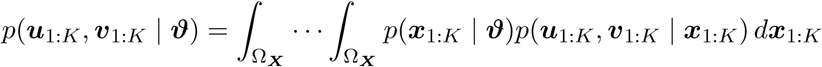

and achieved with a Monte Carlo (MC) estimate (Beaumont, 2003; Andrieu & Roberts, 2009). This pseudo-marginal MCMC algorithm exploits the fact that the MC estimate of the marginal likelihood *p*(***u***_1:*K*_, ***v***_1:*K*_ | ***ϑ***) is unbiased (or has a constant bias independent of the parameters ***ϑ***) and targets the marginal posterior *p*(***ϑ*** | ***u***_1:*K*_, ***v***_1:*K*_).

We adopt a closely related approach developed by Andrieu et al. (2010), which obtains an unbiased sequential Monte Carlo (SMC) estimate of the marginal likelihood *p*(***u***_1:*K*_, ***v***_1:*K*_ | ***ϑ***) and targets the joint posterior *p*(***ϑ, x***_1:*K*_ | ***u***_1:*K*_, ***v***_1:*K*_). This method is called particle marginal Metropolis-Hastings (PMMH) and permits a joint update of the population genetic quantities of interest and the latent population haplotype frequency trajectories. The co-estimation of the haplotype frequency trajectories of the underlying population is interesting in its own right, but our interest here lies only in the population genetic parameters. We therefore employ a special case of the PMMH algorithm in this work, where we do not generate and store the haplotype frequency trajectories of the underlying population in the state of the Markov chain. Full details about the PMMH algorithm can be found in Andrieu et al. (2010). Fearnhead & Künsch (2018) provided a detailed review of MC methods for estimating parameters in the HMM based on the particle filter.

In our Bayesian inference procedure, the implementation of the PMMH algorithm requires the SMC estimate of the marginal likelihood *p*(***u***_1:*K*_, ***v***_1:*K*_ | ***ϑ***). This can be achieved by the bootstrap particle filter introduced by Gordon et al. (1993) in the following manner. For the sampling time point *t*_1_, we first generate a sample of *M* particles, denoted by 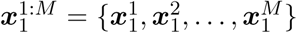, from the prior *p*(***x***_1_ | ***ϑ***) and assign each particle 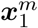a weight given by

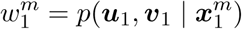

for *m* = 1, 2, …, *M*, where the superscript *m* denotes the particle label. We then calculate the SMC estimate of the marginal likelihood for the observations ***u***_1_ and ***v***_1_ by

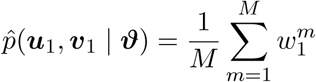

and resample *M* times with replacement from the sample of particles 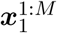 with the probabilities given by the normalised weights 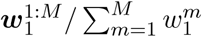. We repeat the following steps for the sampling time points ***t***_2:*K*_ :

Step 1: Generate each particle 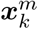by simulating the Wright-Fisher diffusion ***X***(*t*) on the time interval [*t*_*k*−1_, *t*_*k*_] starting at the frequency 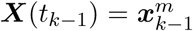 with the Euler-Maruyama scheme for *m* = 1, 2, …, *M*.

Step 2: Assign each particle 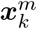a weight given by

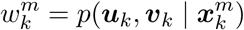

for *m* = 1, 2, …, *M*.

Step 3: Calculate the SMC estimate of the marginal likelihood for the observations ***u***_1:*k*_ and ***v***_1:*k*_ by

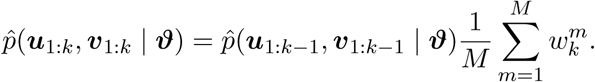

Step 4: Resample *M* times with replacement from the sample of particles 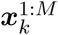 with the probabilities given by the normalised weights 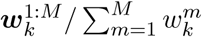.

Our Bayesian inference procedure then consists in the followings. We first generate a sample of initial candidates of the parameters ***ϑ*** from the prior *p*(***ϑ***) and then run a bootstrap particle filter with the proposed parameters ***ϑ*** to obtain the SMC estimate of the marginal likelihood 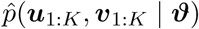. We repeat the following steps until a sufficient number of the samples of the parameters ***ϑ*** have been obtained:

Step 1: Generate a sample of new candidates of the parameters ***ϑ***^*\**^ from the proposal *q*(***ϑ***^*\**^ | ***ϑ***).

Step 2: Run a bootstrap particle filter with the proposed parameters ***ϑ***^*\**^ to obtain the SMC estimate of the marginal likelihood 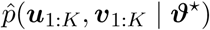.

Step 3: Accept the proposed parameters ***ϑ***^*\**^ with the Metropolis-Hastings ratio

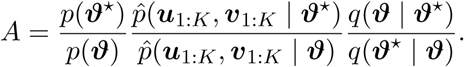

Once enough samples of the parameters ***ϑ*** have been obtained, we can get the minimum mean square error (MMSE) estimates for the population genetic quantities of interest, defined by

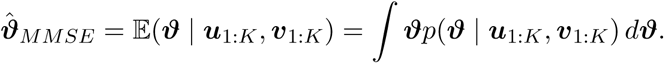

Alternatively, we can compute the posterior *p*(***ϑ*** | ***u***_1:*K*_, ***v***_1:*K*_) from the samples of the parameters ***ϑ*** using nonparametric density estimation techniques (see Izenman, 1991, for a detailed review) and achieve the maximum a posteriori probability (MAP) estimates for the population genetic quantities of interest, defined by

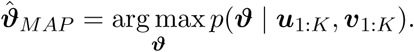

### 2.3. Data availability

Supplemental material available at FigShare. Source code for the method described in this work is available at https://github.com/zhangyi-he/WFM-2L-DiffusApprox-FwdPMMH.

## 3. Results

In this section, we show how our Bayesian inference method performs on simulated datasets with known population genetic parameter values, including a scenario where sampled chromosomes contain unknown alleles. We also present two examples to demonstrate the improvement in the inference of natural selection through explicitly modelling genetic recombination and local linkage. Finally, we apply our approach to the aDNA data associated with horse white spotting patterns from previous studies of Ludwig et al. (2009), Pruvost et al. (2011) and Wutke et al. (2016).

### 3.1. Analysis of simulated data

We run forward-in-time simulations of the two-locus Wright-Fisher model with selection and evaluate the performance of our approach on these replicate simulations by examining the bias and the root mean square error (RMSE) of our Bayesian estimates. In what follows, we take the dominance parameters to be *h*_𝒜_ = 0.5 and *h*_ℬ_ = 0.5 (*i.e*., the heterozygous fitness is the arithmetic average of the homozygous fitness, called genic selection) and choose a population size of *N* = 5000 unless otherwise noted. In principle, however, the conclusions hold for any other values of the dominance parameters *h*_𝒜_, *h*_ℬ_ ∈ [0, 1] and the population size *N* ∈ N.

For each simulated dataset, given the values of the population genetic parameters ***ϑ*** and the initial population haplotype frequencies ***x***_0_, we simulate the haplotype frequency trajectories of the underlying population according to the two-locus Wright-Fisher model with selection. After obtaining the simulated population haplotype frequency trajectories, we draw the unobserved sample haplotype counts independently at each sampling time point according to the multinomial distribution in Eq. (9) first and then we generate the observed sample mutant allele counts and ancestral allele counts with Eqs. (10)-(13).

#### 3.1.1. Power to infer natural selection

We vary the selection coefficients with *s*_𝒜_ ∈ {0.003, 0.01} and *s*_ℬ_ ∈ {0, 0.002, 0.008}, and the recombination rate with *r* ∈ {0.00001, 0.01} in our simulation studies. We perform 100 replicates for each of the 12 possible combinations of the selection coefficients and the recombination rate.

For each replicate, we set the initial population haplotype frequencies ***x***_0_ = (0.04, 0.08, 0.08, 0.8) and simulate the haplotype frequency trajectories of the underlying population according to the two-locus Wright-Fisher model with selection. We sample 50 chromosomes from the underlying population at every 50 generations throughout 500 generations.

We choose a uniform prior over the interval [−1, 1] for the selection coefficients *s*_𝒜_ and *s*_ℬ_, and a flat Dirichlet prior for the initial population haplotype frequencies ***x***_0_ in our Bayesian inference method. We divide each generation into 5 subintervals in the Euler-Maruyama scheme and run the PMMH algorithm with 1500 particles and 10000 iterations. We discard the initial 2000 iterations as the burn-in period and then thin the remaining PMMH output by selecting every fourth value.

The resulting boxplots of the empirical studies are shown in Figure 3 for the allele frequency datasets generated without missing values (*ϕ* = 0 in Eqs. (11) and (12)) and Figure 4 for the allele frequency datasets generated with missing values (*ϕ* = 0.02 in Eqs. (11) and (12)), respectively. In the two figures, the tips of the whiskers denote the 2.5%-quantile and the 97.5%-quantile, and the boxes represent the first and third quartile with the median in the middle. As can be seen from the boxplot results, our estimates for the selection coefficients at both loci show little bias across different parameter ranges, no matter whether sampled chromosomes contain unknown alleles or not, although one can discern a slight bias for small selection coefficients. With the increase of the selection coefficients, our estimates for the selection coefficients become more accurate. See the bias and the RMSE of the resulting MMSE estimates in Supplemental Material, Tables S1 and S2.

**Figure 3:**
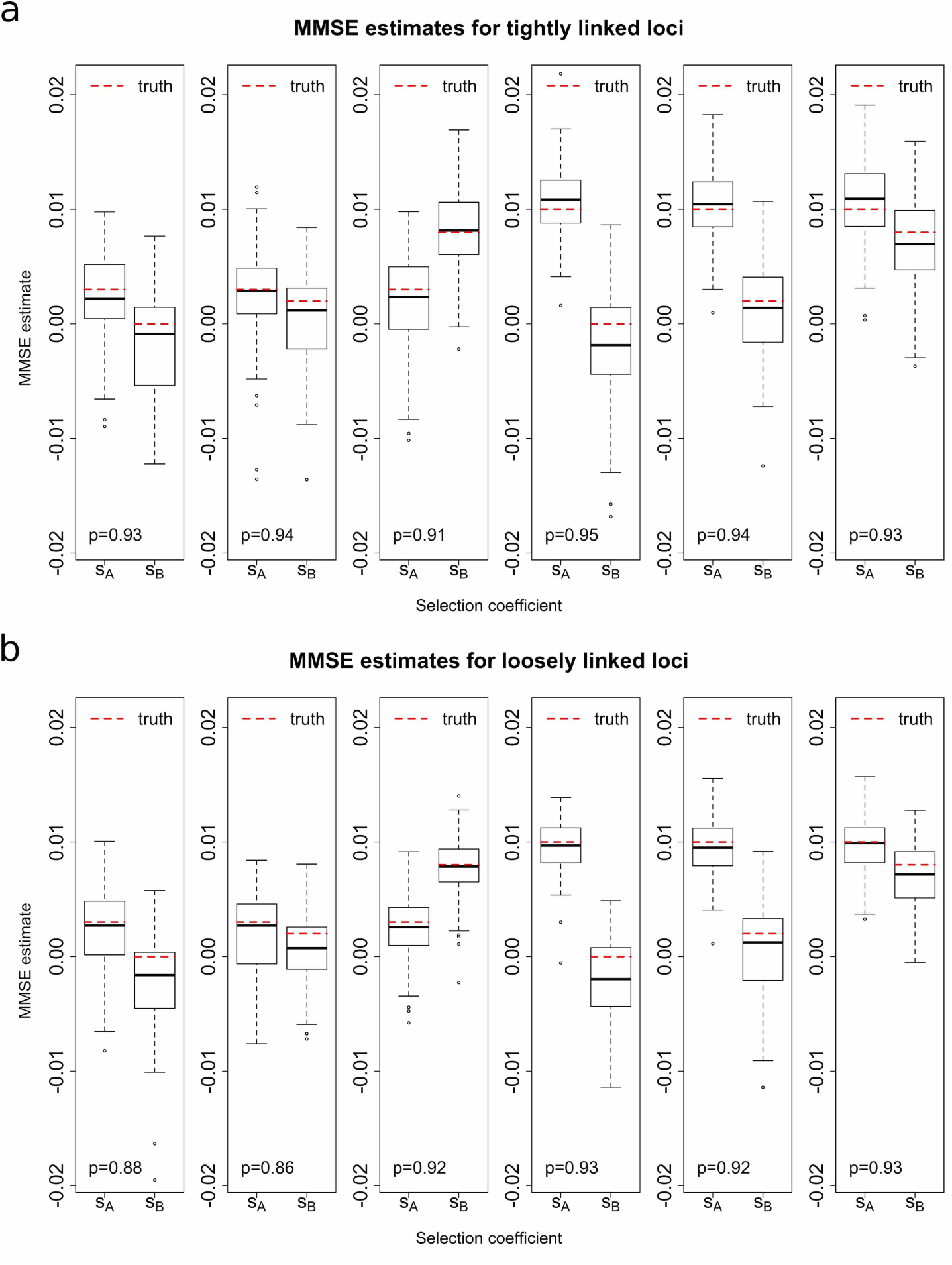
Empirical distributions of the MMSE estimates for 100 *allele frequency* datasets (*without* missing values) simulated with the initial population haplotype frequencies ***x***_0_ = (0.04, 0.08, 0.08, 0.8) and the dominance parameters *h*_𝒜_ = 0.5 and *h*_ℬ_ = 0.5 for the case of (a) tightly linked loci with the recombination rate *r* = 0.00001 and (b) loosely linked loci with the recombination rate *r* = 0.01. The *p* value in the bottom left corner indicates the proportion of the runs where the true values of the selection coefficients both fall within their 95% HPD intervals.

**Figure 4:**
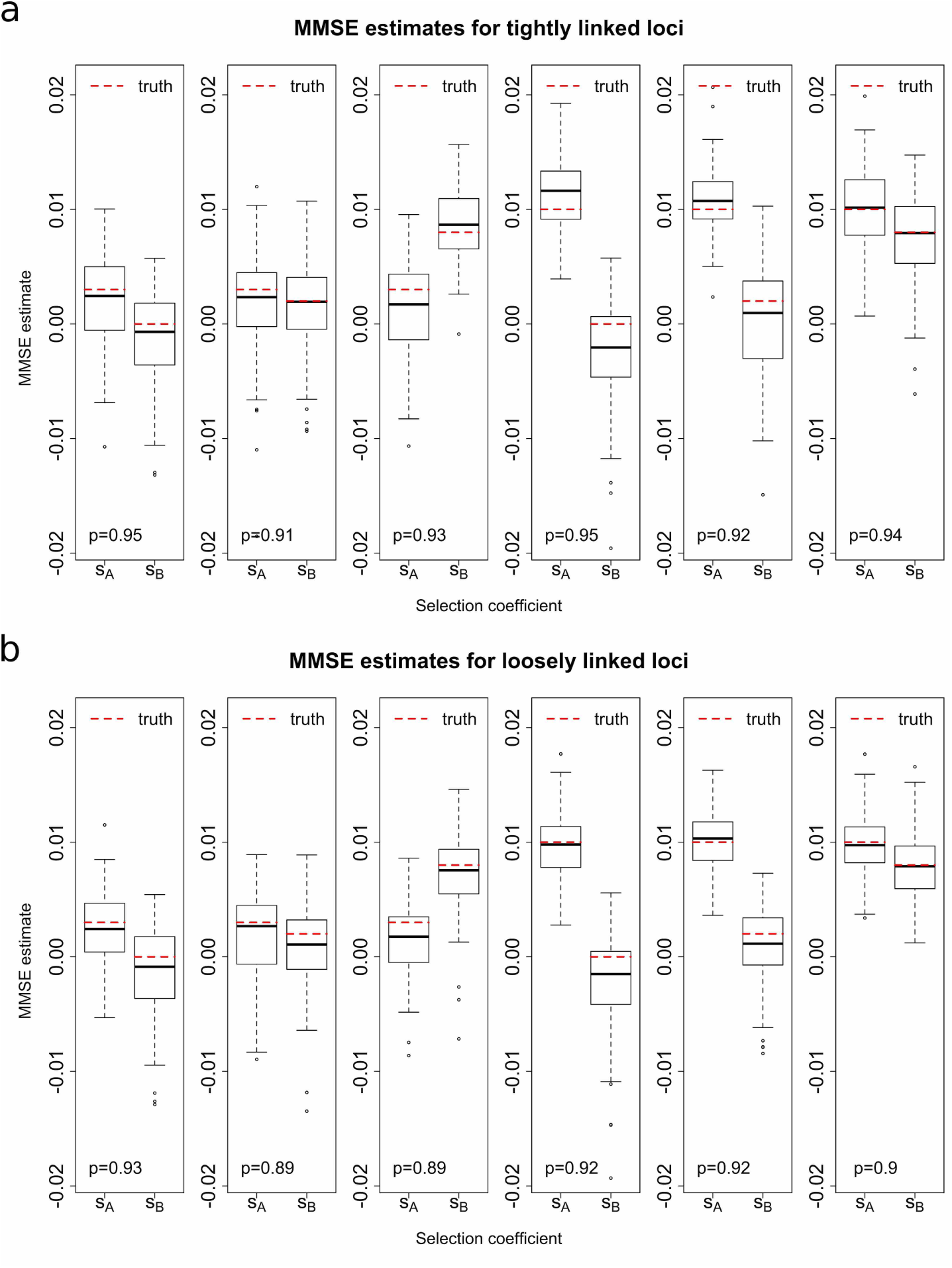
Empirical distributions of the MMSE estimates for 100 *allele frequency* datasets (*with* 2% missing values) simulated with the initial population haplotype frequencies ***x***_0_ = (0.04, 0.08, 0.08, 0.8) and the dominance parameters *h*_𝒜_ = 0.5 and *h*_ℬ_ = 0.5 for the case of (a) tightly linked loci with the recombination rate *r* = 0.00001 and (b) loosely linked loci with the recombination rate *r* = 0.01. The *p* value in the bottom left corner indicates the proportion of the runs where the true values of the selection coefficients both fall within their 95% HPD intervals.

For each combination of the selection coefficients and the recombination rate, we calculate the proportion of the 95% highest posterior density (HPD) intervals that include the true values, shown in the bottom left corner of each boxplot in Figures 3 and 4. On average, 92.00% of the runs result in the true values of the selection coefficients being within the 95% HPD intervals for the simulated datasets without missing values, 93.33% for tightly linked loci and 90.67% for loosely linked loci. For simulated datasets with 2% missing values, 92.08% of the runs result in the true values of the selection coefficients being within the 95% HPD intervals, *i.e*., 93.33% for tightly linked loci and 90.83% for loosely linked loci. We can see that small recombination rates can lead to better results for both loci.

In Figure 5, we illustrate the resulting boxplots of the empirical studies where the simulated data is given as haplotype frequencies (instead of allele frequencies in Figures 3 and 4). Compared to the estimates from allele frequency data, our estimates from haplotype frequency data are closer to their true values with smaller variances, especially for tightly linked loci. On average, 92.50% of the runs result in the true values of the selection coefficients being within their 95% HPD intervals on average, with 93.67% for tightly linked loci and 91.33% for loosely linked loci. The bias and the RMSE of the resulting MMSE estimates are summarised in Supplemental Material, Table S3. This improvement in the performance of our estimates is to be expected as all else being equal haplotype frequency data contain more information than allele frequency data. The complex interplay between the four haplotypes in the sample can be directly observed in haplotype frequency data but only partially observed in allele frequency data.

**Figure 5:**
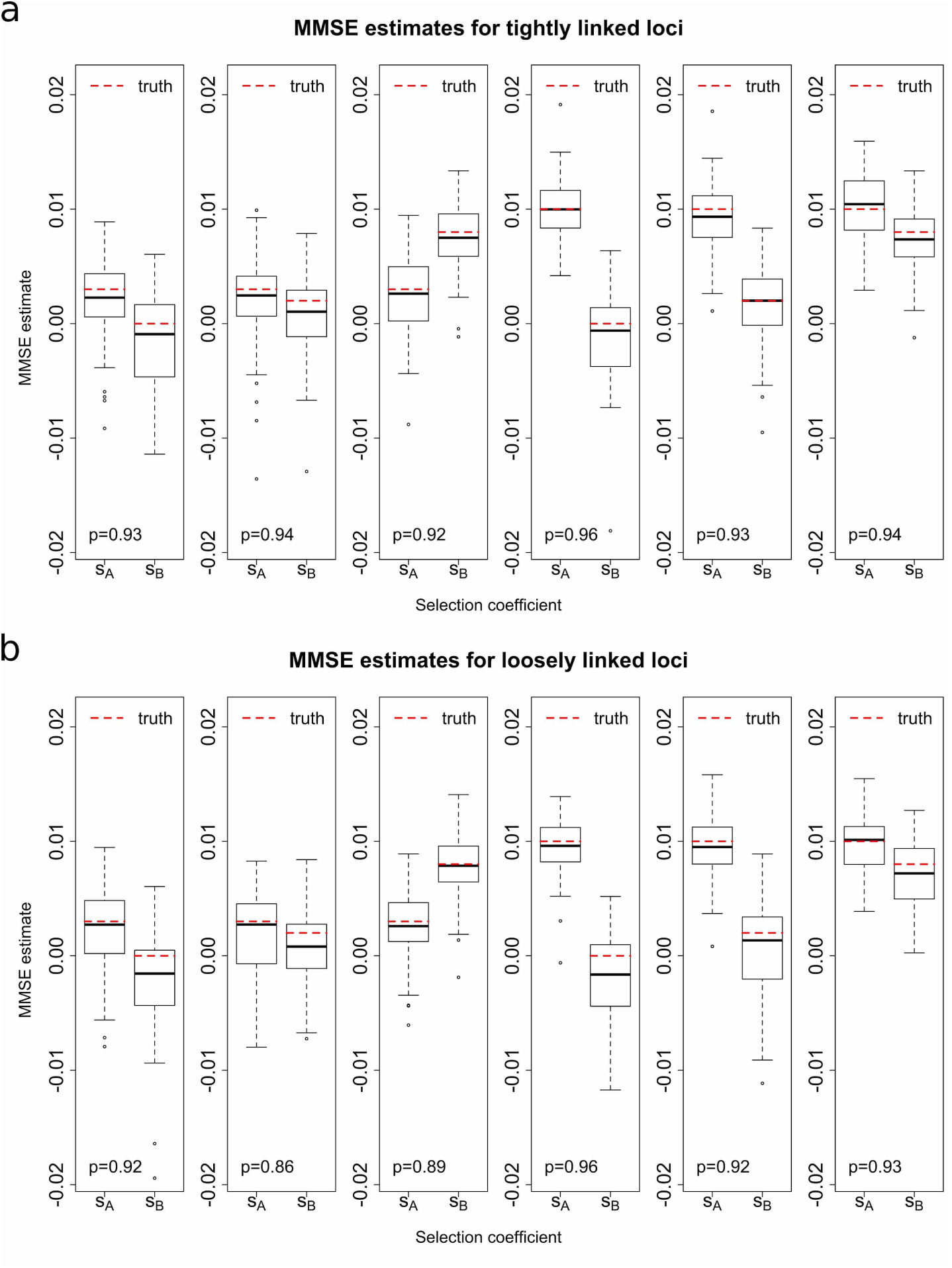
Empirical distributions of the MMSE estimates for 100 *haplotype frequency* datasets simulated with the initial population haplotype frequencies ***x***_0_ = (0.04, 0.08, 0.08, 0.8) and the dominance parameters *h*_𝒜_ = 0.5 and *h*_ℬ_ = 0.5 for the case of (a) tightly linked loci with the recombination rate *r* = 0.00001 and (b) loosely linked loci with the recombination rate *r* = 0.01. The *p* value in the bottom left corner indicates the proportion of the runs where the true values of the selection coefficients both fall within their 95% HPD intervals.

However, as illustrated in Figure 5, our estimates are still slightly biased for small selection coefficients. This may be caused by the initial population frequencies of the haplotypes that contain mutant alleles being close to 0 in our simulation studies. In this case, the population frequency trajectories of these haplotypes will be, with high probability, near 0 during the sampling period for small selection coefficients (see Supplemental Material, Figures S1 and S2).

This can cause a number of simulated datasets to have sample counts 0 for the haplotypes that contain mutant alleles, especially when the selection coefficients are small. Such datasets contain little information on the underlying selection coefficients. As can be observed from Figure 6, the bias can be almost completely eliminated for all combinations of the selection coefficients and the recombination rate if the starting population frequencies of the haplotypes that contain mutant alleles are taken to be intermediate values like ***x***_0_ = (0.1, 0.2, 0.3, 0.4). The bias and the RMSE of the resulting MMSE estimates are summarised in Supplemental Material, Table S4. The haplotype frequency trajectories of the underlying population for the haplotype frequency datasets simulated with the initial population haplotype frequencies ***x***_0_ = (0.1, 0.2, 0.3, 0.4) can be found in Supplemental Material, Figures S3 and S4). We also assess the performance of our method for the case that a new mutation arose in the population (at frequency 1*/*(2*N*)) at *t* = 0 when the neighbouring mutation became established. See Supplemental Material, Figure S5 and Table S5 for boxplots of the resulting MMSE estimates with their bias and RMSE, which show that our approach can still produce precise estimates of the selection coefficients in this case. It should be noticed that in this case we condition the mutant alleles at both loci to survive until the most recent sampling time point and sample 50 chromosomes from the underlying population at every 120 generations throughout 1200 generations so that a significant number of the realisations of the haplotype frequency trajectories of the underlying population can capture a significant proportion of the selective sweep.

**Figure 6:**
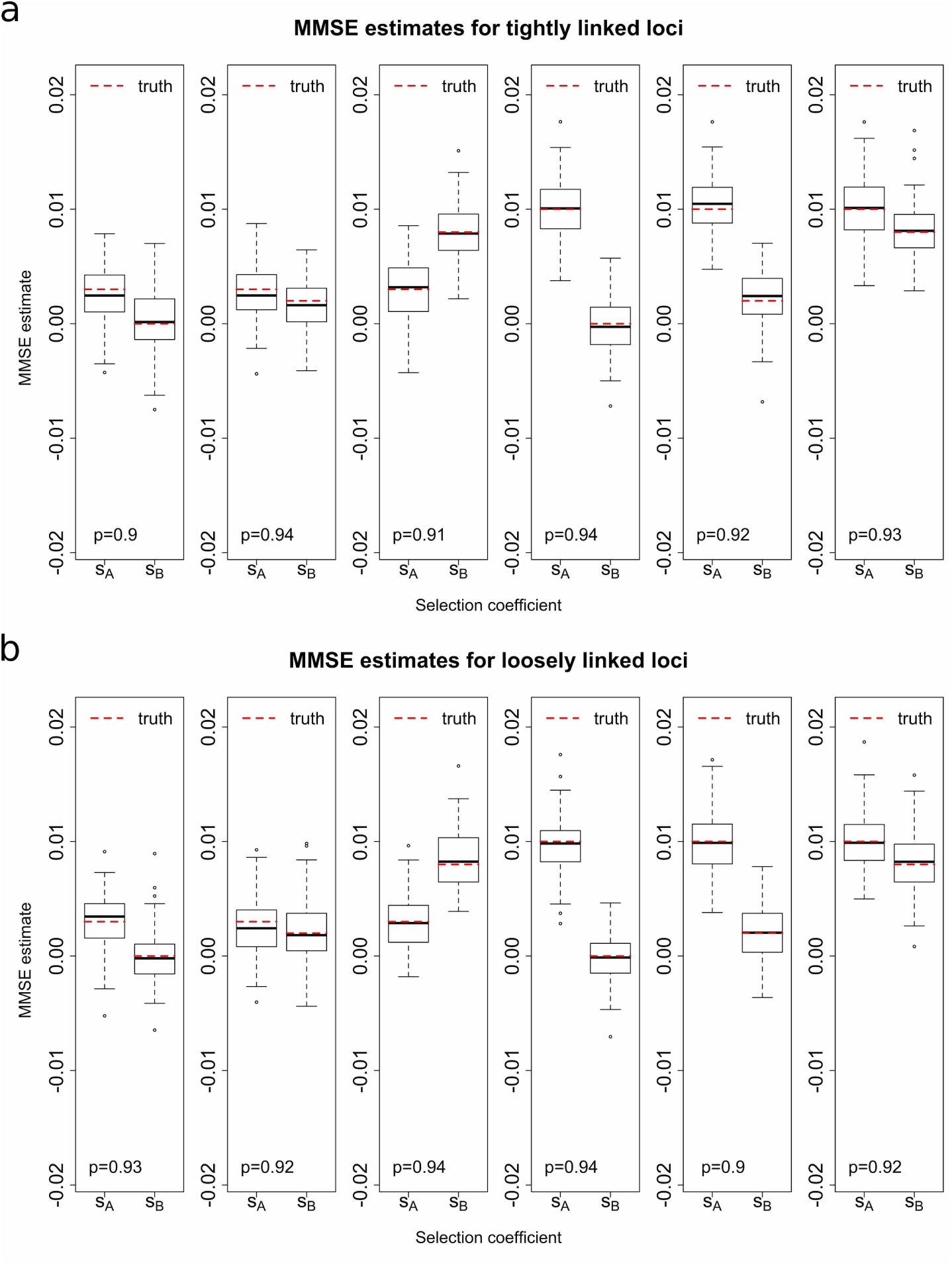
Empirical distributions of the MMSE estimates for 100 *haplotype frequency* datasets simulated with the initial population haplotype frequencies ***x***_0_ = (0.1, 0.2, 0.3, 0.4) and the dominance parameters *h*_𝒜_ = 0.5 and *h*_ℬ_ = 0.5 for the case of (a) tightly linked loci with the recombination rate *r* = 0.00001 and (b) loosely linked loci with the recombination rate *r* = 0.01. The *p* value in the bottom left corner indicates the proportion of the runs where the true values of the selection coefficients both fall within their 95% HPD intervals.

In summary, our Bayesian inference procedure can deliver accurate estimates of the selection coefficients using time series data of allele frequencies across different parameter ranges, regardless of whether sampled chromosomes contain unknown alleles or not. We also simulated datasets with other selection schemes, *e.g*., the dominance parameters *h*_𝒜_ = 0 and *h*_ℬ_ = 1. The resulting boxplots of the simulation studies are shown in Supplemental Material, Figure S6, with the bias and the RMSE of the resulting MMSE estimates summarised in Supplemental Material, Table S6. In addition to MMSE estimates, we also presented the bias and the RMSE of MAP estimates (see Supplemental Material, Tables S7-S12), which display very similar characteristics to the MMSE estimates. The boxplots for MAP estimates show little bias, with the upper and lower quartiles of the MAP estimates being similar to those of the MMSE estimates (see Supplemental Material, Figures S7-S12).

#### 3.1.2. Improvement from modelling genetic recombination and local linkage

In the case where a pair of loci are both suspected to be subject to natural selection, one can still apply a single-locus approach to each locus to estimate selection coefficient. To our knowledge, there has been a considerable amount of work on the statistical inference of natural selection at a single locus from time series data of allele frequencies (*e.g*., Bollback et al., 2008; Malaspinas et al., 2012; Steinrücken et al., 2014; Schraiber et al., 2016; Ferrer-Admetlla et al., 2016; He et al., 2019). However, using a single-locus approach may lead to inaccurate estimates of the selection coefficients when the two loci are linked (He et al., 2020). In the case of tightly linked loci, modelling genetic recombination and local linkage becomes necessary, thus our two-locus method is far more desirable. Below we illustrate with two examples of tightly linked loci with the recombination rate *r* = 0.00001. We simulate the haplotype frequency trajectories of the underlying population through the two-locus Wright-Fisher model with selection and draw 200 chromosomes from the underlying population at generations 0, 100, 200, 300, 400 and 500. In the first example, we consider a positively selected locus 𝒜 tightly linked with a selectively neutral locus ℬ, where we set the selection coefficients *s*_𝒜_ = 0.01 and *s*_ℬ_ = 0, respectively. We take the initial haplotype frequencies of the underlying population to be ***x***_0_ = (0.2, 0.1, 0.3, 0.4). The mutant allele frequency trajectories of the sample are shown in Figure 7a. The posterior probability distributions obtained with our single-locus approach, described in Supplemental Material, File S3, are shown in Figure 7b, and the posterior probability distributions achieved with our two-locus method, described in Section 2.2, are shown in Figure 7c. Bayesian estimates of the selection coefficients *s*_𝒜_ and *s*_ℬ_ are summarised in Table 1.

**Table 1:** A comparison of the Bayesian estimates obtained by using the single-locus method and the two-locus method from the simulated dataset of a positively selected locus tightly linked with a selectively neutral locus.

|  |  | single-locus method | two-locus method |
| --- | --- | --- | --- |
| selection coefficient $s_{\mathcal{A}}$ | MAP ($\times 10^{-2}$) | 1.160 | 0.989 |
| | MMSE ($\times 10^{-2}$) | 1.127 | 1.148 |
| | 95% HPD ($\times 10^{-2}$) | [0.722, 1.566] | [0.652, 1.630] |
| selection coefficient $s_{\mathcal{B}}$ | MAP ($\times 10^{-2}$) | 0.465 | 0.214 |
| | MMSE ($\times 10^{-2}$) | 0.496 | 0.253 |
| | 95% HPD ($\times 10^{-2}$) | [0.080, 0.916] | [-0.495, 1.123] |

**Figure 7:**
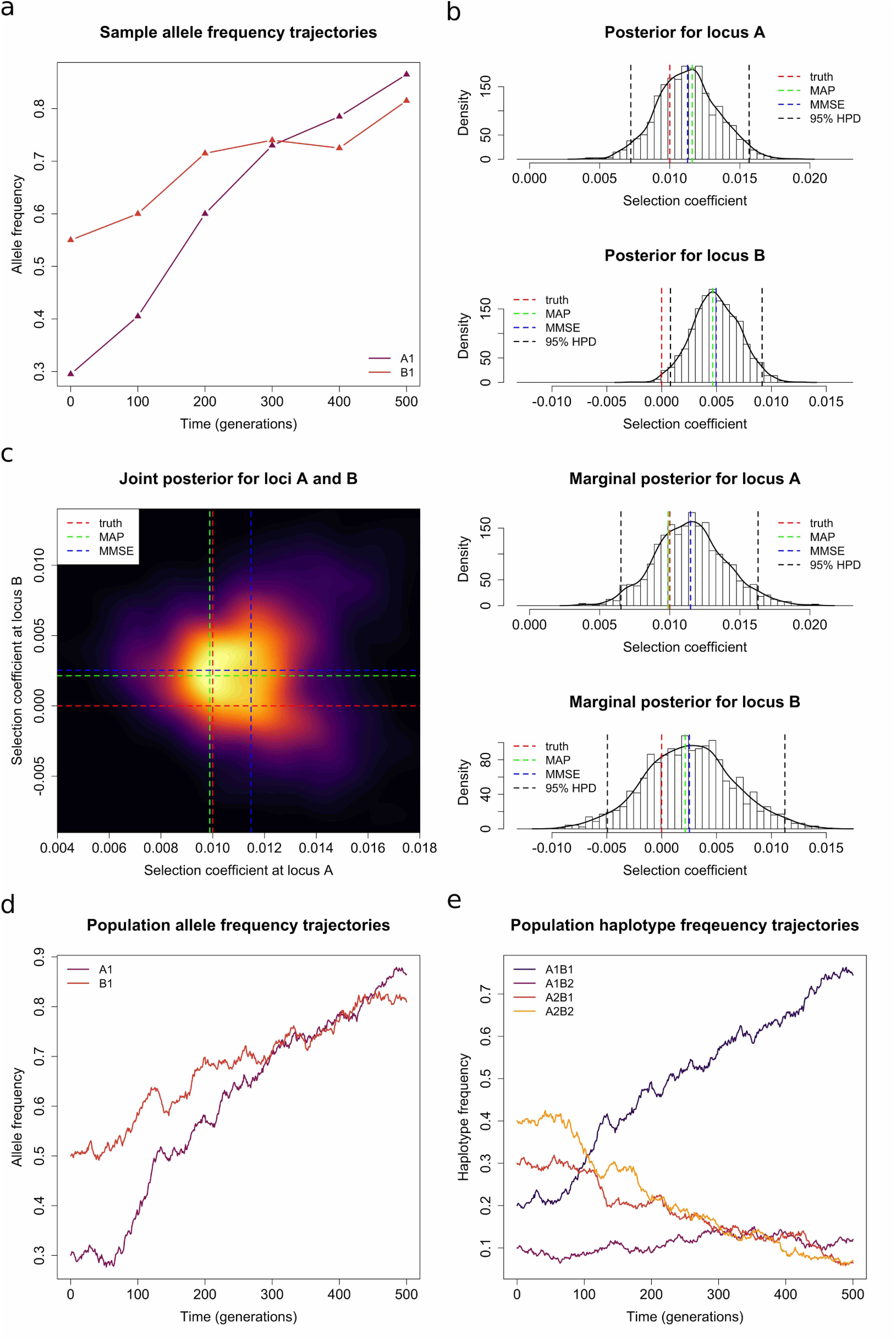
A comparison of the performance differences of the single-locus method and the two-locus method on the simulated dataset of a positively selected locus tightly linked with a selectively neutral locus. (a) Sample mutant allele frequency trajectories. (b) Posteriors obtained with a single-locus method. (c) Posteriors obtained with a two-locus method. (d) Population mutant allele frequency trajectories. (e) Population haplotype frequency trajectories.

One can observe that with a single-locus method, the estimate for the selection coefficient *s*_𝒜_ is reasonably accurate, but the estimate for the selection coefficient *s*_ℬ_ is off by a large amount. The true value for the selection coefficient *s*_ℬ_ is 0, but the single-locus approach produces an estimate of roughly 0.005 and a 95% HPD interval that only encompasses positive values, which is strong evidence for the presence of positive selection. In comparison, the estimates for both of the selection coefficients *s*_𝒜_ and *s*_ℬ_ are fairly accurate with the two-locus method.

To understand the poor performance of the single-locus method in this example, we plot the mutant allele frequency trajectories of the underlying population in Figure 7d and the haplotype frequency trajectories of the underlying population in Figure 7e. The increase in the frequency of the ℬ_1_ allele with time, despite it having a selection coefficient of 0, is caused by the 𝒜_1_ℬ_1_ haplotype, which has a selection coefficient of 0.01. This compensates for the decrease in the frequency of the 𝒜_2_ℬ_1_ haplotype, resulting in an increasing trajectory for the ℬ_1_ allele, albeit with a slower rate of increase than the 𝒜_1_ allele. With the two-locus approach, however, the interplay between all four haplotypes are taken into account and it produces accurate estimates for both of the selection coefficients *s*_𝒜_ and *s*_ℬ_.

In the second example, we consider two positively selected and tightly linked loci 𝒜 and ℬ, where we take the selection coefficients to be *s*_𝒜_ = 0.01 and *s*_ℬ_ = 0.005, respectively, and set the initial haplotype frequencies of the underlying population to be ***x***_0_ = (0.05, 0.05, 0.7, 0.2). The results are illustrated in Figure 8 and summarised in Table 2. In this example, with the single-locus method, the estimate for the selection coefficient *s*_𝒜_ is reasonably accurate, but the estimate for the selection coefficient *s*_ℬ_ is off by a large amount, *i.e*., its true value lies outside the 95% HPD interval. In fact, although the ℬ_1_ allele is favoured by natural selection with a selection coefficient of 0.005, the resulting estimate for the selection coefficient *s*_ℬ_ is roughly −0.0015 with a 95% HPD interval that includes the value 0, which implies no strong evidence for natural selection. In comparison, the two-locus method again produces fairly accurate estimates for both of the selection coefficients *s*_𝒜_ and *s*_ℬ_.

**Table 2:** A comparison of the Bayesian estimates obtained by using the single-locus method and the two-locus method from the simulated dataset of a pair of positively selected and tightly linked loci.

|  |  | single-locus method | two-locus method |
| --- | --- | --- | --- |
| selection coefficient $s_{\mathcal{A}}$ | MAP ($\times 10^{-2}$) | 0.811 | 0.879 |
| | MMSE ($\times 10^{-2}$) | 0.789 | 0.966 |
| | 95% HPD ($\times 10^{-2}$) | [0.338, 1.290] | [0.412, 1.673] |
| selection coefficient $s_{\mathcal{B}}$ | MAP ($\times 10^{-2}$) | -0.176 | 0.219 |
| | MMSE ($\times 10^{-2}$) | -0.148 | 0.267 |
| | 95% HPD ($\times 10^{-2}$) | [-0.560, 0.271] | [-0.433, 0.958] |

**Figure 8:**
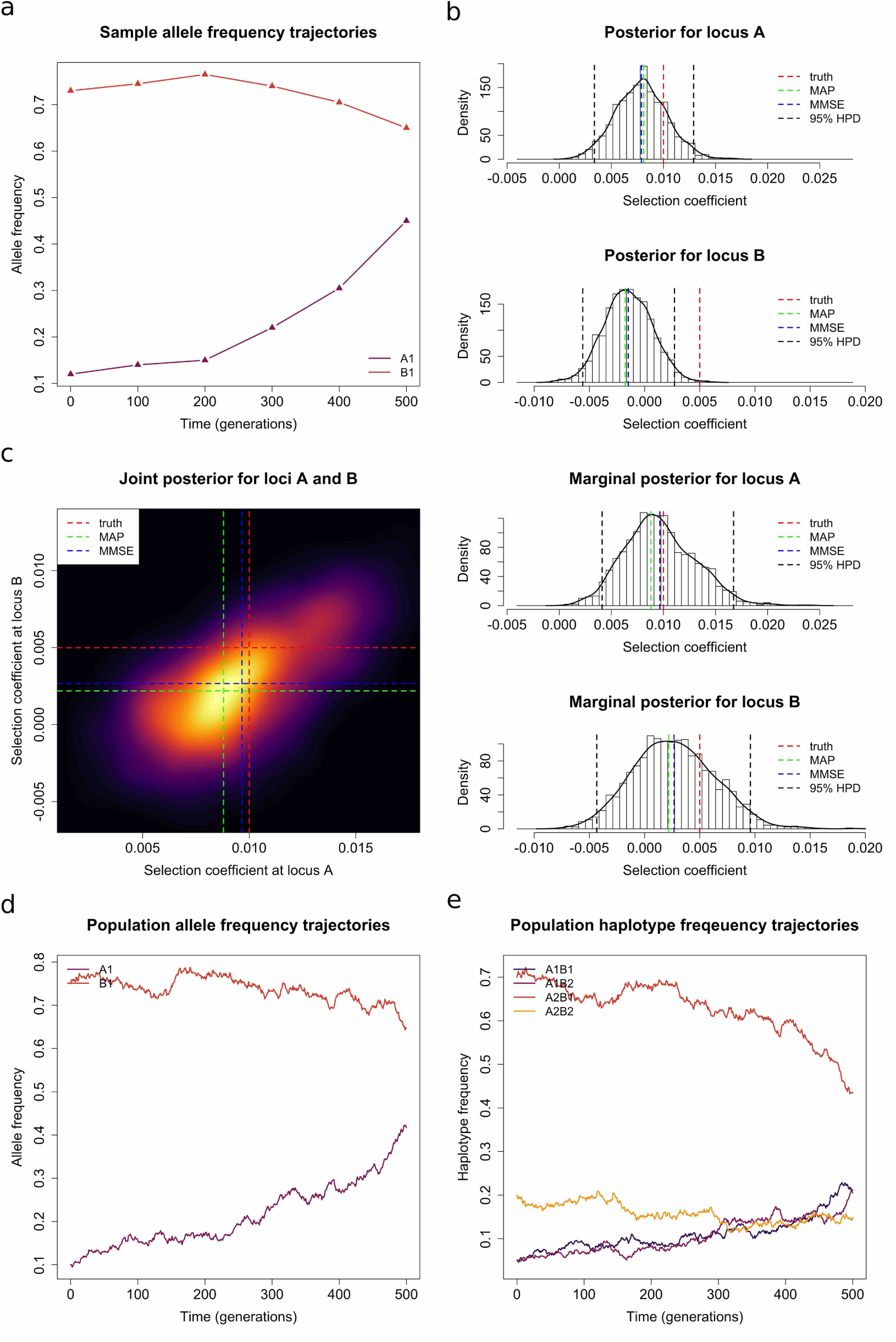
A comparison of the performance differences of the single-locus method and the two-locus method on the simulated dataset of a pair of positively selected and tightly linked loci. (a) Sample mutant allele frequency trajectories. (b) Posteriors obtained with a single-locus method. (c) Posteriors obtained with a two-locus method. (d) Population mutant allele frequency trajectories. (e) Population haplotype frequency trajectories.

As shown in Figure 8, the frequency of the 𝒜_1_ allele increases with time due to the increase in the frequencies of the 𝒜_1_ℬ_1_ and 𝒜_1_ℬ_2_ haplotypes, which are the two most selected haplotypes, with the selection coefficients of 0.015 and 0.01, respectively. The ℬ_1_ allele is made up of the 𝒜_1_ℬ_1_ and 𝒜_2_ℬ_1_ haplotypes, with the selection coefficients of 0.01 and 0.005, respectively, which are the second and third most selected haplotypes. As a result of their initial conditions and selection coefficients, the frequency of the ℬ_1_ allele roughly holds constant in time, since it is somewhat out-competed by the 𝒜_1_ allele. Viewing the trajectory of the ℬ_1_ allele in isolation does not give strong evidence that it is selectively advantageous, which results in an estimate of roughly 0 in its selection coefficient through the single-locus approach. Moreover, even the 95% HPD interval for the single-locus method does not include the true selection coefficient of 0.005 for the ℬ_1_ allele. Using the two-locus approach, we are again able to obtain accurate estimates for both of the selection coefficients *s*_𝒜_ and *s*_ℬ_.

In these two examples, we choose a uniform prior over the interval [−1, 1] for the selection coefficients, and we select a flat Dirichlet prior for the initial population haplotype frequencies in the two-locus method and a uniform prior over the interval [0, 1] for the initial population allele frequency in the single-locus method, respectively. Other settings in the Euler-Maruyama scheme and the PMMH algorithm are the same as we applied in the empirical studies in Section 3.1.1. Compared to existing single-locus approaches, our two-locus method explicitly incorporates the effect of genetic recombination and the information of local linkage through the two-locus Wright-Fisher diffusion with selection. Indeed, the dynamics of the two-locus Wright-Fisher diffusion with selection can demonstrate complex and unpredictable behaviour (see, *e.g*., Yu & Etheridge, 2010; Cuthbertson et al., 2012), which can result in inaccurate estimates of the selection coefficients if one simply employs a single-locus approach. In contrast, applying our two-locus method can yield precise estimates of the selection coefficients at both loci.

### 3.2. Analysis of real data

We apply our Bayesian inference method to real data by re-analysing time serial samples of segregating alleles of the equine homologue of proto-oncogene c-kit (*KIT*). These data come from previous studies of Ludwig et al. (2009), Pruvost et al. (2011) and Wutke et al. (2016), and the sample information and genotyping results for all successfully typed horses can be found in Wutke et al. (2016), which are summarised in Table 3. The *KIT* gene in horses resides on the long arm of chromosome 3 and lies in two intervals associated with white spotting patterns, one in the intron 13 which codes for tobiano (*KIT13*), with the other in intron 16 which codes for sabino (*KIT16*). At the *KIT13* locus, the ancestral allele is designated *KM0*, while the mutant allele, associated with the tobiano pattern and acting as dominant (Brooks et al., 2007), is designated *KM1*. The tobiano pattern is characterised by depigmented patches of skin and associated hair that often cross the dorsal midline and cover the legs. At the *KIT16* locus, the ancestral allele is designated *sb1*, while the mutant allele associated with the sabino pattern and acting as semi-dominant (Brooks & Bailey, 2005), is designated *SB1*. The sabino pattern is characterised by irregularly bordered white patches of skin and associated hair that begin at the extremities and face and may extend up to the belly and midsection.

**Table 3:**
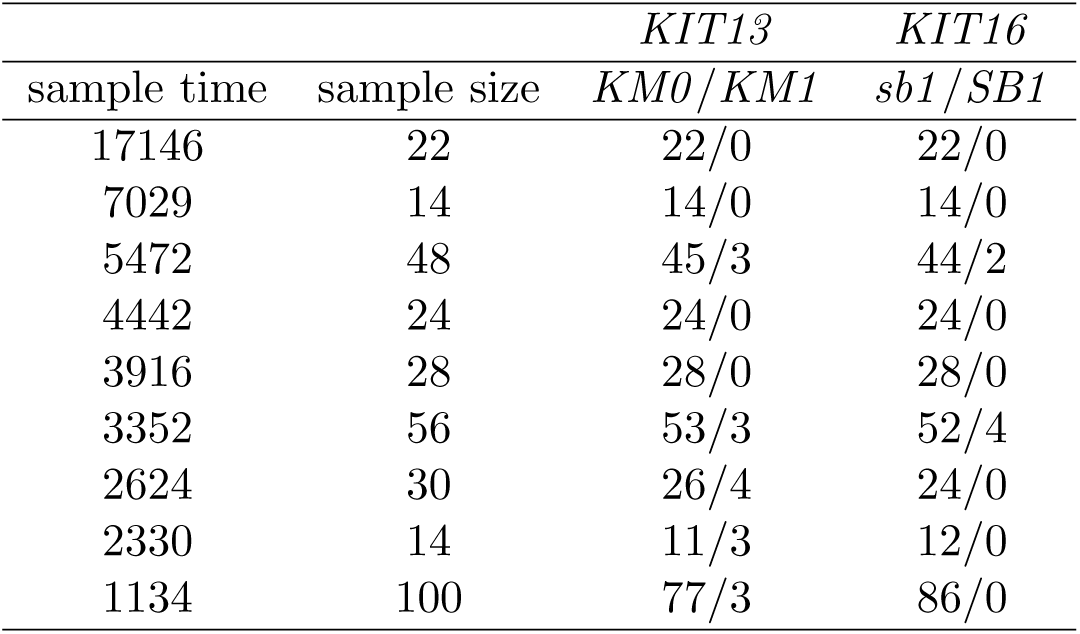
Time serial ancient horse samples of segregating alleles at the *KIT13* and *KIT16* loci. The unit of the sampling time is the year before present (BP)

|  |  | *KIT13* | *KIT16* |
| --- | --- | --- | --- |
| sample time | sample size | *KM0/KM1* | *sb1/SB1* |
| 17146 | 22 | 22/0 | 22/0 |
| 7029 | 14 | 14/0 | 14/0 |
| 5472 | 48 | 45/3 | 44/2 |
| 4442 | 24 | 24/0 | 24/0 |
| 3916 | 28 | 28/0 | 28/0 |
| 3352 | 56 | 53/3 | 52/4 |
| 2624 | 30 | 26/4 | 24/0 |
| 2330 | 14 | 11/3 | 12/0 |
| 1134 | 100 | 77/3 | 86/0 |

We set the dominance parameters *h* = 0 for *KIT13* as the *KM1* allele is dominant, and *h* = 0.5 for *KIT16* as the *SB1* allele is semi-dominant. Following Der Sarkissian et al. (2015), we take the population size to be *N* = 16000 and the average length of a generation of horse to be 8 years, the same as in Schraiber et al. (2016). As can be seen in Table 3, there are various sampling time points when the sequencing of the aDNA material yielded a number of unknown alleles at loci *KIT13* and/or *KIT16*. We show all possible mutant allele frequency trajectories of the sample at the *KIT13* and *KIT16* loci in Figure 9. Neither mutant allele was found in the first two samples dated 17146 and 7029 years BP. Indeed, both sabino and tobiano patterns are only present in domestic horses (Wutke et al., 2016). We assume that both mutant alleles, *KM1* and *SB1*, arose after the domestication of the horse, which is thought to have started in the Eurasian Steppes around 5500 years BP (Outram et al., 2009). We therefore discard the first two samples from our analysis in this section, but for completeness, in Supplemental Material, Figures S13-S18, we also present the results of the inference when these two samples are taken into account.

**Figure 9:**
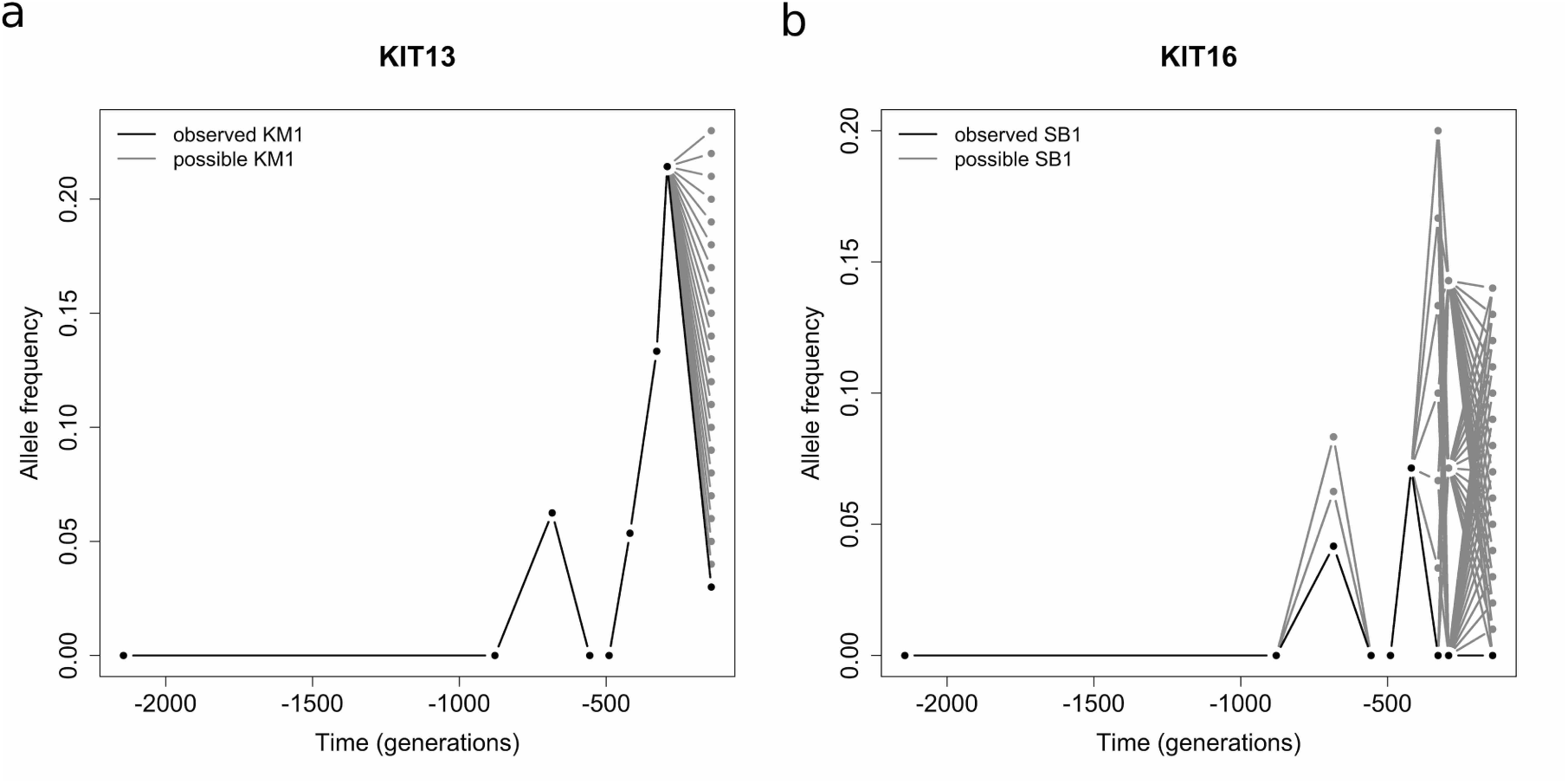
Potential changes in the mutant allele frequencies of the sample over time at loci (a) *KIT13* and (b) *KIT16*. Ancient horse samples were taken at generations -2144, -879, -684, -556, -490, -419, -328, -292 and -142.

As a result of the low quality of the *KIT* dataset, it becomes difficult to intuit whether either or both mutant alleles at the *KIT13* and *KIT16* loci are selected by simply inspecting the mutant allele frequency trajectories of the sample. Using our two-locus Bayesian inference procedure, described in Section 2.2, we jointly estimate the selection coefficients for the mutant alleles at the *KIT13* and *KIT16* loci under the case that sampled chromosomes contain variants with unknown alleles. For the recombination rate, we choose three average rates of recombination, 5 × 10^−9^, 1 × 10^−8^ and 5 × 10^−8^ crossovers/bp, as suggested in Dumont & Payseur (2008), and multiply them by the genetic distance 4688 bp to get the recombination rates between the *KIT13* and *KIT16* loci. All settings in the Euler-Maruyama scheme and the PMMH algorithm are the same as we applied in the previous section. The resulting posterior probability distributions are shown in Figure 10, and the MAP and MMSE estimates, as well as the 95% HPD intervals, are summarised in Table 4.

**Table 4:**
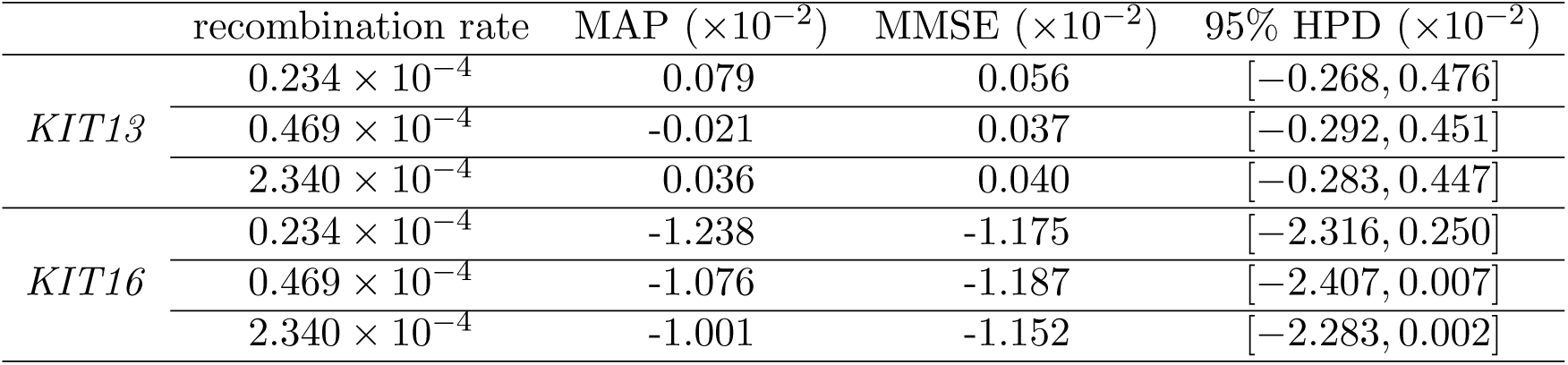
MAP and MMSE estimates, as well as the 95% HPD intervals, for *KIT13* and *KIT16* obtained by using the two-locus method from the samples dated from 5472 years BP (the third sampling time point) with the population size of 16000.

**Figure 10:**
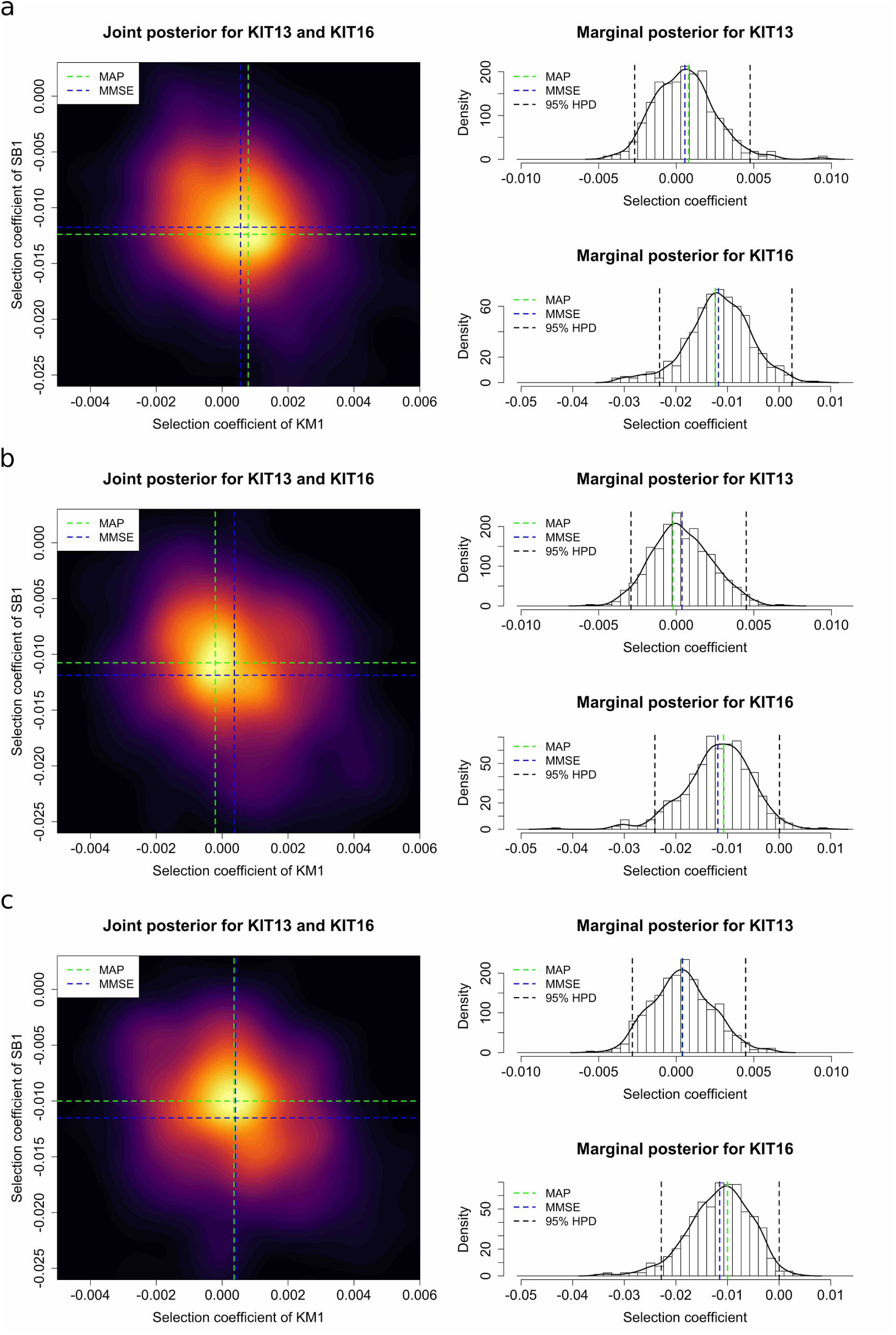
Posterior probability distributions for *KIT13* and *KIT16* obtained by using the two-locus method from the samples dated from 5472 years BP (the third sampling time point) with the population size of 16000 and the average rate of recombination (a) 5 × 10^−9^ crossovers/bp, (b) 1 × 10^−8^ crossovers/bp and (c) 5 × 10^−8^ crossovers/bp.

As can be found in Table 4, the MMSE estimates with different values of the recombination rate are essentially unchanged, while the MAP estimates vary a bit more than the MMSE estimates. This may be caused by the way we achieve our MAP estimates, where the posterior probability distribution is approximated through the two-dimensional kernel density estimation with an axis-aligned bivariate normal kernel (Venables & Ripley, 2002). Therefore, the MAP estimates may depend on the number of the iterations of the PMMH. The resulting Bayesian estimates of the selection coefficients suggest that the *KM1* allele at the *KIT13* locus is weakly positively selected whereas the *SB1* allele at the *KIT16* locus is strongly negatively selected, but the 95% HPD intervals for both selection coefficients include the value 0. For the *KIT13* locus, the posterior probability for positive selection is 0.564, not strong evidence for the *KM1* allele at the *KIT13* locus being positively selected. However, for the *KIT16* locus, the posterior probability for negative selection is 0.982, strong evidence to support the *SB1* allele at the *KIT16* locus being negatively selected. This conclusion is further confirmed with the estimates obtained with different values of the population size (*i.e*., *N* = 8000 and *N* = 32000), which can be found in Supplemental Material, Figures S19 and S20.

We also used our single-locus Bayesian inference procedure, described in Supplemental Material, File S3, to independently estimate the selection coefficients for the mutant alleles at the *KIT13* and *KIT16* loci under the case that sampled chromosomes contain unknown alleles. All settings in the Euler-Maruyama scheme and the PMMH algorithm are the same as we applied in the previous section. The resulting posterior probability distributions are shown in Figure 11, and the MAP and MMSE estimates, as well as the 95% HPD intervals, are summarised in Table 5. The resulting Bayesian estimates of the selection coefficients suggest that the *KM1* allele at the *KIT13* locus is weakly selectively advantageous whereas the *SB1* allele at the *KIT16* locus is weakly selectively deleterious. However, as illustrated in Figure 5, the posterior probability distributions for the *KIT13* and *KIT16* loci are both roughly symmetric about 0. This indicates that there is no evidence to support the *KM1* allele at the *KIT13* locus or the *SB1* allele at the *KIT16* locus being selected, which is consistent with the findings of Ludwig et al. (2009) obtained by using the approach of Bollback et al. (2008). Compared to the results shown in Figure 10 and Table 4, we fail to tease apart negative selection at the *KIT16* locus without considering genetic recombination effect and local linkage information. We present an example that mimics the *KIT13* and *KIT16* loci, *i.e*., a negatively selected locus tightly linked with a selectively neutral locus, which shows similar results to those using the real dataset (see Supplemental Material, Figure S21 and Table S13).

**Table 5:**
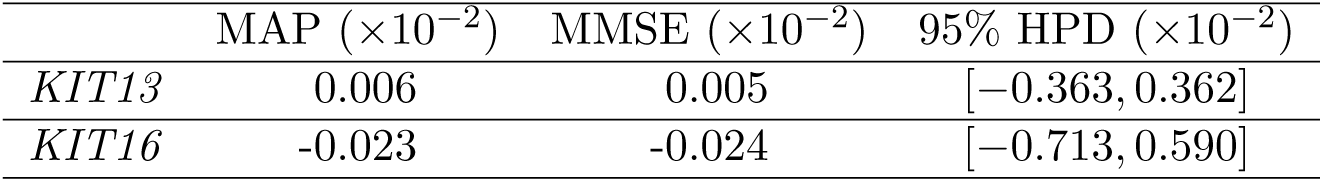
MAP and MMSE estimates, as well as the 95% HPD intervals, for *KIT13* and *KIT16* obtained by using the single-locus method from the samples dated from 5472 years BP (the third sampling time point) with the population size of 16000.

| | MAP ($\times 10^{-2}$) | MMSE ($\times 10^{-2}$) | 95% HPD ($\times 10^{-2}$) |
| --- | --- | --- | --- |
| *KIT13* | 0.006 | 0.005 | $[-0.363, 0.362]$ |
| *KIT16* | -0.023 | -0.024 | $[-0.713, 0.590]$ |

**Figure 11:**
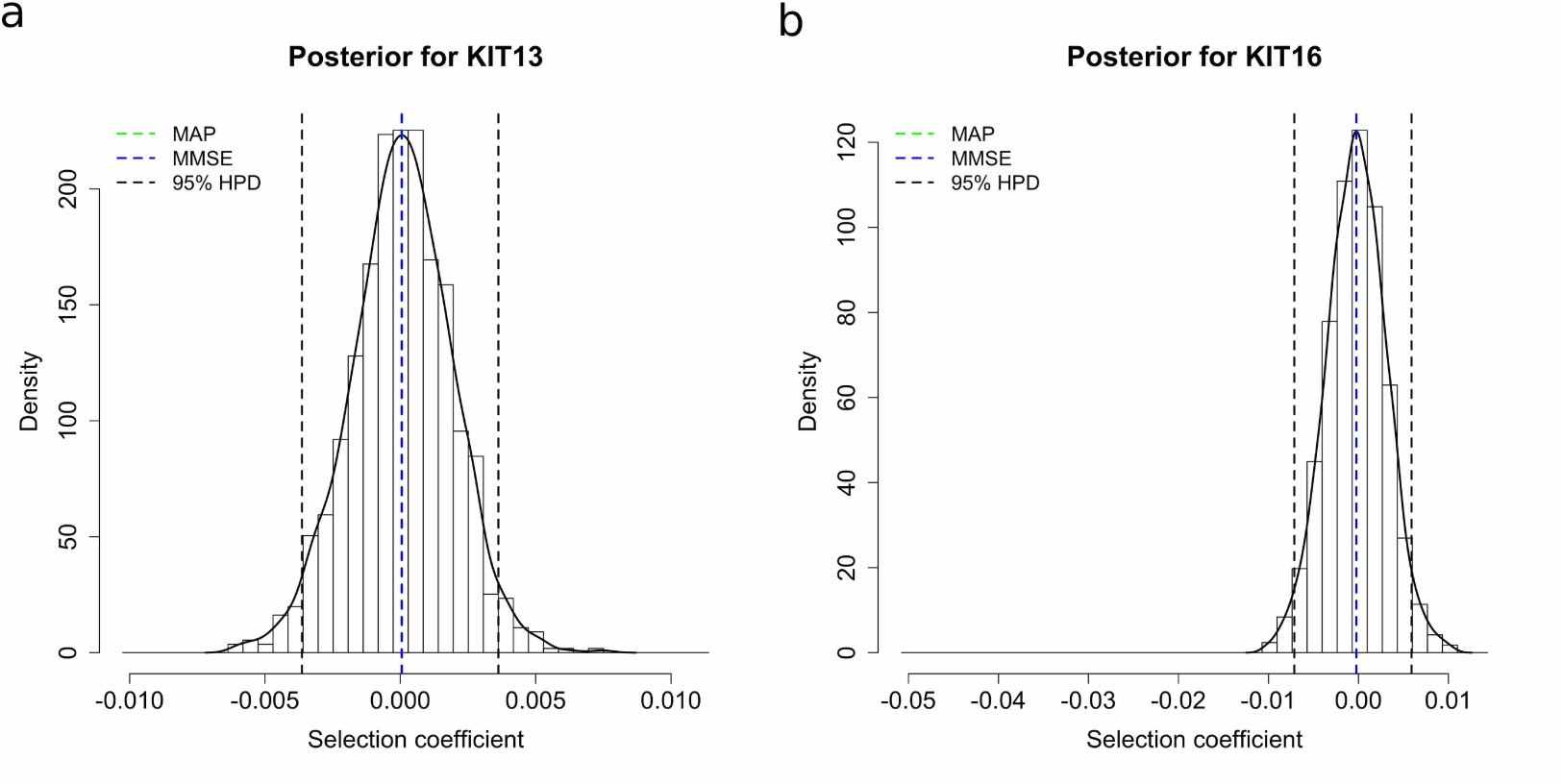
Posterior probability distributions for (a) *KIT13* and (b) *KIT16* obtained by using the single-locus method from the samples dated from 5472 years BP (the third sampling time point) with the population size of 16000.

### 3.3. Computational issues

In the PMMH algorithm, it is desirable to generate a large number of particles in the bootstrap particle filter to yield an accurate estimate of the marginal likelihood *p*(***u***_1:*K*_, ***v***_1:*K*_ | ***ϑ***). However, this can be computational burdensome since each iteration of the PMMH algorithm requires a run of the bootstrap particle filter even though fewer iterations are required. Balancing the particle number and the MCMC iteration number to obtain good performance at a reasonable computational cost was investigated by Pitt et al. (2012) and Doucet et al. (2015). In pseudo-marginal algorithms, if the estimates of the marginal likelihood are too noisy, the chain tends to be ‘sticky’ with excessive autocorrelation (Beaumont, 2003). A simple rule-of-thumb is to select a particle number such that the standard deviation of the log-likelihood estimates is in the range from 1.0 to 1.7. Nevertheless, the PMMH algorithm exactly targets the marginal posterior *p*(***ϑ*** | ***u***_1:*K*_, ***v***_1:*K*_) for any number of particles.

In each run of the bootstrap particle filter, we simulate the particles according to the two-locus Wright-Fisher diffusion with selection using the Euler-Maruyama scheme. It is desirable to take a large *L* in the Euler-Maruyama scheme to get an accurate approximation of the Wright-Fisher diffusion, but large *L* increases the computational load. Stramer & Bognar (2011) suggested choosing *L* to be *L*^*\**^ such that the estimates of the marginal likelihood are approximately the same for any *L* > *L*^*\**^, where *L*^*\**^ can be obtained through extensive simulations.

In practice, we divide each generation into 5 subintervals in the Euler-Maruyama scheme, *i.e*., *L* = 5. Our running time for a single iteration of the PMMH algorithm with 1500 particles (see Figure 12), achieving the standard deviation of the log-likelihood at approximately 1.504, on a single core of an Intel Core i7 processor at 4.2 GHz, is around 12.360 seconds for the *KIT* dataset. In principle, every particle can be simulated in parallel on a different core. Running 10000 iterations of the PMMH is sufficient for a relatively smooth resulting posterior surface, as shown in Figure 10. We discard the initial 2000 iterations as the burn-in period and then thin the remaining PMMH output, taking every fourth value and regarding these as independent. Dahlin & Schön (2015) outlined a selected number of possible improvements and best practices for implementation. All of our code in this work is written in R with C++ by using Rcpp and RcppArmadillo.

**Figure 12:**
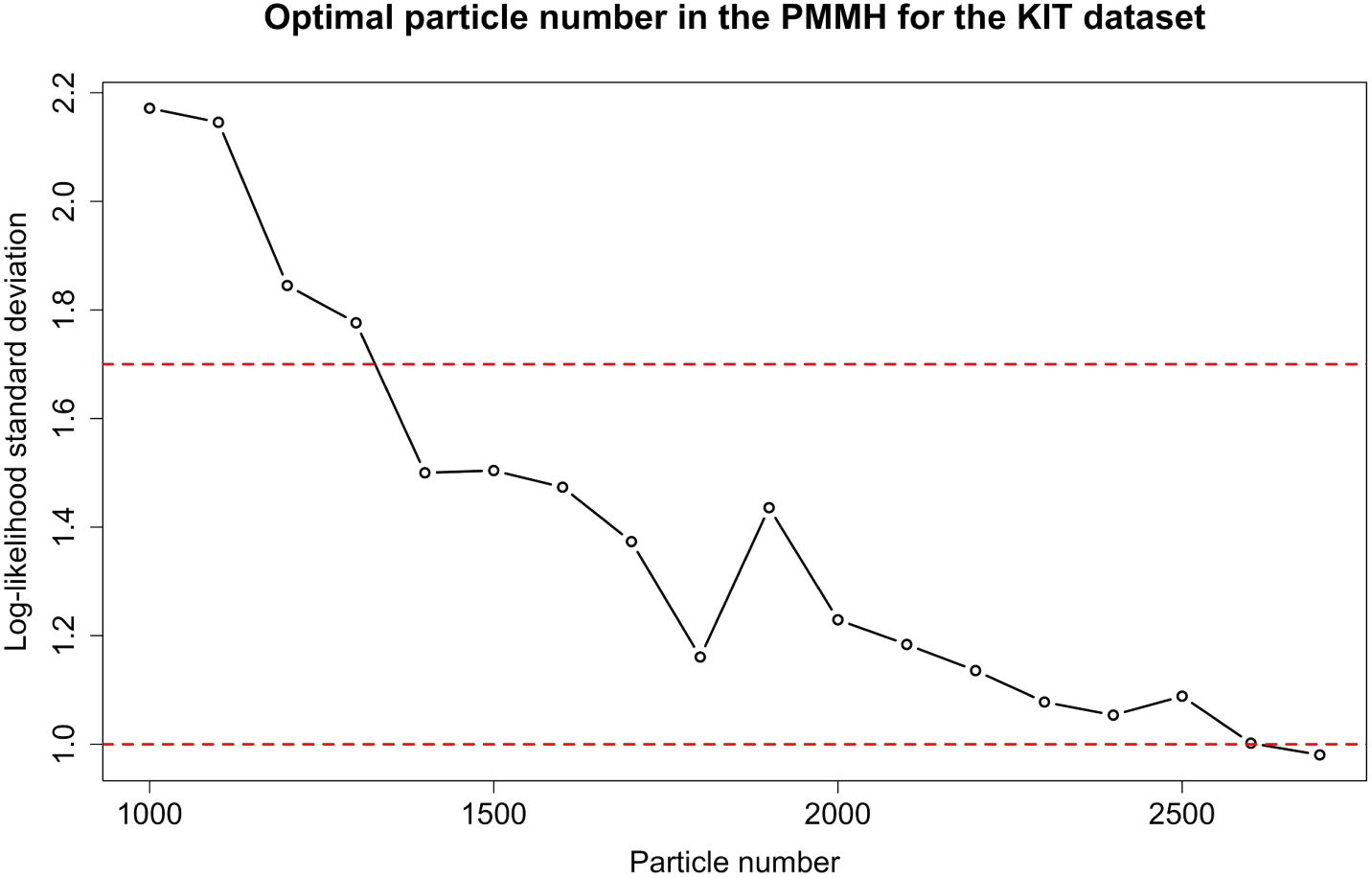
Changes in the standard deviation of the log-likelihood with the number of particles adopted in the PMMH algorithm for the *KIT* dataset.

Exact-approximate particle filtering approaches such as the PMMH algorithm we use in this work seem to be useful for the inference of population genetic parameters from time series data of allele frequencies. This methodology can be generalised to a range of complex evolutionary scenarios, *e.g*., changing population size. Although computationally demanding, improvements to the PMMH algorithm continue to be developed (*e.g*., Yildirim et al., 2018).

## 4. Discussion

In this work, we have developed a novel MCMC-based method to jointly infer natural selection at two linked loci from time series data of allele frequencies while explicitly taking in account effects of genetic recombination and local linkage. Our Bayesian inference procedure is built on an HMM framework incorporating the two-locus Wright-Fisher diffusion with selection. Our Bayesian estimates of the selection coefficients are achieved with the PMMH algorithm. We have demonstrated that our approach can accurately and efficiently estimate selection coefficients from simulated data, regardless of whether sampled chromosomes contain unknown alleles or not. We found that under certain circumstances, especially in the case of tightly linked loci, existing single-locus approaches fail to deliver precise estimates for selection coefficients, but our two-locus method still works well. We applied our Bayesian inference procedure to the *KIT* gene in horses, which is involved in the formation of white spotting patterns.

As noted earlier, the ancient horse DNA dataset has been the subject of earlier analyses by Malaspinas et al. (2012), Steinrücken et al. (2014), Schraiber et al. (2016) and He et al. (2019). Compared with many datasets describing experimental evolution under controlled laboratory or field mesocosm conditions, aDNA datasets are more likely to be composed of short degraded DNA fragments, typically with a high degree of genotyping error (Racimo et al., 2016). However, aDNA data provide an opportunity to investigate the chronology and tempo of natural selection across evolutionary timescales, which has an advantage of being associated with an interesting narrative (MacHugh et al., 2017). A motivation for the analysis is to see whether the statistical developments described here can shed further light on these data. We have found strong evidence showing that the sabino pattern caused by the *SB1* allele at locus *KIT16* has been selectively deleterious but no evidence showing that the tobiano pattern caused by the *KM1* allele at locus *KIT13* has been selectively advantageous. One explanation for our findings may be that there was a decreasing attractiveness of spotted horses since the Middle Ages due to the religious and cultural ideas (Wutke et al., 2016). Based on ancient Roman records, solid horses were preferred to spotted horses as the latter were considered to be of inferior quality. During medieval times, spotted horses had a negative connotation after several epidemics, resulting in a lower religious prestige for these patterns. Additionally, people might no longer see the need to distinguish wild (solid) horses from domesticated (spotted) horses as wild populations gradually became scarcer and approached extinction. Further reasons for the spotted horses being selectively deleterious might have been novel developments in weaponry such as the longbow, with these horses being an easier target than solid ones (see Wutke et al., 2016, and references therein).

In addition to our method, Terhorst et al. (2015) is the only existing approach that can model linked loci and genetic drift for the inference of natural selection from temporal changes in allele frequencies. In Terhorst et al. (2015), the underlying population dynamics at multiple linked loci was modelled using the Wright-Fisher model in their HMM framework, and the likelihood computation was carried out by approximating the Wright-Fisher model through a deterministic path with added Gaussian noises, which aims to fit a mathematically convenient transition probability density function by equating the first two moments of the Wright-Fisher model. Such a moment-based approximation works well for many applications when modelling the allele frequencies with intermediate values. However, as soon as the allele frequencies get close to their boundaries 0 or 1 (*i.e*., allele loss or fixation), the Wright-Fisher model will be poorly approximated due to the infinite support of the Gaussian distribution that will leak probability mass into the values of the allele frequency that are smaller than 0 or larger than 1, which is not mathematically possible. This issue becomes more problematic in the inference of natural selection since natural selection is expected to rapidly drive the allele frequencies towards the boundaries.

The MCMC-based method we have developed in this work is built on the standard diffusion limit of the Wright-Fisher model of the stochastic evolutionary dynamics under natural selection at a pair of linked loci, which is shown to be a good approximation even if the allele frequencies get close to their boundaries 0 or 1 (He et al., 2020). The diffusion approximation enables our approach to work well for the allele frequencies with all possible values. Our method can handle sampled chromosomes that contain unknown alleles, which one might expect to encounter in real data, especially in aDNA studies. Even though we have only illustrated the utility of our method on aDNA data in this work, our Bayesian inference procedure can also be used to analyse Pool-Seq time series data from E&R experiments, as in Terhorst et al. (2015). Given the PMMH algorithm we have used to infer natural selection in this work, our method lends itself naturally to joint estimates of the haplotype frequency trajectories of the underlying population without any increase in computational complexity. Moreover, our method can be readily extended to model a range of complex evolutionary scenarios, *e.g*., time-varying population size and selection coefficients, as it is built on simulating the Wright-Fisher diffusion.

One limitation of our approach is that we assume that mutant alleles were created before the initial sampling time point. Once a sample contains at least one copy of the mutant allele, we can reasonably assume that the mutant allele arose before the time of that sample. However in the case of earlier samples without any mutant allele, there is uncertainty in pinpointing when the mutant allele arose. This problem can be remedied by co-estimating the allele age as in *e.g*., Malaspinas et al. (2012), Schraiber et al. (2016) and He et al. (2019), but these works all investigate natural selection at a single locus. Jointly estimating the selection coefficients at linked loci along with the allele ages can be expected to be cumbersome as there are many cases to take into account. In the case of the ancient horse DNA data, we did not wish to make the assumption that the mutant alleles, *KM1* and *SB1*, arose earlier than the time that horses were domesticated. However, we can compare the inference results obtained with different choices of the initial sampling time point (see Supplemental Material, Tables S14-S16) and reach the same conclusion that there is no strong evidence for the *KM1* allele at locus *KIT13* to be positively selected, but there is strong evidence for the *SB1* allele at locus *KIT16* to be negatively selected.

Our Bayesian statistical framework lends itself to being extended to infer natural selection at multiple linked loci from time series data of allele frequencies, which might further improve the inference results of natural selection. The challenge is that with the increase in the number of linked loci, modelling the underlying population dynamics subject to natural selection becomes increasingly difficult. For example, there are eight haplotypes to take into account in the case of three linked loci each with two alleles. As a tractable alternative, we can apply our approach to multiple linked loci in a pairwise manner by using the PMMH algorithm within the Gibbs sampler, but it might only work for a small number of linked loci due to the computational cost of our two-locus approach. In practice, it will be necessary to find a good approximation of the Wright-Fisher model for the method to be computationally feasible, which will be the topic of future investigation. An important consideration is to what degree the results of the inference of natural selection are affected by the choice of stochastic or deterministic dynamics for the allele frequency trajectories (Jewett et al., 2016), and whether in many scenarios approximation with a deterministic model is satisfactory.

## Acknowledgements

This work was funded in part by the Engineering and Physical Sciences Research Council (EPSRC) Grant EP/I028498/1 to F.Y.

