## Supplemental Material for "Detecting and quantifying natural selection at two linked loci from time series data of allele frequencies with forward-in-time simulations"

#### File S1. Additional results for the analysis of simulated data

| | Bias | | RMSE | |
|---|---|---|---|---|
| $(s_{\mathcal{A}}, s_{\mathcal{B}})$ | $s_{\mathcal{A}}$ | $s_{\mathcal{B}}$ | $s_{\mathcal{A}}$ | $s_{\mathcal{B}}$ |
| $(0.3 \times 10^{-2}, 0.0 \times 10^{-2})$ | $0.567 \times 10^{-3}$ | $1.810 \times 10^{-3}$ | $3.798 \times 10^{-3}$ | $4.746 \times 10^{-3}$ |
| $(0.3 \times 10^{-2}, 0.2 \times 10^{-2})$ | $0.531 \times 10^{-3}$ | $1.593 \times 10^{-3}$ | $4.088 \times 10^{-3}$ | $4.291 \times 10^{-3}$ |
| $(0.3 \times 10^{-2}, 0.8 \times 10^{-2})$ | $1.047 \times 10^{-3}$ | $0.029 \times 10^{-3}$ | $4.451 \times 10^{-3}$ | $3.439 \times 10^{-3}$ |
| $(1.0 \times 10^{-2}, 0.0 \times 10^{-2})$ | $-0.890 \times 10^{-3}$ | $1.861 \times 10^{-3}$ | $3.084 \times 10^{-3}$ | $5.016 \times 10^{-3}$ |
| $(1.0 \times 10^{-2}, 0.2 \times 10^{-2})$ | $-0.111 \times 10^{-3}$ | $0.831 \times 10^{-3}$ | $3.250 \times 10^{-3}$ | $4.199 \times 10^{-3}$ |
| $(1.0 \times 10^{-2}, 0.8 \times 10^{-2})$ | $-0.781 \times 10^{-3}$ | $0.650 \times 10^{-3}$ | $3.749 \times 10^{-3}$ | $3.745 \times 10^{-3}$ |

(a) The case of tightly linked loci with the recombination rate $r = 0.00001$.

| | Bias | | RMSE | |
|---|---|---|---|---|
| $(s_{\mathcal{A}}, s_{\mathcal{B}})$ | $s_{\mathcal{A}}$ | $s_{\mathcal{B}}$ | $s_{\mathcal{A}}$ | $s_{\mathcal{B}}$ |
| $(0.3 \times 10^{-2}, 0.0 \times 10^{-2})$ | $0.649 \times 10^{-3}$ | $2.189 \times 10^{-3}$ | $3.501 \times 10^{-3}$ | $4.588 \times 10^{-3}$ |
| $(0.3 \times 10^{-2}, 0.2 \times 10^{-2})$ | $1.334 \times 10^{-3}$ | $1.411 \times 10^{-3}$ | $4.023 \times 10^{-3}$ | $3.331 \times 10^{-3}$ |
| $(0.3 \times 10^{-2}, 0.8 \times 10^{-2})$ | $0.579 \times 10^{-3}$ | $0.354 \times 10^{-3}$ | $2.940 \times 10^{-3}$ | $2.877 \times 10^{-3}$ |
| $(1.0 \times 10^{-2}, 0.0 \times 10^{-2})$ | $0.469 \times 10^{-3}$ | $2.056 \times 10^{-3}$ | $2.387 \times 10^{-3}$ | $4.299 \times 10^{-3}$ |
| $(1.0 \times 10^{-2}, 0.2 \times 10^{-2})$ | $0.437 \times 10^{-3}$ | $1.309 \times 10^{-3}$ | $2.499 \times 10^{-3}$ | $4.023 \times 10^{-3}$ |
| $(1.0 \times 10^{-2}, 0.8 \times 10^{-2})$ | $0.294 \times 10^{-3}$ | $0.890 \times 10^{-3}$ | $2.467 \times 10^{-3}$ | $2.971 \times 10^{-3}$ |

(b) The case of loosely linked loci with the recombination rate $r = 0.01$.

Table S1: Bias and RMSE of the MMSE estimates for 100 *allele frequency* datasets (*without* missing values) simulated with the initial population haplotype frequencies $\boldsymbol{x}_0 = (0.04, 0.08, 0.08, 0.8)$ and the dominance parameters $h_{\mathcal{A}} = 0.5$ and $h_{\mathcal{B}} = 0.5$.

| | Bias | | RMSE | |
|---|---|---|---|---|
| $(s_{\mathcal{A}}, s_{\mathcal{B}})$ | $s_{\mathcal{A}}$ | $s_{\mathcal{B}}$ | $s_{\mathcal{A}}$ | $s_{\mathcal{B}}$ |
| $(0.3 \times 10^{-2}, 0.0 \times 10^{-2})$ | $0.943 \times 10^{-3}$ | $1.389 \times 10^{-3}$ | $4.097 \times 10^{-3}$ | $4.500 \times 10^{-3}$ |
| $(0.3 \times 10^{-2}, 0.2 \times 10^{-2})$ | $1.246 \times 10^{-3}$ | $0.582 \times 10^{-3}$ | $4.949 \times 10^{-3}$ | $4.088 \times 10^{-3}$ |
| $(0.3 \times 10^{-2}, 0.8 \times 10^{-2})$ | $1.758 \times 10^{-3}$ | $-0.696 \times 10^{-3}$ | $4.744 \times 10^{-3}$ | $3.225 \times 10^{-3}$ |
| $(1.0 \times 10^{-2}, 0.0 \times 10^{-2})$ | $-1.249 \times 10^{-3}$ | $2.304 \times 10^{-3}$ | $3.138 \times 10^{-3}$ | $5.038 \times 10^{-3}$ |
| $(1.0 \times 10^{-2}, 0.2 \times 10^{-2})$ | $-0.878 \times 10^{-3}$ | $1.726 \times 10^{-3}$ | $2.880 \times 10^{-3}$ | $5.151 \times 10^{-3}$ |
| $(1.0 \times 10^{-2}, 0.8 \times 10^{-2})$ | $-0.107 \times 10^{-3}$ | $0.481 \times 10^{-3}$ | $3.700 \times 10^{-3}$ | $4.108 \times 10^{-3}$ |

(a) The case of tightly linked loci with the recombination rate $r = 0.00001$.

| | Bias | | RMSE | |
|---|---|---|---|---|
| $(s_{\mathcal{A}}, s_{\mathcal{B}})$ | $s_{\mathcal{A}}$ | $s_{\mathcal{B}}$ | $s_{\mathcal{A}}$ | $s_{\mathcal{B}}$ |
| $(0.3 \times 10^{-2}, 0.0 \times 10^{-2})$ | $0.434 \times 10^{-3}$ | $1.268 \times 10^{-3}$ | $3.014 \times 10^{-3}$ | $4.018 \times 10^{-3}$ |
| $(0.3 \times 10^{-2}, 0.2 \times 10^{-2})$ | $1.087 \times 10^{-3}$ | $1.138 \times 10^{-3}$ | $4.071 \times 10^{-3}$ | $3.892 \times 10^{-3}$ |
| $(0.3 \times 10^{-2}, 0.8 \times 10^{-2})$ | $1.570 \times 10^{-3}$ | $0.643 \times 10^{-3}$ | $3.538 \times 10^{-3}$ | $3.375 \times 10^{-3}$ |
| $(1.0 \times 10^{-2}, 0.0 \times 10^{-2})$ | $0.322 \times 10^{-3}$ | $2.458 \times 10^{-3}$ | $2.695 \times 10^{-3}$ | $4.952 \times 10^{-3}$ |
| $(1.0 \times 10^{-2}, 0.2 \times 10^{-2})$ | $-0.044 \times 10^{-3}$ | $1.344 \times 10^{-3}$ | $2.622 \times 10^{-3}$ | $4.485 \times 10^{-3}$ |
| $(1.0 \times 10^{-2}, 0.8 \times 10^{-2})$ | $0.271 \times 10^{-3}$ | $0.108 \times 10^{-3}$ | $2.663 \times 10^{-3}$ | $3.015 \times 10^{-3}$ |

(b) The case of loosely linked loci with the recombination rate $r = 0.01$.

Table S2: Bias and RMSE of the MMSE estimates for 100 *allele frequency* datasets (*with* 2% missing values) simulated with the initial population haplotype frequencies $\boldsymbol{x}_0 = (0.04, 0.08, 0.08, 0.8)$ and the dominance parameters $h_{\mathcal{A}} = 0.5$ and $h_{\mathcal{B}} = 0.5$.

| | Bias | | RMSE | |
|---|---|---|---|---|
| $(s_{\mathcal{A}}, s_{\mathcal{B}})$ | $s_{\mathcal{A}}$ | $s_{\mathcal{B}}$ | $s_{\mathcal{A}}$ | $s_{\mathcal{B}}$ |
| $(0.3 \times 10^{-2}, 0.0 \times 10^{-2})$ | $0.760 \times 10^{-3}$ | $1.571 \times 10^{-3}$ | $3.503 \times 10^{-3}$ | $4.329 \times 10^{-3}$ |
| $(0.3 \times 10^{-2}, 0.2 \times 10^{-2})$ | $0.895 \times 10^{-3}$ | $1.293 \times 10^{-3}$ | $3.559 \times 10^{-3}$ | $3.537 \times 10^{-3}$ |
| $(0.3 \times 10^{-2}, 0.8 \times 10^{-2})$ | $0.607 \times 10^{-3}$ | $0.608 \times 10^{-3}$ | $3.424 \times 10^{-3}$ | $3.091 \times 10^{-3}$ |
| $(1.0 \times 10^{-2}, 0.0 \times 10^{-2})$ | $-0.028 \times 10^{-3}$ | $1.023 \times 10^{-3}$ | $2.614 \times 10^{-3}$ | $3.592 \times 10^{-3}$ |
| $(1.0 \times 10^{-2}, 0.2 \times 10^{-2})$ | $0.735 \times 10^{-3}$ | $0.175 \times 10^{-3}$ | $3.042 \times 10^{-3}$ | $3.118 \times 10^{-3}$ |
| $(1.0 \times 10^{-2}, 0.8 \times 10^{-2})$ | $-0.239 \times 10^{-3}$ | $0.647 \times 10^{-3}$ | $2.876 \times 10^{-3}$ | $2.680 \times 10^{-3}$ |

(a) The case of tightly linked loci with the recombination rate $r = 0.00001$.

| | Bias | | RMSE | |
|---|---|---|---|---|
| $(s_{\mathcal{A}}, s_{\mathcal{B}})$ | $s_{\mathcal{A}}$ | $s_{\mathcal{B}}$ | $s_{\mathcal{A}}$ | $s_{\mathcal{B}}$ |
| $(0.3 \times 10^{-2}, 0.0 \times 10^{-2})$ | $0.698 \times 10^{-3}$ | $2.113 \times 10^{-3}$ | $3.512 \times 10^{-3}$ | $4.561 \times 10^{-3}$ |
| $(0.3 \times 10^{-2}, 0.2 \times 10^{-2})$ | $1.307 \times 10^{-3}$ | $1.331 \times 10^{-3}$ | $4.028 \times 10^{-3}$ | $3.291 \times 10^{-3}$ |
| $(0.3 \times 10^{-2}, 0.8 \times 10^{-2})$ | $0.449 \times 10^{-3}$ | $0.342 \times 10^{-3}$ | $2.921 \times 10^{-3}$ | $2.886 \times 10^{-3}$ |
| $(1.0 \times 10^{-2}, 0.0 \times 10^{-2})$ | $0.535 \times 10^{-3}$ | $1.859 \times 10^{-3}$ | $2.402 \times 10^{-3}$ | $4.131 \times 10^{-3}$ |
| $(1.0 \times 10^{-2}, 0.2 \times 10^{-2})$ | $0.438 \times 10^{-3}$ | $1.192 \times 10^{-3}$ | $2.506 \times 10^{-3}$ | $3.943 \times 10^{-3}$ |
| $(1.0 \times 10^{-2}, 0.8 \times 10^{-2})$ | $0.256 \times 10^{-3}$ | $0.776 \times 10^{-3}$ | $2.468 \times 10^{-3}$ | $2.862 \times 10^{-3}$ |

(b) The case of loosely linked loci with the recombination rate $r = 0.01$.

Table S3: Bias and RMSE of the MMSE estimates for 100 *haplotype frequency* datasets simulated with the initial population haplotype frequencies $\boldsymbol{x}_0 = (0.04, 0.08, 0.08, 0.8)$ and the dominance parameters $h_{\mathcal{A}} = 0.5$ and $h_{\mathcal{B}} = 0.5$.

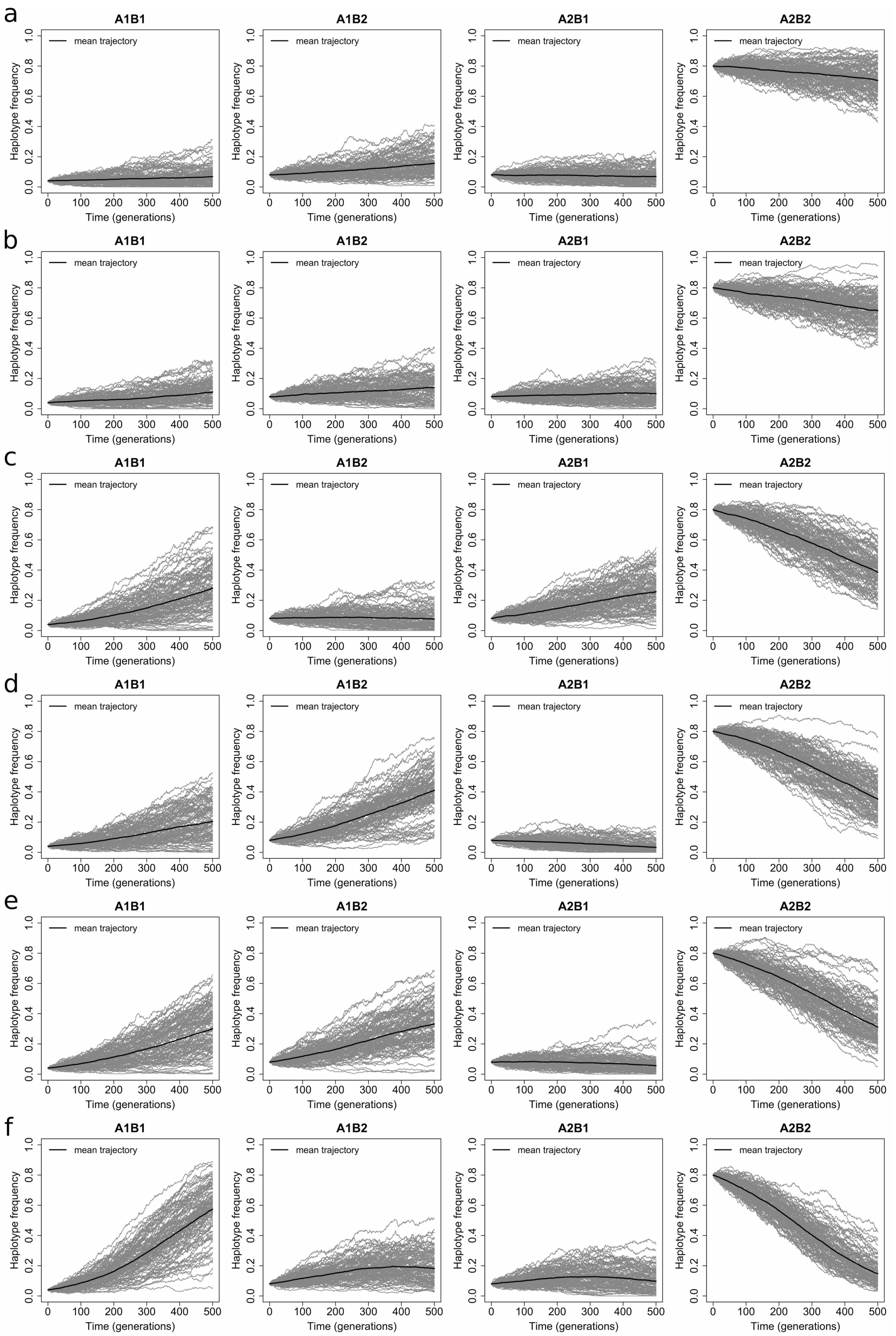


Figure S1: Haplotype frequency trajectories of the underlying population for 100 datasets simulated with the initial population haplotype frequencies $\boldsymbol{x}_0 = (0.04, 0.08, 0.08, 0.8)$ for the case of tightly linked loci with the recombination rate $r = 0.00001$, where the dominance parameters $h_{\mathcal{A}} = 0.5$ and $h_{\mathcal{B}} = 0.5$, and the selection coefficients (a) $s_{\mathcal{A}} = 0.003$ and $s_{\mathcal{B}} = 0$, (b) $s_{\mathcal{A}} = 0.003$ and $s_{\mathcal{B}} = 0.002$, (c) $s_{\mathcal{A}} = 0.003$ and $s_{\mathcal{B}} = 0.008$, (d) $s_{\mathcal{A}} = 0.01$ and $s_{\mathcal{B}} = 0$, (e) $s_{\mathcal{A}} = 0.01$ and $s_{\mathcal{B}} = 0.002$, and (f) $s_{\mathcal{A}} = 0.01$ and $s_{\mathcal{B}} = 0.008$.

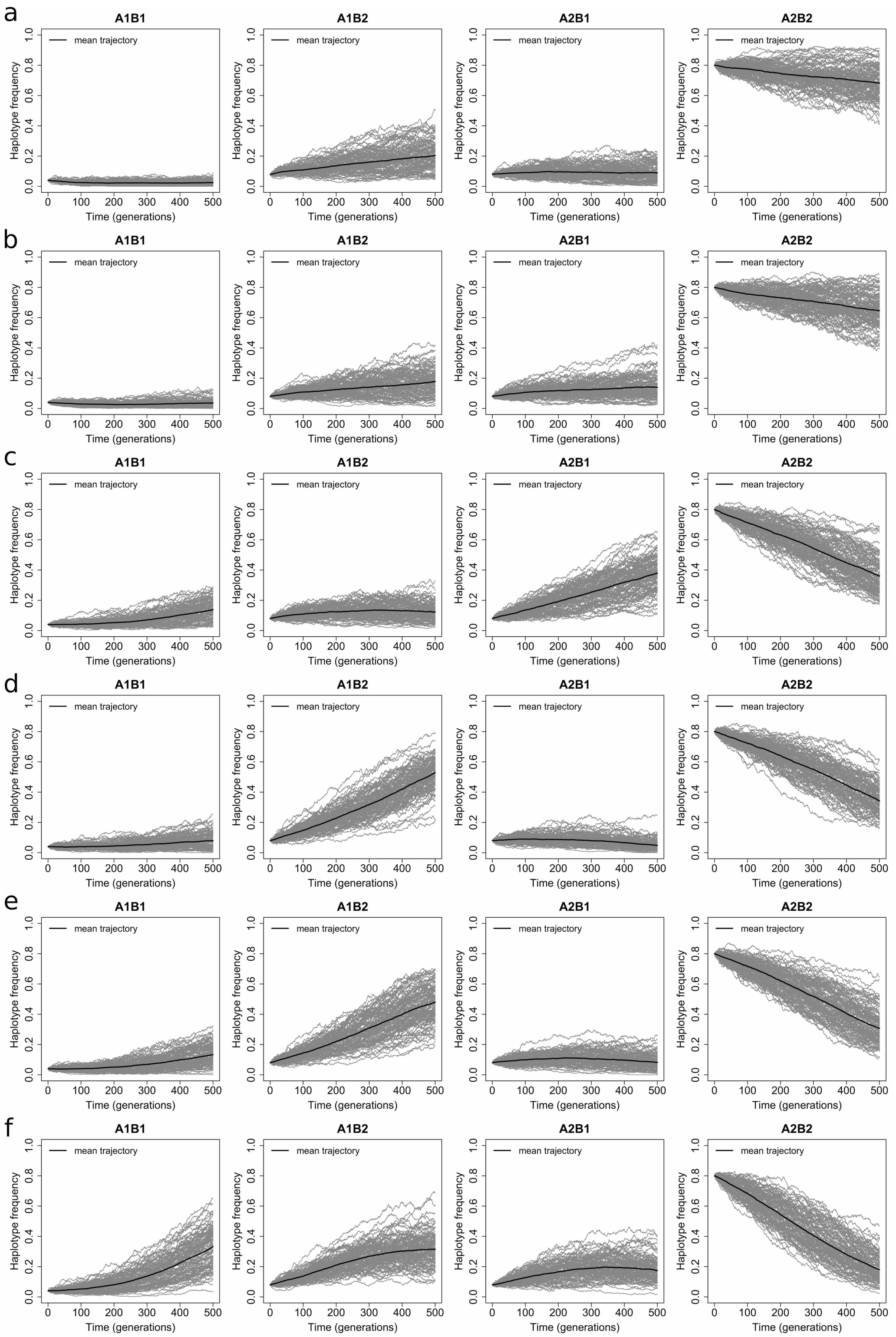


Figure S2: Haplotype frequency trajectories of the underlying population for 100 datasets simulated with the initial population haplotype frequencies $\boldsymbol{x}_0 = (0.04, 0.08, 0.08, 0.8)$ for the case of loosely linked loci with the recombination rate $r = 0.01$, where the dominance parameters $h_{\mathcal{A}} = 0.5$ and $h_{\mathcal{B}} = 0.5$, and the selection coefficients (a) $s_{\mathcal{A}} = 0.003$ and $s_{\mathcal{B}} = 0$, (b) $s_{\mathcal{A}} = 0.003$ and $s_{\mathcal{B}} = 0.002$, (c) $s_{\mathcal{A}} = 0.003$ and $s_{\mathcal{B}} = 0.008$, (d) $s_{\mathcal{A}} = 0.01$ and $s_{\mathcal{B}} = 0$, (e) $s_{\mathcal{A}} = 0.01$ and $s_{\mathcal{B}} = 0.002$, and (f) $s_{\mathcal{A}} = 0.01$ and $s_{\mathcal{B}} = 0.008$.

| | Bias | | RMSE | |
|---|---|---|---|---|
| $(s_{\mathcal{A}}, s_{\mathcal{B}})$ | $s_{\mathcal{A}}$ | $s_{\mathcal{B}}$ | $s_{\mathcal{A}}$ | $s_{\mathcal{B}}$ |
| $(0.3 \times 10^{-2}, 0.0 \times 10^{-2})$ | $0.359 \times 10^{-3}$ | $-0.123 \times 10^{-3}$ | $2.403 \times 10^{-3}$ | $2.602 \times 10^{-3}$ |
| $(0.3 \times 10^{-2}, 0.2 \times 10^{-2})$ | $0.292 \times 10^{-3}$ | $0.347 \times 10^{-3}$ | $2.333 \times 10^{-3}$ | $2.155 \times 10^{-3}$ |
| $(0.3 \times 10^{-2}, 0.8 \times 10^{-2})$ | $0.070 \times 10^{-3}$ | $-0.015 \times 10^{-3}$ | $2.686 \times 10^{-3}$ | $2.328 \times 10^{-3}$ |
| $(1.0 \times 10^{-2}, 0.0 \times 10^{-2})$ | $-0.064 \times 10^{-3}$ | $0.187 \times 10^{-3}$ | $2.551 \times 10^{-3}$ | $2.295 \times 10^{-3}$ |
| $(1.0 \times 10^{-2}, 0.2 \times 10^{-2})$ | $-0.465 \times 10^{-3}$ | $-0.263 \times 10^{-3}$ | $2.550 \times 10^{-3}$ | $2.356 \times 10^{-3}$ |
| $(1.0 \times 10^{-2}, 0.8 \times 10^{-2})$ | $-0.080 \times 10^{-3}$ | $-0.155 \times 10^{-3}$ | $2.659 \times 10^{-3}$ | $2.451 \times 10^{-3}$ |

(a) The case of tightly linked loci with the recombination rate $r = 0.00001$.

| | Bias | | RMSE | |
|---|---|---|---|---|
| $(s_{\mathcal{A}}, s_{\mathcal{B}})$ | $s_{\mathcal{A}}$ | $s_{\mathcal{B}}$ | $s_{\mathcal{A}}$ | $s_{\mathcal{B}}$ |
| $(0.3 \times 10^{-2}, 0.0 \times 10^{-2})$ | $-0.008 \times 10^{-3}$ | $0.034 \times 10^{-3}$ | $2.464 \times 10^{-3}$ | $2.295 \times 10^{-3}$ |
| $(0.3 \times 10^{-2}, 0.2 \times 10^{-2})$ | $0.462 \times 10^{-3}$ | $-0.141 \times 10^{-3}$ | $2.421 \times 10^{-3}$ | $2.529 \times 10^{-3}$ |
| $(0.3 \times 10^{-2}, 0.8 \times 10^{-2})$ | $0.015 \times 10^{-3}$ | $-0.514 \times 10^{-3}$ | $2.367 \times 10^{-3}$ | $2.507 \times 10^{-3}$ |
| $(1.0 \times 10^{-2}, 0.0 \times 10^{-2})$ | $0.292 \times 10^{-3}$ | $0.173 \times 10^{-3}$ | $2.467 \times 10^{-3}$ | $2.158 \times 10^{-3}$ |
| $(1.0 \times 10^{-2}, 0.2 \times 10^{-2})$ | $0.012 \times 10^{-3}$ | $0.037 \times 10^{-3}$ | $2.566 \times 10^{-3}$ | $2.272 \times 10^{-3}$ |
| $(1.0 \times 10^{-2}, 0.8 \times 10^{-2})$ | $-0.011 \times 10^{-3}$ | $-0.033 \times 10^{-3}$ | $2.373 \times 10^{-3}$ | $2.527 \times 10^{-3}$ |

(b) The case of loosely linked loci with the recombination rate $r = 0.01$.

Table S4: Bias and RMSE of the MMSE estimates for 100 *haplotype frequency* datasets simulated with the initial population haplotype frequencies $\boldsymbol{x}_0 = (0.1, 0.2, 0.3, 0.4)$ and the dominance parameters $h_{\mathcal{A}} = 0.5$ and $h_{\mathcal{B}} = 0.5$.

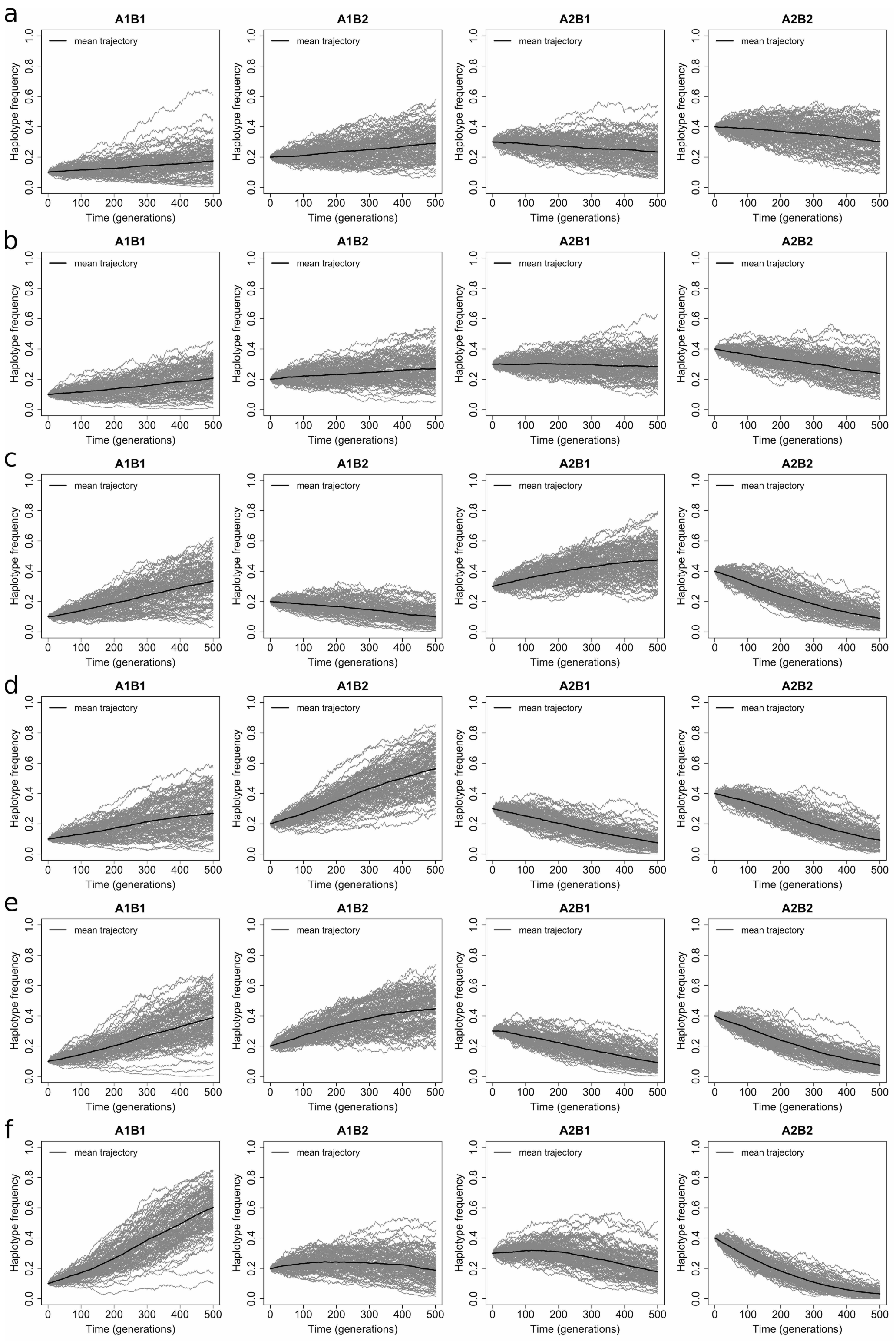


Figure S3: Haplotype frequency trajectories of the underlying population for 100 datasets simulated with the initial population haplotype frequencies $\boldsymbol{x}_0 = (0.1, 0.2, 0.3, 0.4)$ for the case of tightly linked loci with the recombination rate $r = 0.00001$, where the dominance parameters $h_{\mathcal{A}} = 0.5$ and $h_{\mathcal{B}} = 0.5$, and the selection coefficients (a) $s_{\mathcal{A}} = 0.003$ and $s_{\mathcal{B}} = 0$, (b) $s_{\mathcal{A}} = 0.003$ and $s_{\mathcal{B}} = 0.002$, (c) $s_{\mathcal{A}} = 0.003$ and $s_{\mathcal{B}} = 0.008$, (d) $s_{\mathcal{A}} = 0.01$ and $s_{\mathcal{B}} = 0$, (e) $s_{\mathcal{A}} = 0.01$ and $s_{\mathcal{B}} = 0.002$, and (f) $s_{\mathcal{A}} = 0.01$ and $s_{\mathcal{B}} = 0.008$.

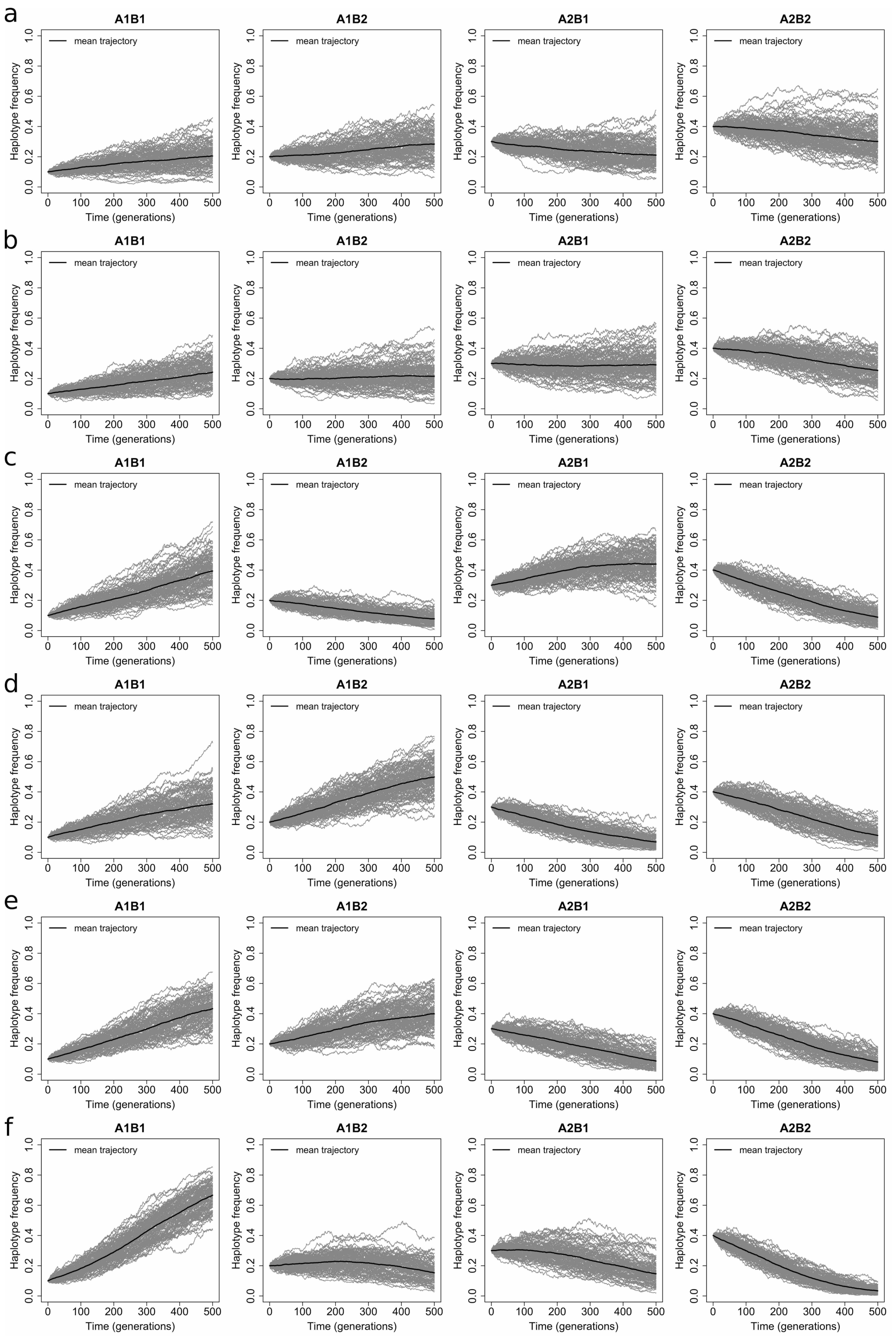


Figure S4: Haplotype frequency trajectories of the underlying population for 100 datasets simulated with the initial population haplotype frequencies $\boldsymbol{x}_0 = (0.1, 0.2, 0.3, 0.4)$ for the case of loosely linked loci with the recombination rate $r = 0.01$, where the dominance parameters $h_{\mathcal{A}} = 0.5$ and $h_{\mathcal{B}} = 0.5$, and the selection coefficients (a) $s_{\mathcal{A}} = 0.003$ and $s_{\mathcal{B}} = 0$, (b) $s_{\mathcal{A}} = 0.003$ and $s_{\mathcal{B}} = 0.002$, (c) $s_{\mathcal{A}} = 0.003$ and $s_{\mathcal{B}} = 0.008$, (d) $s_{\mathcal{A}} = 0.01$ and $s_{\mathcal{B}} = 0$, (e) $s_{\mathcal{A}} = 0.01$ and $s_{\mathcal{B}} = 0.002$, and (f) $s_{\mathcal{A}} = 0.01$ and $s_{\mathcal{B}} = 0.008$.

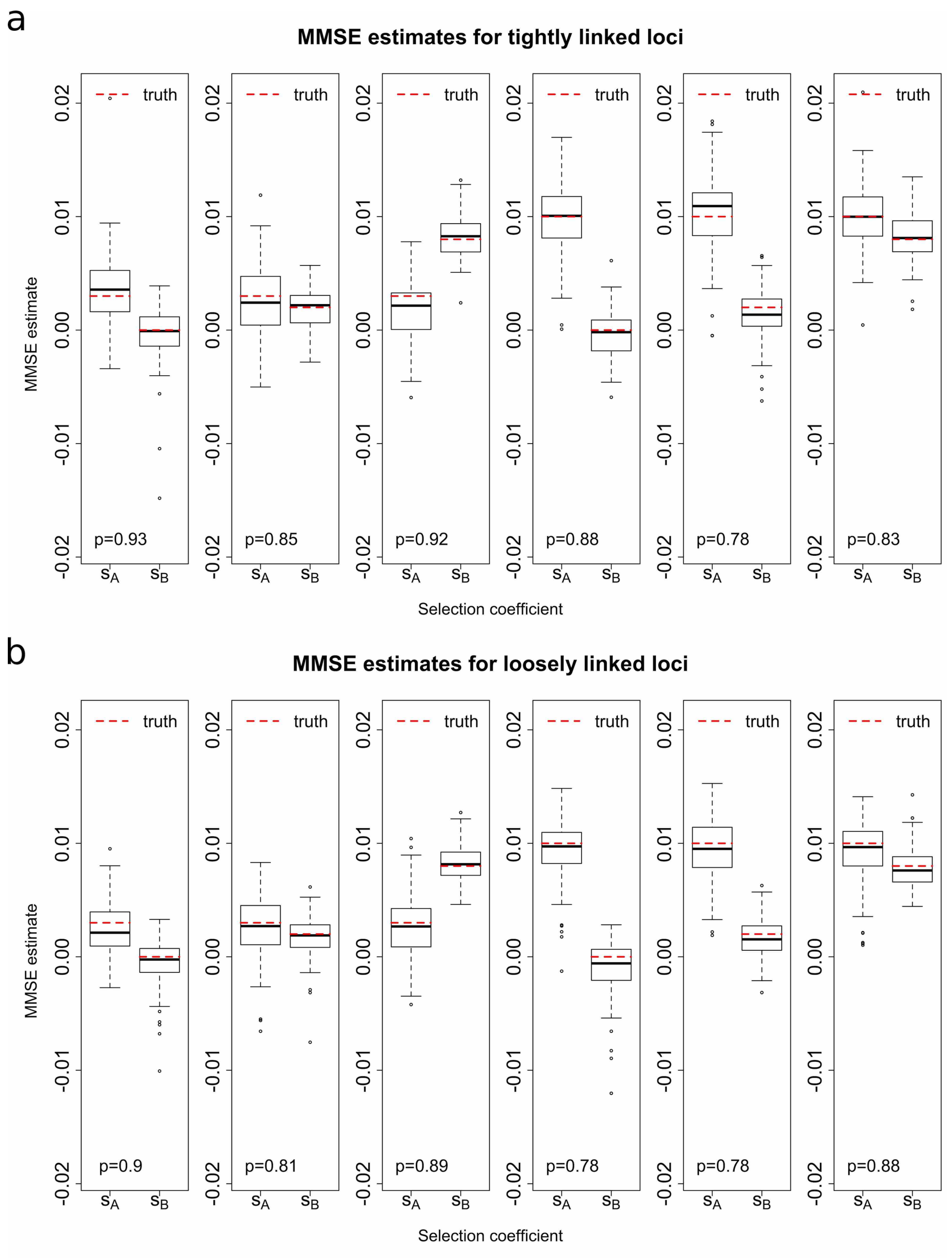


Figure S5: Empirical distributions of the MMSE estimates for 100 *haplotype frequency* datasets simulated with the initial population haplotype frequencies $\boldsymbol{x}_0 = (0.0001, 0, 0.1999, 0.8)$ and the dominance parameters $h_{\mathcal{A}} = 0.5$ and $h_{\mathcal{B}} = 0.5$ for the case of (a) tightly linked loci with the recombination rate $r = 0.00001$ and (b) loosely linked loci with the recombination rate $r = 0.01$. The $p$ value in the bottom left corner indicates the proportion of the runs where the true values of the selection coefficients both fall within their 95% HPD intervals. It should be noted that in this case we condition the mutant alleles at both loci to survive until the most recent sampling time point and sample 50 chromosomes from the underlying population at every 120 generations throughout 1200 generations.

| | Bias | | RMSE | |
|---|---|---|---|---|
| $(s_{\mathcal{A}}, s_{\mathcal{B}})$ | $s_{\mathcal{A}}$ | $s_{\mathcal{B}}$ | $s_{\mathcal{A}}$ | $s_{\mathcal{B}}$ |
| $(0.3 \times 10^{-2}, 0.0 \times 10^{-2})$ | $-0.404 \times 10^{-3}$ | $0.336 \times 10^{-3}$ | $3.259 \times 10^{-3}$ | $2.496 \times 10^{-3}$ |
| $(0.3 \times 10^{-2}, 0.2 \times 10^{-2})$ | $0.549 \times 10^{-3}$ | $0.091 \times 10^{-3}$ | $3.103 \times 10^{-3}$ | $1.715 \times 10^{-3}$ |
| $(0.3 \times 10^{-2}, 0.8 \times 10^{-2})$ | $1.329 \times 10^{-3}$ | $-0.279 \times 10^{-3}$ | $2.980 \times 10^{-3}$ | $1.947 \times 10^{-3}$ |
| $(1.0 \times 10^{-2}, 0.0 \times 10^{-2})$ | $0.212 \times 10^{-3}$ | $0.436 \times 10^{-3}$ | $2.992 \times 10^{-3}$ | $2.091 \times 10^{-3}$ |
| $(1.0 \times 10^{-2}, 0.2 \times 10^{-2})$ | $-0.334 \times 10^{-3}$ | $0.600 \times 10^{-3}$ | $3.515 \times 10^{-3}$ | $2.296 \times 10^{-3}$ |
| $(1.0 \times 10^{-2}, 0.8 \times 10^{-2})$ | $-0.022 \times 10^{-3}$ | $-0.230 \times 10^{-3}$ | $2.687 \times 10^{-3}$ | $2.072 \times 10^{-3}$ |

(a) The case of tightly linked loci with the recombination rate $r = 0.00001$.

| | Bias | | RMSE | |
|---|---|---|---|---|
| $(s_{\mathcal{A}}, s_{\mathcal{B}})$ | $s_{\mathcal{A}}$ | $s_{\mathcal{B}}$ | $s_{\mathcal{A}}$ | $s_{\mathcal{B}}$ |
| $(0.3 \times 10^{-2}, 0.0 \times 10^{-2})$ | $0.515 \times 10^{-3}$ | $0.577 \times 10^{-3}$ | $2.233 \times 10^{-3}$ | $2.247 \times 10^{-3}$ |
| $(0.3 \times 10^{-2}, 0.2 \times 10^{-2})$ | $0.493 \times 10^{-3}$ | $0.246 \times 10^{-3}$ | $2.934 \times 10^{-3}$ | $1.953 \times 10^{-3}$ |
| $(0.3 \times 10^{-2}, 0.8 \times 10^{-2})$ | $0.401 \times 10^{-3}$ | $-0.231 \times 10^{-3}$ | $2.929 \times 10^{-3}$ | $1.505 \times 10^{-3}$ |
| $(1.0 \times 10^{-2}, 0.0 \times 10^{-2})$ | $0.611 \times 10^{-3}$ | $1.185 \times 10^{-3}$ | $2.544 \times 10^{-3}$ | $3.032 \times 10^{-3}$ |
| $(1.0 \times 10^{-2}, 0.2 \times 10^{-2})$ | $0.503 \times 10^{-3}$ | $0.339 \times 10^{-3}$ | $2.728 \times 10^{-3}$ | $1.744 \times 10^{-3}$ |
| $(1.0 \times 10^{-2}, 0.8 \times 10^{-2})$ | $0.743 \times 10^{-3}$ | $0.078 \times 10^{-3}$ | $2.895 \times 10^{-3}$ | $1.751 \times 10^{-3}$ |

(b) The case of loosely linked loci with the recombination rate $r = 0.01$.

Table S5: Bias and RMSE of the MMSE estimates for 100 *haplotype frequency* datasets simulated with the initial population haplotype frequencies $\boldsymbol{x}_0 = (0.0001, 0, 0.1999, 0.8)$ and the dominance parameters $h_{\mathcal{A}} = 0.5$ and $h_{\mathcal{B}} = 0.5$. It should be noted that in this case we condition the mutant alleles at both loci to survive until the most recent sampling time point and sample 50 chromosomes from the underlying population at every 120 generations throughout 1200 generations.

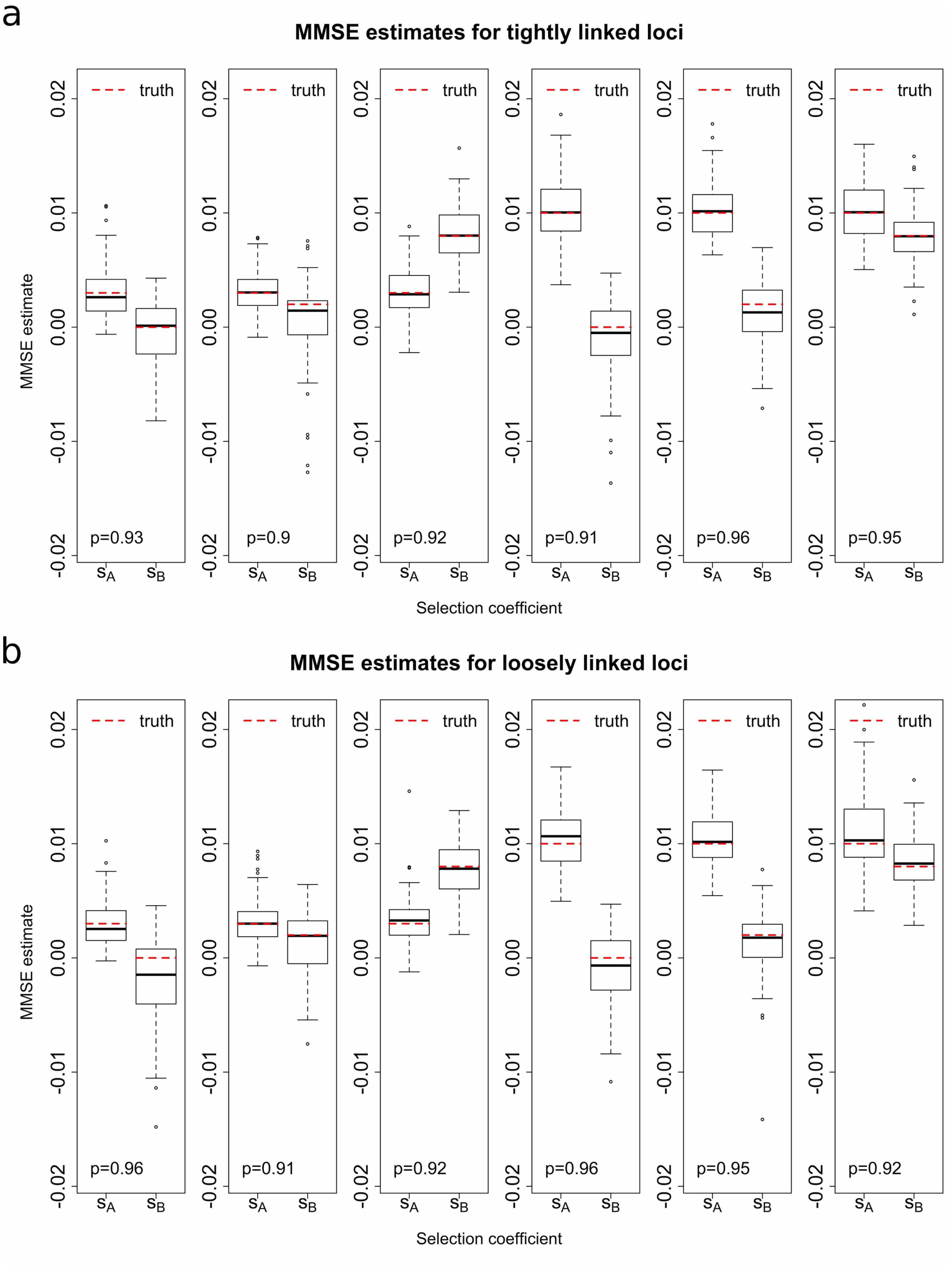


Figure S6: Empirical distributions of the MMSE estimates for 100 *haplotype frequency* datasets simulated with the initial population haplotype frequencies $\boldsymbol{x}_0 = (0.1, 0.2, 0.3, 0.4)$ and the dominance parameters $h_{\mathcal{A}} = 0$ and $h_{\mathcal{B}} = 1$ for the case of (a) tightly linked loci with the recombination rate $r = 0.00001$ and (b) loosely linked loci with the recombination rate $r = 0.01$. The $p$ value in the bottom left corner indicates the proportion of the runs where the true values of the selection coefficients both fall within their 95% HPD intervals.

| | Bias | | RMSE | |
|---|---|---|---|---|
| $(s_\mathcal{A}, s_\mathcal{B})$ | $s_\mathcal{A}$ | $s_\mathcal{B}$ | $s_\mathcal{A}$ | $s_\mathcal{B}$ |
| $(0.3 \times 10^{-2}, 0.0 \times 10^{-2})$ | $-0.026 \times 10^{-3}$ | $0.402 \times 10^{-3}$ | $2.269 \times 10^{-3}$ | $2.752 \times 10^{-3}$ |
| $(0.3 \times 10^{-2}, 0.2 \times 10^{-2})$ | $-0.123 \times 10^{-3}$ | $1.271 \times 10^{-3}$ | $1.885 \times 10^{-3}$ | $3.581 \times 10^{-3}$ |
| $(0.3 \times 10^{-2}, 0.8 \times 10^{-2})$ | $-0.020 \times 10^{-3}$ | $-0.202 \times 10^{-3}$ | $2.089 \times 10^{-3}$ | $2.261 \times 10^{-3}$ |
| $(1.0 \times 10^{-2}, 0.0 \times 10^{-2})$ | $-0.369 \times 10^{-3}$ | $0.906 \times 10^{-3}$ | $2.941 \times 10^{-3}$ | $3.403 \times 10^{-3}$ |
| $(1.0 \times 10^{-2}, 0.2 \times 10^{-2})$ | $-0.178 \times 10^{-3}$ | $0.901 \times 10^{-3}$ | $2.436 \times 10^{-3}$ | $2.982 \times 10^{-3}$ |
| $(1.0 \times 10^{-2}, 0.8 \times 10^{-2})$ | $-0.109 \times 10^{-3}$ | $-0.005 \times 10^{-3}$ | $2.330 \times 10^{-3}$ | $2.261 \times 10^{-3}$ |

(a) The case of tightly linked loci with the recombination rate $r = 0.00001$.

| | Bias | | RMSE | |
|---|---|---|---|---|
| $(s_\mathcal{A}, s_\mathcal{B})$ | $s_\mathcal{A}$ | $s_\mathcal{B}$ | $s_\mathcal{A}$ | $s_\mathcal{B}$ |
| $(0.3 \times 10^{-2}, 0.0 \times 10^{-2})$ | $0.007 \times 10^{-3}$ | $1.693 \times 10^{-3}$ | $1.931 \times 10^{-3}$ | $3.938 \times 10^{-3}$ |
| $(0.3 \times 10^{-2}, 0.2 \times 10^{-2})$ | $-0.151 \times 10^{-3}$ | $0.792 \times 10^{-3}$ | $2.013 \times 10^{-3}$ | $2.900 \times 10^{-3}$ |
| $(0.3 \times 10^{-2}, 0.8 \times 10^{-2})$ | $-0.286 \times 10^{-3}$ | $0.209 \times 10^{-3}$ | $2.059 \times 10^{-3}$ | $2.265 \times 10^{-3}$ |
| $(1.0 \times 10^{-2}, 0.0 \times 10^{-2})$ | $-0.549 \times 10^{-3}$ | $0.947 \times 10^{-3}$ | $2.898 \times 10^{-3}$ | $3.554 \times 10^{-3}$ |
| $(1.0 \times 10^{-2}, 0.2 \times 10^{-2})$ | $-0.477 \times 10^{-3}$ | $0.663 \times 10^{-3}$ | $2.594 \times 10^{-3}$ | $2.877 \times 10^{-3}$ |
| $(1.0 \times 10^{-2}, 0.8 \times 10^{-2})$ | $-0.870 \times 10^{-3}$ | $-0.295 \times 10^{-3}$ | $3.403 \times 10^{-3}$ | $2.324 \times 10^{-3}$ |

(b) The case of loosely linked loci with the recombination rate $r = 0.01$.

Table S6: Bias and RMSE of the MMSE estimates for 100 *haplotype frequency* datasets simulated with the initial population haplotype frequencies $\boldsymbol{x}_0 = (0.1, 0.2, 0.3, 0.4)$ and the dominance parameters $h_\mathcal{A} = 0$ and $h_\mathcal{B} = 1$.

| | Bias | | RMSE | |
|---|---|---|---|---|
| $(s_\mathcal{A}, s_\mathcal{B})$ | $s_\mathcal{A}$ | $s_\mathcal{B}$ | $s_\mathcal{A}$ | $s_\mathcal{B}$ |
| $(0.3 \times 10^{-2}, 0.0 \times 10^{-2})$ | $0.722 \times 10^{-3}$ | $1.731 \times 10^{-3}$ | $3.713 \times 10^{-3}$ | $4.629 \times 10^{-3}$ |
| $(0.3 \times 10^{-2}, 0.2 \times 10^{-2})$ | $0.826 \times 10^{-3}$ | $1.445 \times 10^{-3}$ | $3.891 \times 10^{-3}$ | $4.040 \times 10^{-3}$ |
| $(0.3 \times 10^{-2}, 0.8 \times 10^{-2})$ | $0.954 \times 10^{-3}$ | $0.319 \times 10^{-3}$ | $4.429 \times 10^{-3}$ | $3.387 \times 10^{-3}$ |
| $(1.0 \times 10^{-2}, 0.0 \times 10^{-2})$ | $-0.403 \times 10^{-3}$ | $1.774 \times 10^{-3}$ | $3.285 \times 10^{-3}$ | $4.853 \times 10^{-3}$ |
| $(1.0 \times 10^{-2}, 0.2 \times 10^{-2})$ | $0.381 \times 10^{-3}$ | $0.837 \times 10^{-3}$ | $3.370 \times 10^{-3}$ | $4.314 \times 10^{-3}$ |
| $(1.0 \times 10^{-2}, 0.8 \times 10^{-2})$ | $-0.243 \times 10^{-3}$ | $0.935 \times 10^{-3}$ | $3.544 \times 10^{-3}$ | $3.465 \times 10^{-3}$ |

(a) The case of tightly linked loci with the recombination rate $r = 0.00001$.

| | Bias | | RMSE | |
|---|---|---|---|---|
| $(s_\mathcal{A}, s_\mathcal{B})$ | $s_\mathcal{A}$ | $s_\mathcal{B}$ | $s_\mathcal{A}$ | $s_\mathcal{B}$ |
| $(0.3 \times 10^{-2}, 0.0 \times 10^{-2})$ | $0.522 \times 10^{-3}$ | $2.150 \times 10^{-3}$ | $3.556 \times 10^{-3}$ | $4.640 \times 10^{-3}$ |
| $(0.3 \times 10^{-2}, 0.2 \times 10^{-2})$ | $1.223 \times 10^{-3}$ | $1.453 \times 10^{-3}$ | $4.064 \times 10^{-3}$ | $3.556 \times 10^{-3}$ |
| $(0.3 \times 10^{-2}, 0.8 \times 10^{-2})$ | $0.773 \times 10^{-3}$ | $0.424 \times 10^{-3}$ | $3.063 \times 10^{-3}$ | $2.987 \times 10^{-3}$ |
| $(1.0 \times 10^{-2}, 0.0 \times 10^{-2})$ | $0.481 \times 10^{-3}$ | $1.939 \times 10^{-3}$ | $2.454 \times 10^{-3}$ | $4.393 \times 10^{-3}$ |
| $(1.0 \times 10^{-2}, 0.2 \times 10^{-2})$ | $0.506 \times 10^{-3}$ | $1.302 \times 10^{-3}$ | $2.642 \times 10^{-3}$ | $3.973 \times 10^{-3}$ |
| $(1.0 \times 10^{-2}, 0.8 \times 10^{-2})$ | $0.284 \times 10^{-3}$ | $0.920 \times 10^{-3}$ | $2.662 \times 10^{-3}$ | $3.058 \times 10^{-3}$ |

(b) The case of loosely linked loci with the recombination rate $r = 0.01$.

Table S7: Bias and RMSE of the MAP estimates for 100 *allele frequency* datasets (*without* missing values) simulated with the initial population haplotype frequencies $\boldsymbol{x}_0 = (0.04, 0.08, 0.08, 0.8)$ and the dominance parameters $h_\mathcal{A} = 0.5$ and $h_\mathcal{B} = 0.5$.

| | Bias | | RMSE | |
|---|---|---|---|---|
| $(s_{\mathcal{A}}, s_{\mathcal{B}})$ | $s_{\mathcal{A}}$ | $s_{\mathcal{B}}$ | $s_{\mathcal{A}}$ | $s_{\mathcal{B}}$ |
| $(0.3 \times 10^{-2}, 0.0 \times 10^{-2})$ | $0.926 \times 10^{-3}$ | $1.427 \times 10^{-3}$ | $3.950 \times 10^{-3}$ | $4.306 \times 10^{-3}$ |
| $(0.3 \times 10^{-2}, 0.2 \times 10^{-2})$ | $1.201 \times 10^{-3}$ | $0.589 \times 10^{-3}$ | $4.681 \times 10^{-3}$ | $4.044 \times 10^{-3}$ |
| $(0.3 \times 10^{-2}, 0.8 \times 10^{-2})$ | $1.722 \times 10^{-3}$ | $-0.368 \times 10^{-3}$ | $5.140 \times 10^{-3}$ | $3.435 \times 10^{-3}$ |
| $(1.0 \times 10^{-2}, 0.0 \times 10^{-2})$ | $-0.735 \times 10^{-3}$ | $2.394 \times 10^{-3}$ | $3.352 \times 10^{-3}$ | $5.579 \times 10^{-3}$ |
| $(1.0 \times 10^{-2}, 0.2 \times 10^{-2})$ | $-0.413 \times 10^{-3}$ | $1.719 \times 10^{-3}$ | $2.871 \times 10^{-3}$ | $5.372 \times 10^{-3}$ |
| $(1.0 \times 10^{-2}, 0.8 \times 10^{-2})$ | $0.308 \times 10^{-3}$ | $0.822 \times 10^{-3}$ | $3.502 \times 10^{-3}$ | $4.072 \times 10^{-3}$ |

(a) The case of tightly linked loci with the recombination rate $r = 0.00001$.

| | Bias | | RMSE | |
|---|---|---|---|---|
| $(s_{\mathcal{A}}, s_{\mathcal{B}})$ | $s_{\mathcal{A}}$ | $s_{\mathcal{B}}$ | $s_{\mathcal{A}}$ | $s_{\mathcal{B}}$ |
| $(0.3 \times 10^{-2}, 0.0 \times 10^{-2})$ | $0.515 \times 10^{-3}$ | $1.409 \times 10^{-3}$ | $3.191 \times 10^{-3}$ | $3.937 \times 10^{-3}$ |
| $(0.3 \times 10^{-2}, 0.2 \times 10^{-2})$ | $0.943 \times 10^{-3}$ | $1.112 \times 10^{-3}$ | $3.997 \times 10^{-3}$ | $3.934 \times 10^{-3}$ |
| $(0.3 \times 10^{-2}, 0.8 \times 10^{-2})$ | $1.506 \times 10^{-3}$ | $0.828 \times 10^{-3}$ | $3.640 \times 10^{-3}$ | $3.573 \times 10^{-3}$ |
| $(1.0 \times 10^{-2}, 0.0 \times 10^{-2})$ | $0.168 \times 10^{-3}$ | $2.365 \times 10^{-3}$ | $2.964 \times 10^{-3}$ | $4.715 \times 10^{-3}$ |
| $(1.0 \times 10^{-2}, 0.2 \times 10^{-2})$ | $0.211 \times 10^{-3}$ | $1.182 \times 10^{-3}$ | $2.802 \times 10^{-3}$ | $4.506 \times 10^{-3}$ |
| $(1.0 \times 10^{-2}, 0.8 \times 10^{-2})$ | $0.220 \times 10^{-3}$ | $0.284 \times 10^{-3}$ | $2.721 \times 10^{-3}$ | $3.005 \times 10^{-3}$ |

(b) The case of loosely linked loci with the recombination rate $r = 0.01$.

Table S8: Bias and RMSE of the MAP estimates for 100 *allele frequency* datasets (*with* 2% missing values) simulated with the initial population haplotype frequencies $\boldsymbol{x}_0 = (0.04, 0.08, 0.08, 0.8)$ and the dominance parameters $h_{\mathcal{A}} = 0.5$ and $h_{\mathcal{B}} = 0.5$.

| | Bias | | RMSE | |
|---|---|---|---|---|
| $(s_\mathcal{A}, s_\mathcal{B})$ | $s_\mathcal{A}$ | $s_\mathcal{B}$ | $s_\mathcal{A}$ | $s_\mathcal{B}$ |
| $(0.3 \times 10^{-2}, 0.0 \times 10^{-2})$ | $0.736 \times 10^{-3}$ | $1.299 \times 10^{-3}$ | $3.487 \times 10^{-3}$ | $4.114 \times 10^{-3}$ |
| $(0.3 \times 10^{-2}, 0.2 \times 10^{-2})$ | $0.946 \times 10^{-3}$ | $1.171 \times 10^{-3}$ | $3.601 \times 10^{-3}$ | $3.666 \times 10^{-3}$ |
| $(0.3 \times 10^{-2}, 0.8 \times 10^{-2})$ | $0.641 \times 10^{-3}$ | $0.704 \times 10^{-3}$ | $3.706 \times 10^{-3}$ | $3.165 \times 10^{-3}$ |
| $(1.0 \times 10^{-2}, 0.0 \times 10^{-2})$ | $0.031 \times 10^{-3}$ | $1.067 \times 10^{-3}$ | $2.919 \times 10^{-3}$ | $3.806 \times 10^{-3}$ |
| $(1.0 \times 10^{-2}, 0.2 \times 10^{-2})$ | $0.712 \times 10^{-3}$ | $0.221 \times 10^{-3}$ | $3.113 \times 10^{-3}$ | $3.234 \times 10^{-3}$ |
| $(1.0 \times 10^{-2}, 0.8 \times 10^{-2})$ | $-0.310 \times 10^{-3}$ | $0.692 \times 10^{-3}$ | $3.037 \times 10^{-3}$ | $2.835 \times 10^{-3}$ |

(a) The case of tightly linked loci with the recombination rate $r = 0.00001$.

| | Bias | | RMSE | |
|---|---|---|---|---|
| $(s_\mathcal{A}, s_\mathcal{B})$ | $s_\mathcal{A}$ | $s_\mathcal{B}$ | $s_\mathcal{A}$ | $s_\mathcal{B}$ |
| $(0.3 \times 10^{-2}, 0.0 \times 10^{-2})$ | $0.594 \times 10^{-3}$ | $1.818 \times 10^{-3}$ | $3.726 \times 10^{-3}$ | $4.455 \times 10^{-3}$ |
| $(0.3 \times 10^{-2}, 0.2 \times 10^{-2})$ | $1.230 \times 10^{-3}$ | $1.269 \times 10^{-3}$ | $4.231 \times 10^{-3}$ | $3.358 \times 10^{-3}$ |
| $(0.3 \times 10^{-2}, 0.8 \times 10^{-2})$ | $0.335 \times 10^{-3}$ | $0.473 \times 10^{-3}$ | $3.090 \times 10^{-3}$ | $3.002 \times 10^{-3}$ |
| $(1.0 \times 10^{-2}, 0.0 \times 10^{-2})$ | $0.497 \times 10^{-3}$ | $1.754 \times 10^{-3}$ | $2.539 \times 10^{-3}$ | $4.099 \times 10^{-3}$ |
| $(1.0 \times 10^{-2}, 0.2 \times 10^{-2})$ | $0.294 \times 10^{-3}$ | $1.071 \times 10^{-3}$ | $2.767 \times 10^{-3}$ | $3.978 \times 10^{-3}$ |
| $(1.0 \times 10^{-2}, 0.8 \times 10^{-2})$ | $0.447 \times 10^{-3}$ | $0.737 \times 10^{-3}$ | $2.729 \times 10^{-3}$ | $2.956 \times 10^{-3}$ |

(b) The case of loosely linked loci with the recombination rate $r = 0.01$.

Table S9: Bias and RMSE of the MAP estimates for 100 *haplotype frequency* datasets simulated with the initial population haplotype frequencies $\boldsymbol{x}_0 = (0.04, 0.08, 0.08, 0.8)$ and the dominance parameters $h_\mathcal{A} = 0.5$ and $h_\mathcal{B} = 0.5$.

| | Bias | | RMSE | |
|---|---|---|---|---|
| $(s_{\mathcal{A}}, s_{\mathcal{B}})$ | $s_{\mathcal{A}}$ | $s_{\mathcal{B}}$ | $s_{\mathcal{A}}$ | $s_{\mathcal{B}}$ |
| $(0.3 \times 10^{-2}, 0.0 \times 10^{-2})$ | $0.473 \times 10^{-3}$ | $0.062 \times 10^{-3}$ | $2.561 \times 10^{-3}$ | $2.766 \times 10^{-3}$ |
| $(0.3 \times 10^{-2}, 0.2 \times 10^{-2})$ | $0.229 \times 10^{-3}$ | $0.395 \times 10^{-3}$ | $2.487 \times 10^{-3}$ | $2.326 \times 10^{-3}$ |
| $(0.3 \times 10^{-2}, 0.8 \times 10^{-2})$ | $0.072 \times 10^{-3}$ | $0.154 \times 10^{-3}$ | $2.798 \times 10^{-3}$ | $2.492 \times 10^{-3}$ |
| $(1.0 \times 10^{-2}, 0.0 \times 10^{-2})$ | $-0.067 \times 10^{-3}$ | $0.207 \times 10^{-3}$ | $2.639 \times 10^{-3}$ | $2.549 \times 10^{-3}$ |
| $(1.0 \times 10^{-2}, 0.2 \times 10^{-2})$ | $-0.435 \times 10^{-3}$ | $-0.202 \times 10^{-3}$ | $2.732 \times 10^{-3}$ | $2.441 \times 10^{-3}$ |
| $(1.0 \times 10^{-2}, 0.8 \times 10^{-2})$ | $0.065 \times 10^{-3}$ | $-0.147 \times 10^{-3}$ | $2.794 \times 10^{-3}$ | $2.540 \times 10^{-3}$ |

(a) The case of tightly linked loci with the recombination rate $r = 0.00001$.

| | Bias | | RMSE | |
|---|---|---|---|---|
| $(s_{\mathcal{A}}, s_{\mathcal{B}})$ | $s_{\mathcal{A}}$ | $s_{\mathcal{B}}$ | $s_{\mathcal{A}}$ | $s_{\mathcal{B}}$ |
| $(0.3 \times 10^{-2}, 0.0 \times 10^{-2})$ | $-0.125 \times 10^{-3}$ | $0.095 \times 10^{-3}$ | $2.565 \times 10^{-3}$ | $2.393 \times 10^{-3}$ |
| $(0.3 \times 10^{-2}, 0.2 \times 10^{-2})$ | $0.591 \times 10^{-3}$ | $-0.195 \times 10^{-3}$ | $2.504 \times 10^{-3}$ | $2.647 \times 10^{-3}$ |
| $(0.3 \times 10^{-2}, 0.8 \times 10^{-2})$ | $0.022 \times 10^{-3}$ | $-0.056 \times 10^{-3}$ | $2.598 \times 10^{-3}$ | $2.691 \times 10^{-3}$ |
| $(1.0 \times 10^{-2}, 0.0 \times 10^{-2})$ | $0.375 \times 10^{-3}$ | $0.194 \times 10^{-3}$ | $2.684 \times 10^{-3}$ | $2.222 \times 10^{-3}$ |
| $(1.0 \times 10^{-2}, 0.2 \times 10^{-2})$ | $-0.021 \times 10^{-3}$ | $0.168 \times 10^{-3}$ | $2.818 \times 10^{-3}$ | $2.449 \times 10^{-3}$ |
| $(1.0 \times 10^{-2}, 0.8 \times 10^{-2})$ | $0.195 \times 10^{-3}$ | $0.064 \times 10^{-3}$ | $2.418 \times 10^{-3}$ | $2.397 \times 10^{-3}$ |

(b) The case of loosely linked loci with the recombination rate $r = 0.01$.

Table S10: Bias and RMSE of the MAP estimates for 100 *haplotype frequency* datasets simulated with the initial population haplotype frequencies $\boldsymbol{x}_0 = (0.1, 0.2, 0.3, 0.4)$ and the dominance parameters $h_{\mathcal{A}} = 0.5$ and $h_{\mathcal{B}} = 0.5$.

| | Bias | | RMSE | |
|---|---|---|---|---|
| $(s_\mathcal{A}, s_\mathcal{B})$ | $s_\mathcal{A}$ | $s_\mathcal{B}$ | $s_\mathcal{A}$ | $s_\mathcal{B}$ |
| $(0.3 \times 10^{-2}, 0.0 \times 10^{-2})$ | $-0.269 \times 10^{-3}$ | $0.259 \times 10^{-3}$ | $4.287 \times 10^{-3}$ | $2.899 \times 10^{-3}$ |
| $(0.3 \times 10^{-2}, 0.2 \times 10^{-2})$ | $0.465 \times 10^{-3}$ | $0.051 \times 10^{-3}$ | $4.322 \times 10^{-3}$ | $2.466 \times 10^{-3}$ |
| $(0.3 \times 10^{-2}, 0.8 \times 10^{-2})$ | $1.164 \times 10^{-3}$ | $-0.022 \times 10^{-3}$ | $3.554 \times 10^{-3}$ | $2.359 \times 10^{-3}$ |
| $(1.0 \times 10^{-2}, 0.0 \times 10^{-2})$ | $0.124 \times 10^{-3}$ | $0.542 \times 10^{-3}$ | $3.638 \times 10^{-3}$ | $2.776 \times 10^{-3}$ |
| $(1.0 \times 10^{-2}, 0.2 \times 10^{-2})$ | $-0.512 \times 10^{-3}$ | $0.597 \times 10^{-3}$ | $3.938 \times 10^{-3}$ | $2.725 \times 10^{-3}$ |
| $(1.0 \times 10^{-2}, 0.8 \times 10^{-2})$ | $0.075 \times 10^{-3}$ | $-0.131 \times 10^{-3}$ | $3.521 \times 10^{-3}$ | $3.153 \times 10^{-3}$ |

(a) The case of tightly linked loci with the recombination rate $r = 0.00001$.

| | Bias | | RMSE | |
|---|---|---|---|---|
| $(s_\mathcal{A}, s_\mathcal{B})$ | $s_\mathcal{A}$ | $s_\mathcal{B}$ | $s_\mathcal{A}$ | $s_\mathcal{B}$ |
| $(0.3 \times 10^{-2}, 0.0 \times 10^{-2})$ | $0.301 \times 10^{-3}$ | $0.563 \times 10^{-3}$ | $3.118 \times 10^{-3}$ | $2.457 \times 10^{-3}$ |
| $(0.3 \times 10^{-2}, 0.2 \times 10^{-2})$ | $0.375 \times 10^{-3}$ | $0.052 \times 10^{-3}$ | $3.889 \times 10^{-3}$ | $2.112 \times 10^{-3}$ |
| $(0.3 \times 10^{-2}, 0.8 \times 10^{-2})$ | $0.204 \times 10^{-3}$ | $-0.624 \times 10^{-3}$ | $3.426 \times 10^{-3}$ | $2.019 \times 10^{-3}$ |
| $(1.0 \times 10^{-2}, 0.0 \times 10^{-2})$ | $0.202 \times 10^{-3}$ | $1.242 \times 10^{-3}$ | $3.174 \times 10^{-3}$ | $3.164 \times 10^{-3}$ |
| $(1.0 \times 10^{-2}, 0.2 \times 10^{-2})$ | $0.472 \times 10^{-3}$ | $0.522 \times 10^{-3}$ | $3.370 \times 10^{-3}$ | $2.147 \times 10^{-3}$ |
| $(1.0 \times 10^{-2}, 0.8 \times 10^{-2})$ | $0.538 \times 10^{-3}$ | $-0.199 \times 10^{-3}$ | $3.233 \times 10^{-3}$ | $2.374 \times 10^{-3}$ |

(b) The case of loosely linked loci with the recombination rate $r = 0.01$.

Table S11: Bias and RMSE of the MAP estimates for 100 *haplotype frequency* datasets simulated with the initial population haplotype frequencies $\boldsymbol{x}_0 = (0.0001, 0, 0.1999, 0.8)$ and the dominance parameters $h_\mathcal{A} = 0.5$ and $h_\mathcal{B} = 0.5$. It should be noted that in this case we condition the mutant alleles at both loci to survive until the most recent sampling time point and sample 50 chromosomes from the underlying population at every 120 generations throughout 1200 generations.

| | Bias | | RMSE | |
|---|---|---|---|---|
| $(s_{\mathcal{A}}, s_{\mathcal{B}})$ | $s_{\mathcal{A}}$ | $s_{\mathcal{B}}$ | $s_{\mathcal{A}}$ | $s_{\mathcal{B}}$ |
| $(0.3 \times 10^{-2}, 0.0 \times 10^{-2})$ | $0.034 \times 10^{-3}$ | $0.351 \times 10^{-3}$ | $2.328 \times 10^{-3}$ | $3.134 \times 10^{-3}$ |
| $(0.3 \times 10^{-2}, 0.2 \times 10^{-2})$ | $-0.113 \times 10^{-3}$ | $1.277 \times 10^{-3}$ | $2.095 \times 10^{-3}$ | $3.562 \times 10^{-3}$ |
| $(0.3 \times 10^{-2}, 0.8 \times 10^{-2})$ | $0.034 \times 10^{-3}$ | $-0.042 \times 10^{-3}$ | $2.165 \times 10^{-3}$ | $2.319 \times 10^{-3}$ |
| $(1.0 \times 10^{-2}, 0.0 \times 10^{-2})$ | $-0.434 \times 10^{-3}$ | $0.778 \times 10^{-3}$ | $3.046 \times 10^{-3}$ | $3.626 \times 10^{-3}$ |
| $(1.0 \times 10^{-2}, 0.2 \times 10^{-2})$ | $-0.006 \times 10^{-3}$ | $0.860 \times 10^{-3}$ | $2.446 \times 10^{-3}$ | $3.109 \times 10^{-3}$ |
| $(1.0 \times 10^{-2}, 0.8 \times 10^{-2})$ | $-0.310 \times 10^{-3}$ | $0.036 \times 10^{-3}$ | $2.463 \times 10^{-3}$ | $2.397 \times 10^{-3}$ |

(a) The case of tightly linked loci with the recombination rate $r = 0.00001$.

| | Bias | | RMSE | |
|---|---|---|---|---|
| $(s_{\mathcal{A}}, s_{\mathcal{B}})$ | $s_{\mathcal{A}}$ | $s_{\mathcal{B}}$ | $s_{\mathcal{A}}$ | $s_{\mathcal{B}}$ |
| $(0.3 \times 10^{-2}, 0.0 \times 10^{-2})$ | $-0.130 \times 10^{-3}$ | $1.537 \times 10^{-3}$ | $2.009 \times 10^{-3}$ | $3.920 \times 10^{-3}$ |
| $(0.3 \times 10^{-2}, 0.2 \times 10^{-2})$ | $-0.170 \times 10^{-3}$ | $0.790 \times 10^{-3}$ | $2.074 \times 10^{-3}$ | $2.892 \times 10^{-3}$ |
| $(0.3 \times 10^{-2}, 0.8 \times 10^{-2})$ | $-0.332 \times 10^{-3}$ | $0.229 \times 10^{-3}$ | $2.157 \times 10^{-3}$ | $2.437 \times 10^{-3}$ |
| $(1.0 \times 10^{-2}, 0.0 \times 10^{-2})$ | $-0.376 \times 10^{-3}$ | $1.141 \times 10^{-3}$ | $3.009 \times 10^{-3}$ | $3.897 \times 10^{-3}$ |
| $(1.0 \times 10^{-2}, 0.2 \times 10^{-2})$ | $-0.269 \times 10^{-3}$ | $0.617 \times 10^{-3}$ | $2.605 \times 10^{-3}$ | $2.869 \times 10^{-3}$ |
| $(1.0 \times 10^{-2}, 0.8 \times 10^{-2})$ | $-0.755 \times 10^{-3}$ | $-0.179 \times 10^{-3}$ | $3.258 \times 10^{-3}$ | $2.332 \times 10^{-3}$ |

(b) The case of loosely linked loci with the recombination rate $r = 0.01$.

Table S12: Bias and RMSE of the MAP estimates for 100 *haplotype frequency* datasets simulated with the initial population haplotype frequencies $\boldsymbol{x}_0 = (0.1, 0.2, 0.3, 0.4)$ and the dominance parameters $h_{\mathcal{A}} = 0$ and $h_{\mathcal{B}} = 1$.

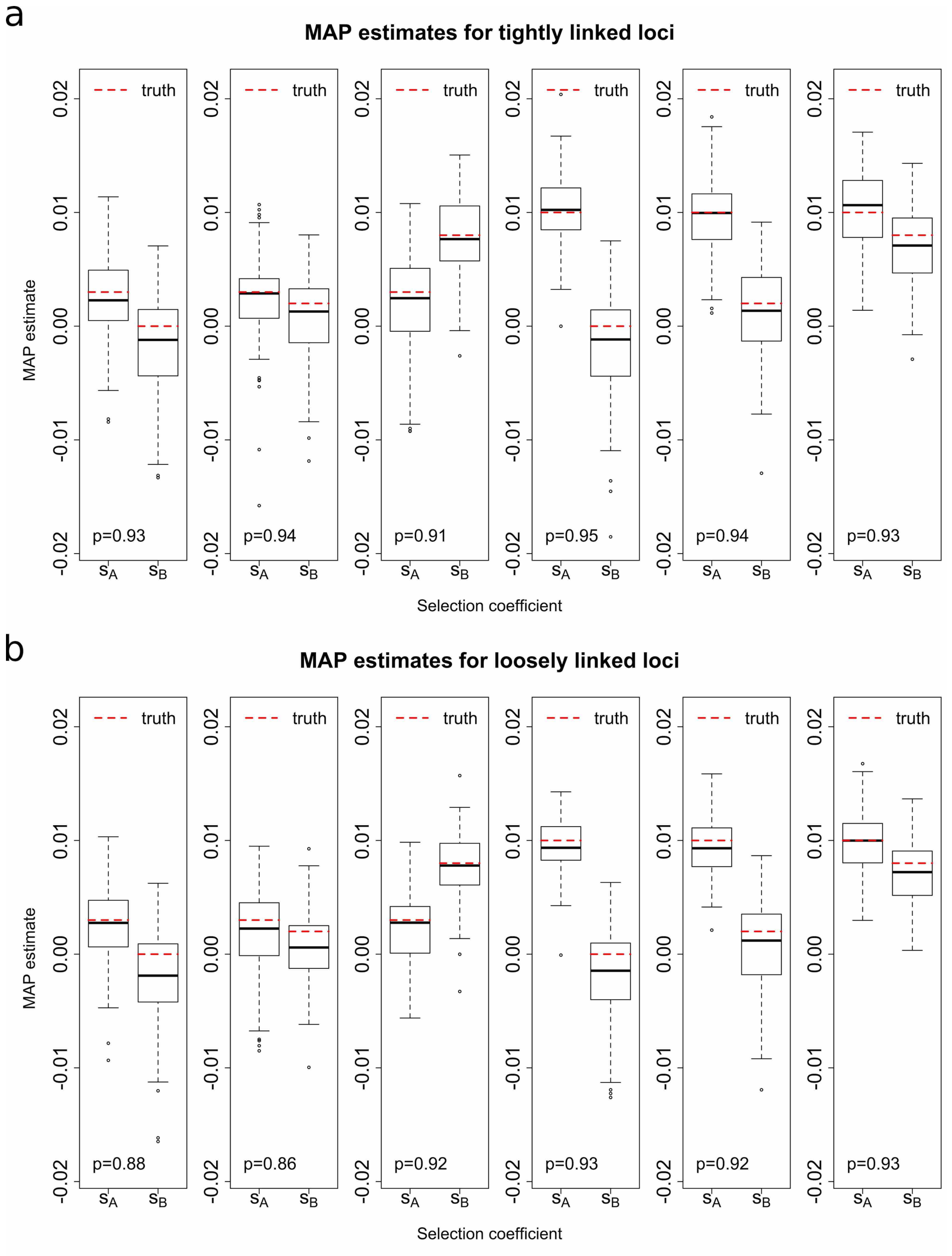


Figure S7: Empirical distributions of the MAP estimates for 100 *allele frequency* datasets (*without* missing values) simulated with the initial population haplotype frequencies $\boldsymbol{x}_0 = (0.04, 0.08, 0.08, 0.8)$ and the dominance parameters $h_{\mathcal{A}} = 0.5$ and $h_{\mathcal{B}} = 0.5$ for the case of (a) tightly linked loci with the recombination rate $r = 0.00001$ and (b) loosely linked loci with the recombination rate $r = 0.01$. The $p$ value in the bottom left corner indicates the proportion of the runs where the true values of the selection coefficients both fall within their 95% HPD intervals.

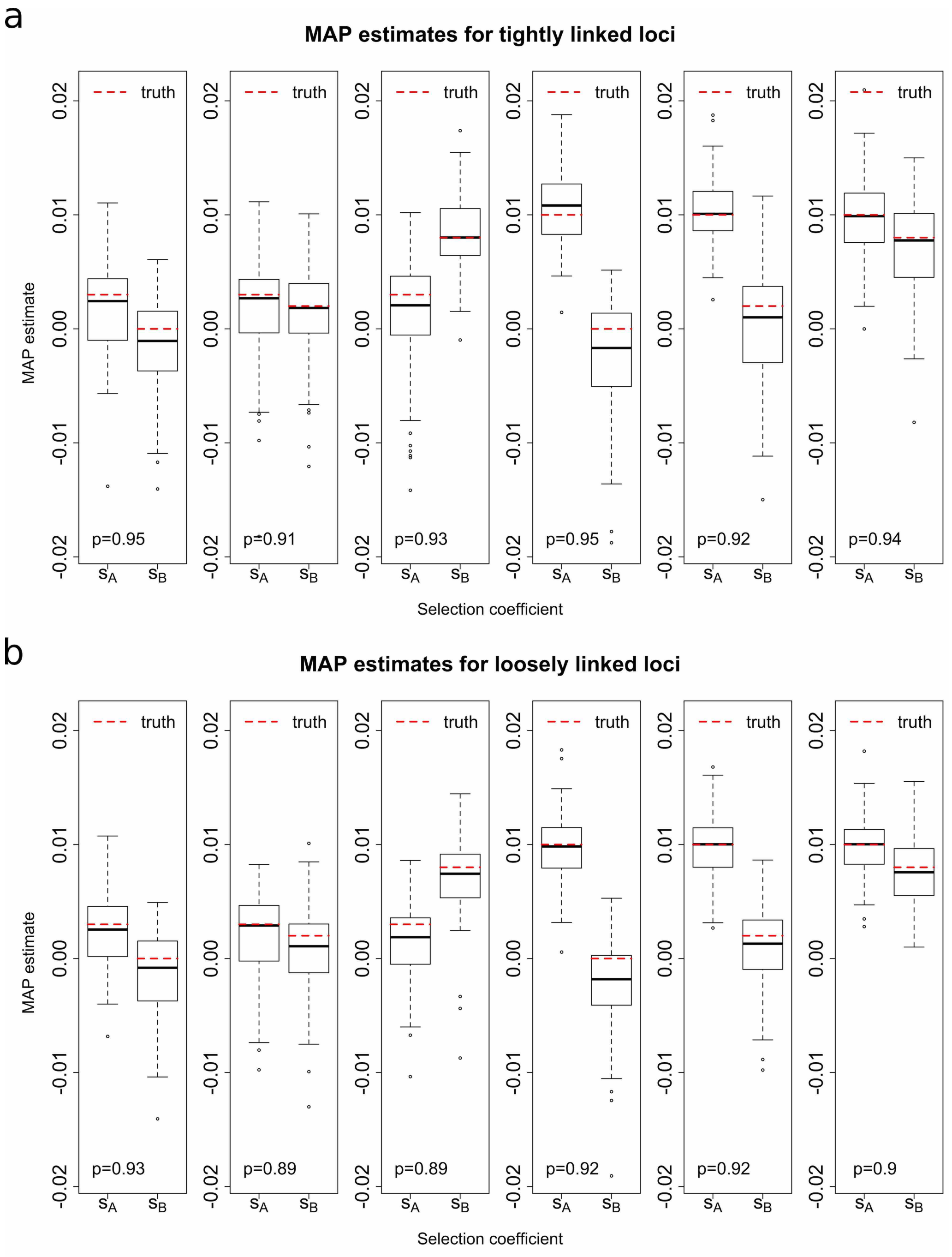


Figure S8: Empirical distributions of the MAP estimates for 100 *allele frequency* datasets (*with* 2% missing values) simulated with the initial population haplotype frequencies $\boldsymbol{x}_0 = (0.04, 0.08, 0.08, 0.8)$ and the dominance parameters $h_\mathcal{A} = 0.5$ and $h_\mathcal{B} = 0.5$ for the case of (a) tightly linked loci with the recombination rate $r = 0.00001$ and (b) loosely linked loci with the recombination rate $r = 0.01$. The $p$ value in the bottom left corner indicates the proportion of the runs where the true values of the selection coefficients both fall within their 95% HPD intervals.

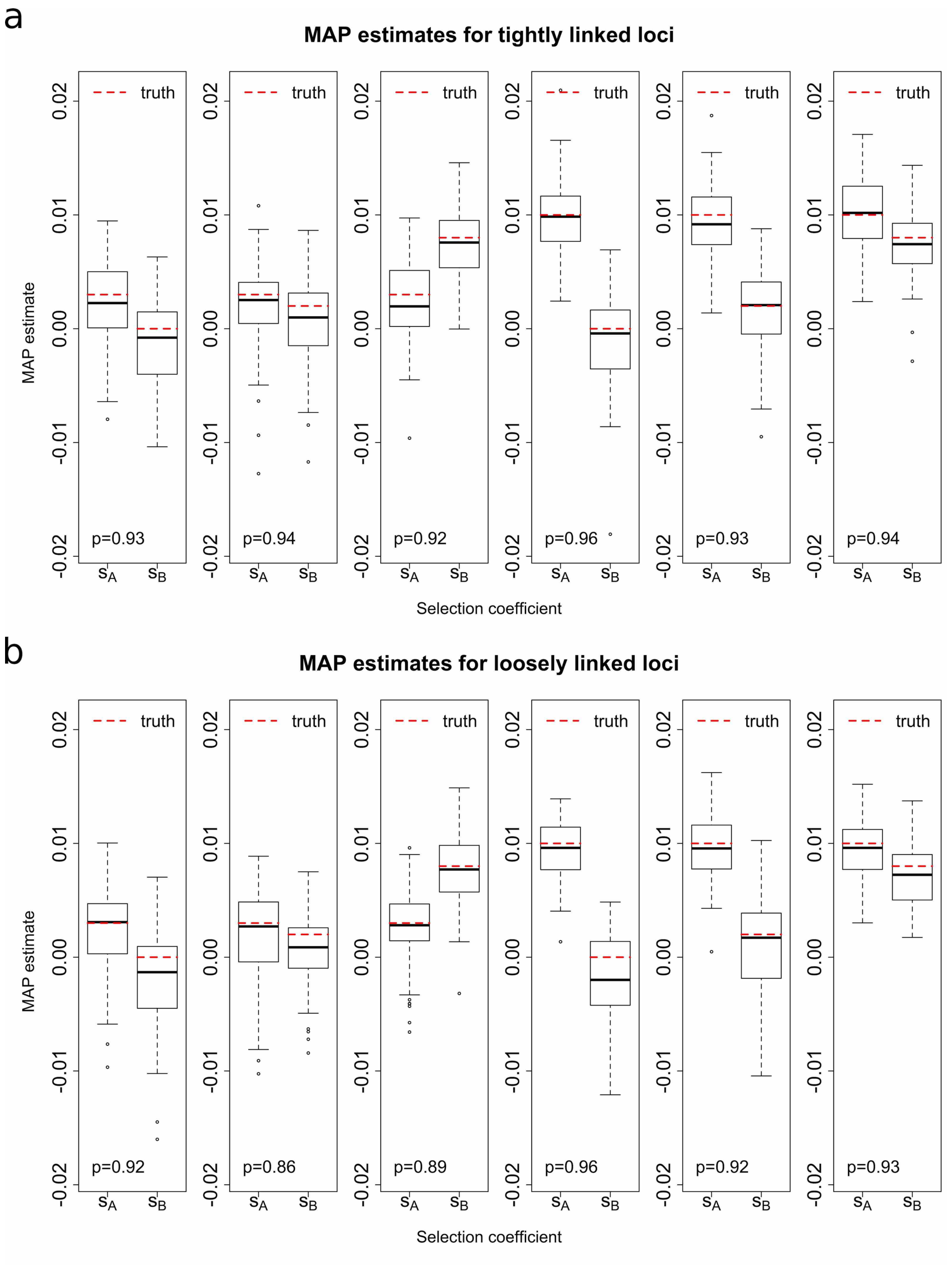


Figure S9: Empirical distributions of the MAP estimates for 100 *haplotype frequency* datasets simulated with the initial population haplotype frequencies $\boldsymbol{x}_0 = (0.04, 0.08, 0.08, 0.8)$ and the dominance parameters $h_{\mathcal{A}} = 0.5$ and $h_{\mathcal{B}} = 0.5$ for the case of (a) tightly linked loci with the recombination rate $r = 0.00001$ and (b) loosely linked loci with the recombination rate $r = 0.01$. The $p$ value in the bottom left corner indicates the proportion of the runs where the true values of the selection coefficients both fall within their 95% HPD intervals.

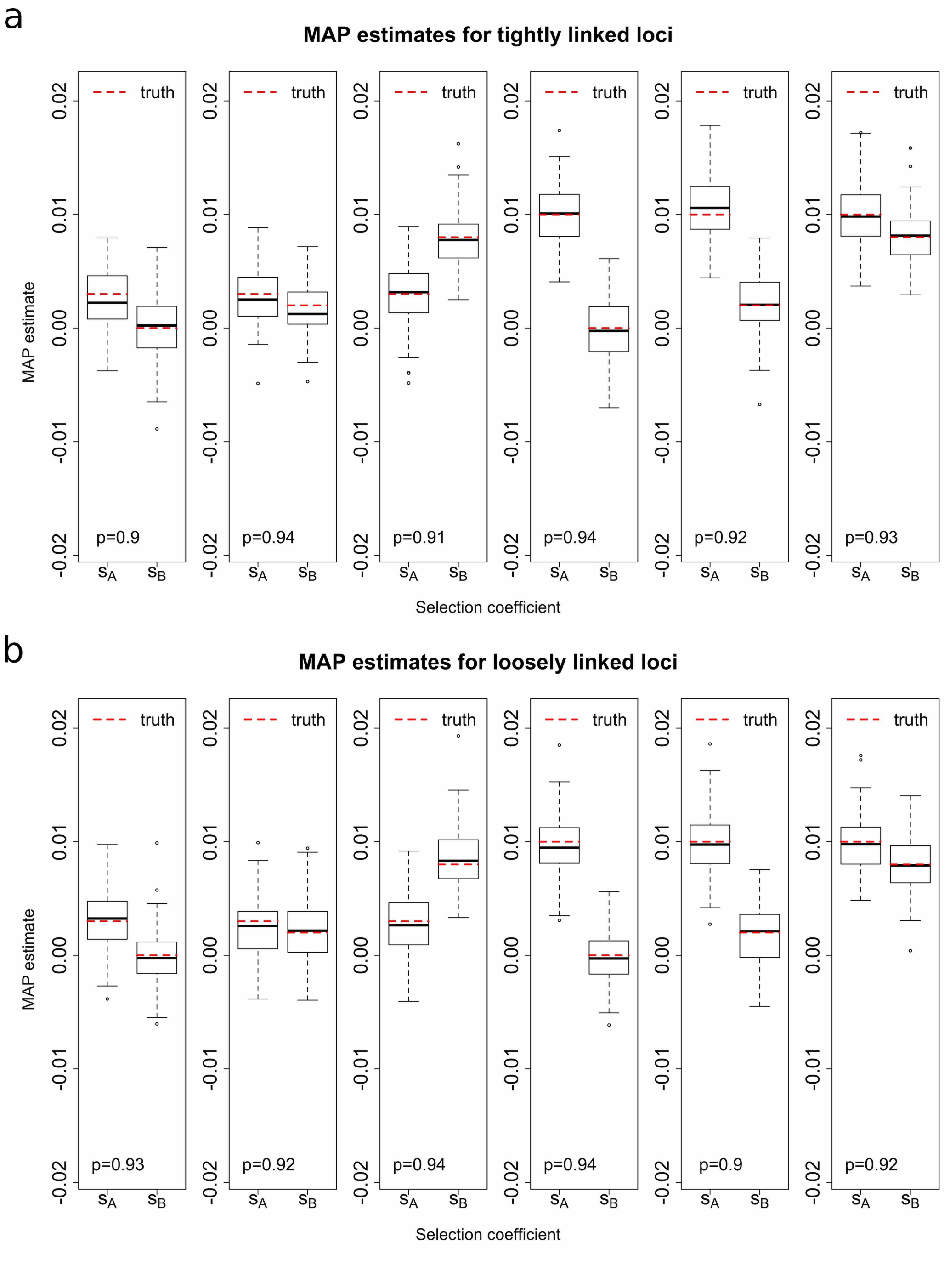


Figure S10: Empirical distributions of the MAP estimates for 100 *haplotype frequency* datasets simulated with the initial population haplotype frequencies $\boldsymbol{x}_0 = (0.1, 0.2, 0.3, 0.4)$ and the dominance parameters $h_{\mathcal{A}} = 0.5$ and $h_{\mathcal{B}} = 0.5$ for the case of (a) tightly linked loci with the recombination rate $r = 0.00001$ and (b) loosely linked loci with the recombination rate $r = 0.01$. The $p$ value in the bottom left corner indicates the proportion of the runs where the true values of the selection coefficients both fall within their 95% HPD intervals.

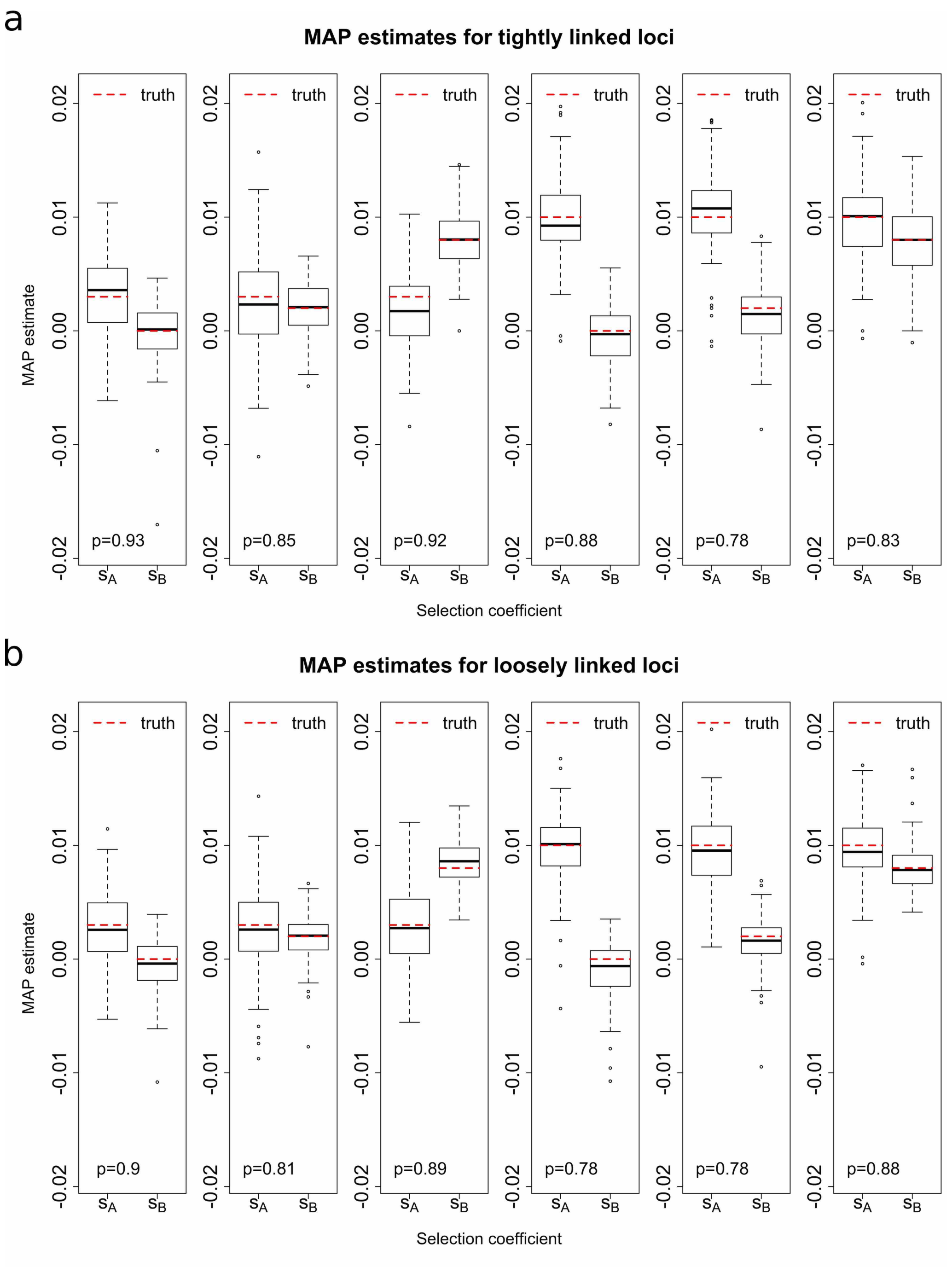


Figure S11: Empirical distributions of the MAP estimates for 100 *haplotype frequency* datasets simulated with the initial population haplotype frequencies $\boldsymbol{x}_0 = (0.0001, 0, 0.1999, 0.8)$ and the dominance parameters $h_{\mathcal{A}} = 0.5$ and $h_{\mathcal{B}} = 0.5$ for the case of (a) tightly linked loci with the recombination rate $r = 0.00001$ and (b) loosely linked loci with the recombination rate $r = 0.01$. The $p$ value in the bottom left corner indicates the proportion of the runs where the true values of the selection coefficients both fall within their 95% HPD intervals. It should be noted that in this case we condition the mutant alleles at both loci to survive until the most recent sampling time point and sample 50 chromosomes from the underlying population at every 120 generations throughout 1200 generations.

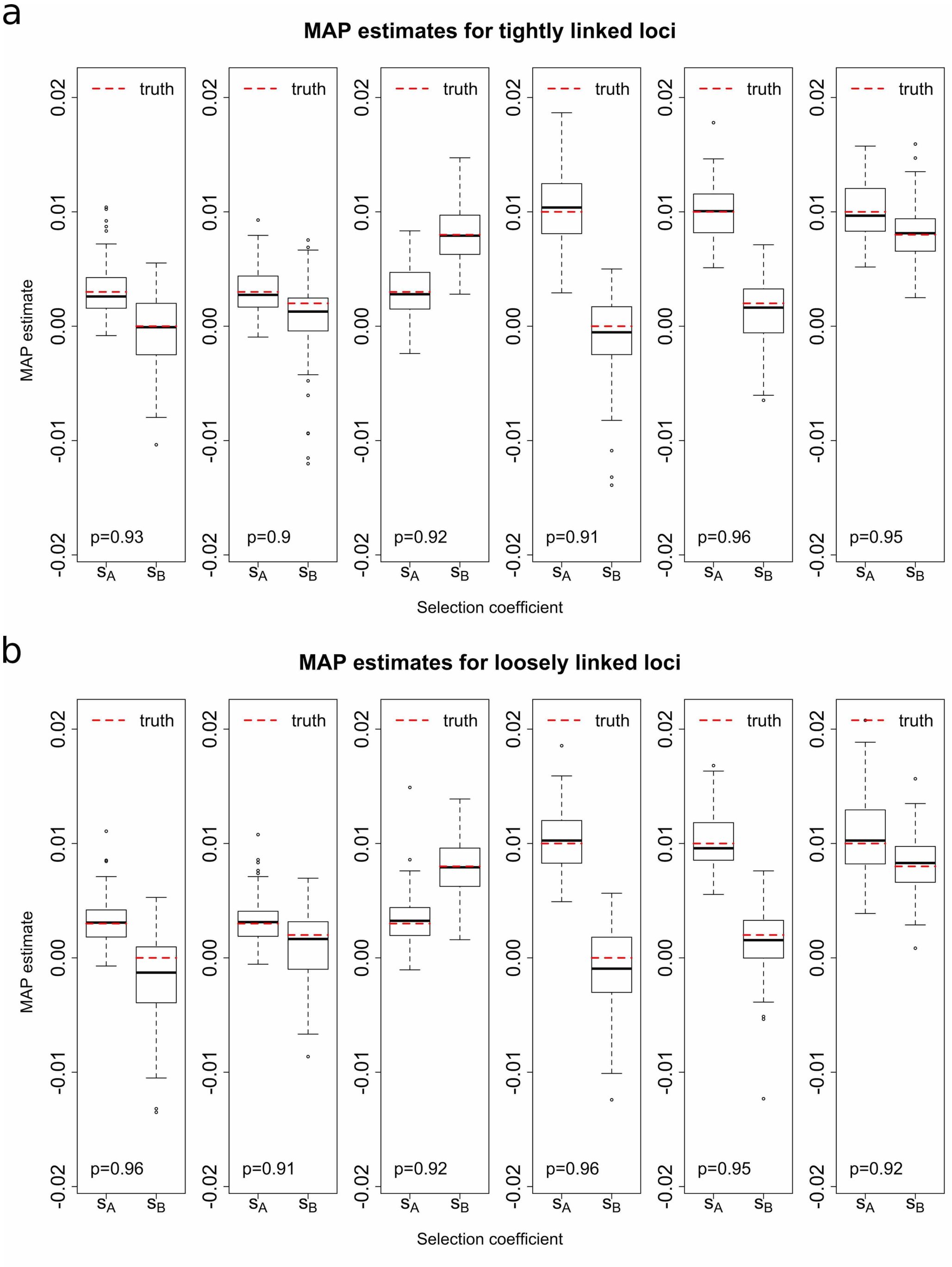


Figure S12: Empirical distributions of the MAP estimates for 100 *haplotype frequency* datasets simulated with the initial population haplotype frequencies $\boldsymbol{x}_0 = (0.1, 0.2, 0.3, 0.4)$ and the dominance parameters $h_{\mathcal{A}} = 0$ and $h_{\mathcal{B}} = 1$ for the case of (a) tightly linked loci with the recombination rate $r = 0.00001$ and (b) loosely linked loci with the recombination rate $r = 0.01$. The $p$ value in the bottom left corner indicates the proportion of the runs where the true values of the selection coefficients both fall within their 95% HPD intervals.

### File S2. Additional results for the analysis of real data

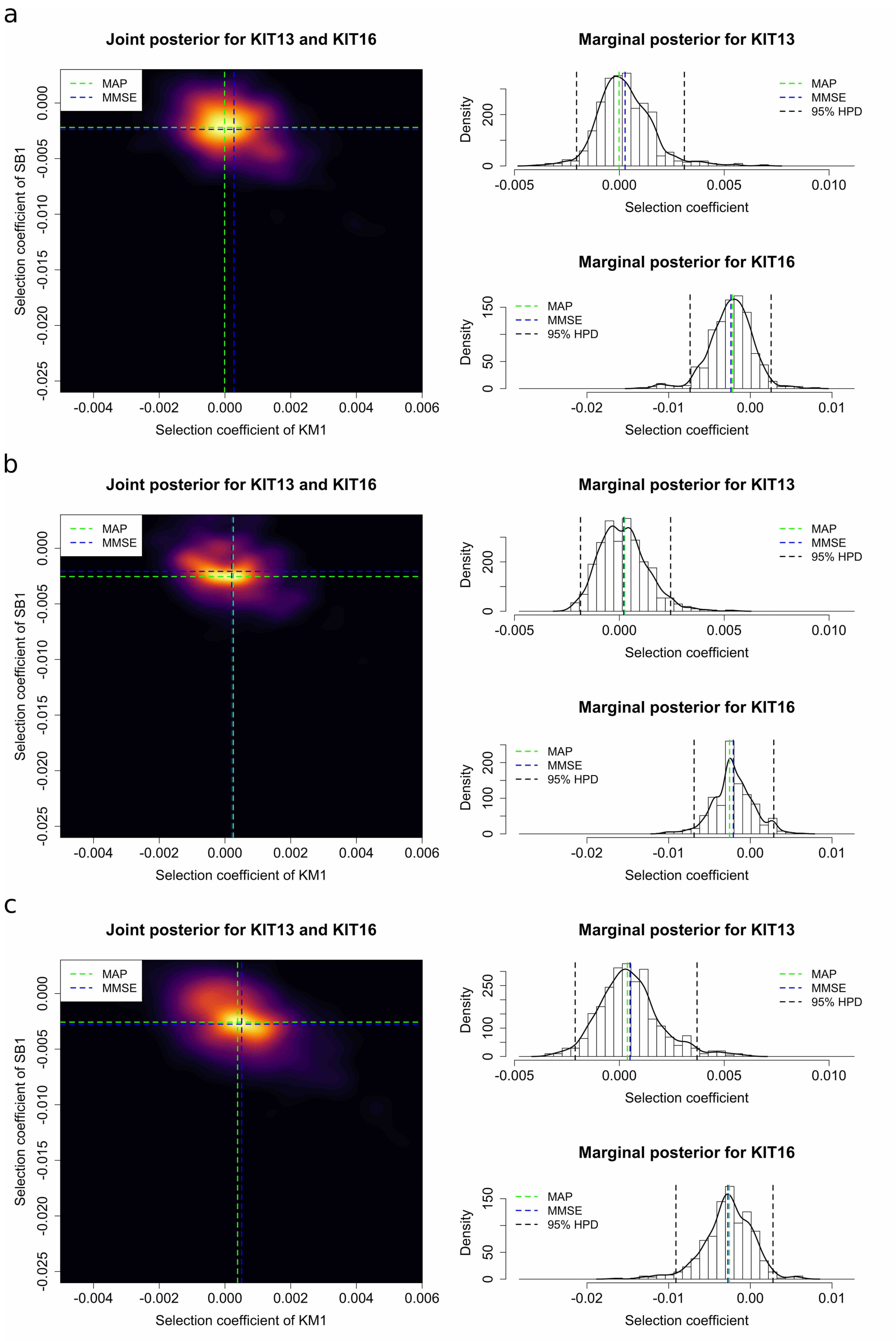


Figure S13: Posterior probability distributions for *KIT13* and *KIT16* obtained by using the two-locus method from the samples dated from 17146 years BP (the first sampling time point) with the population size of 8000 and the average rate of recombination (a) $5 \times 10^{-9}$ crossovers/bp, (b) $1 \times 10^{-8}$ crossovers/bp and (c) $5 \times 10^{-8}$ crossovers/bp.

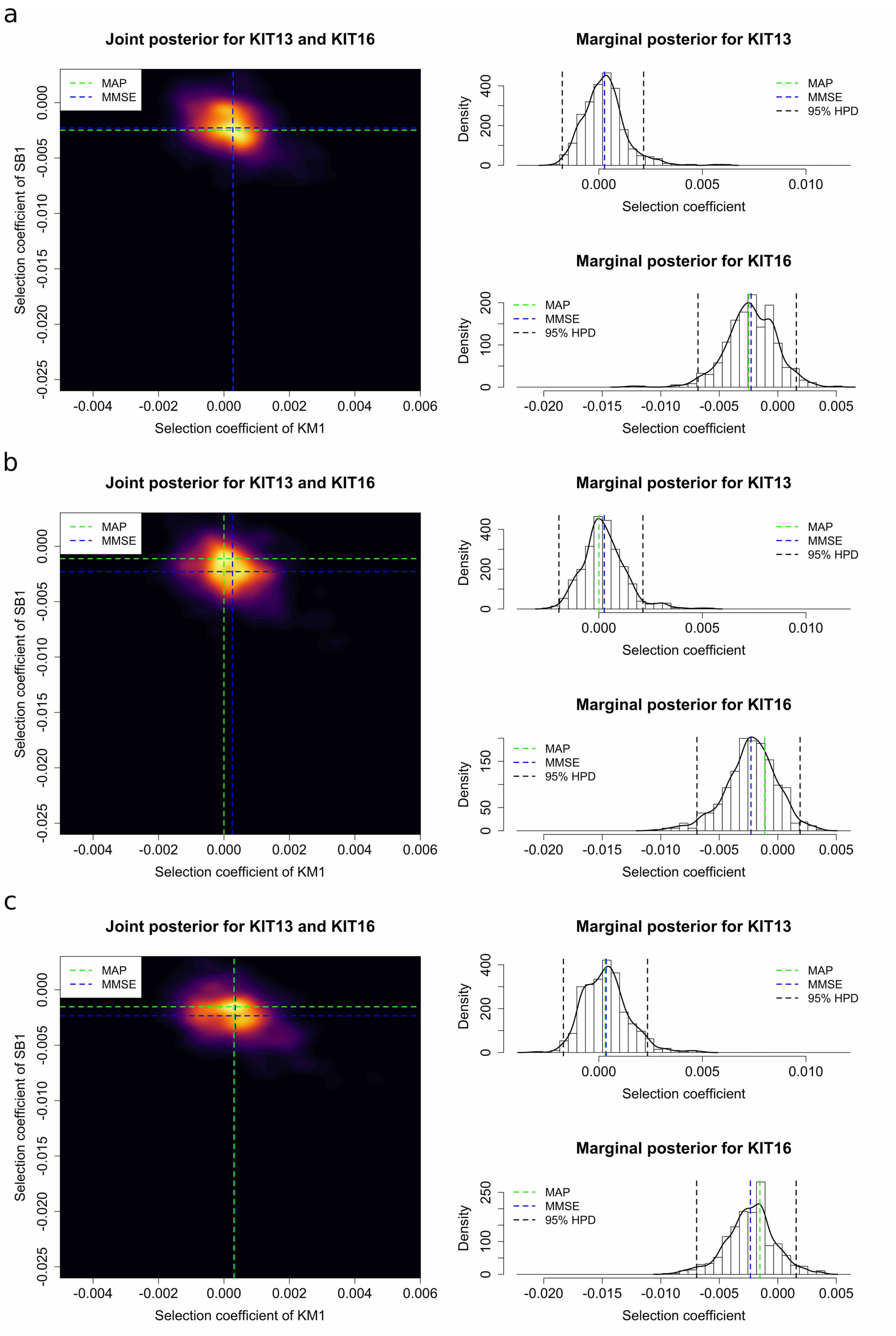


Figure S14: Posterior probability distributions for *KIT13* and *KIT16* obtained by using the two-locus method from the samples dated from 17146 years BP (the first sampling time point) with the population size of 16000 and the average rate of recombination (a) $5 \times 10^{-9}$ crossovers/bp, (b) $1 \times 10^{-8}$ crossovers/bp and (c) $5 \times 10^{-8}$ crossovers/bp.

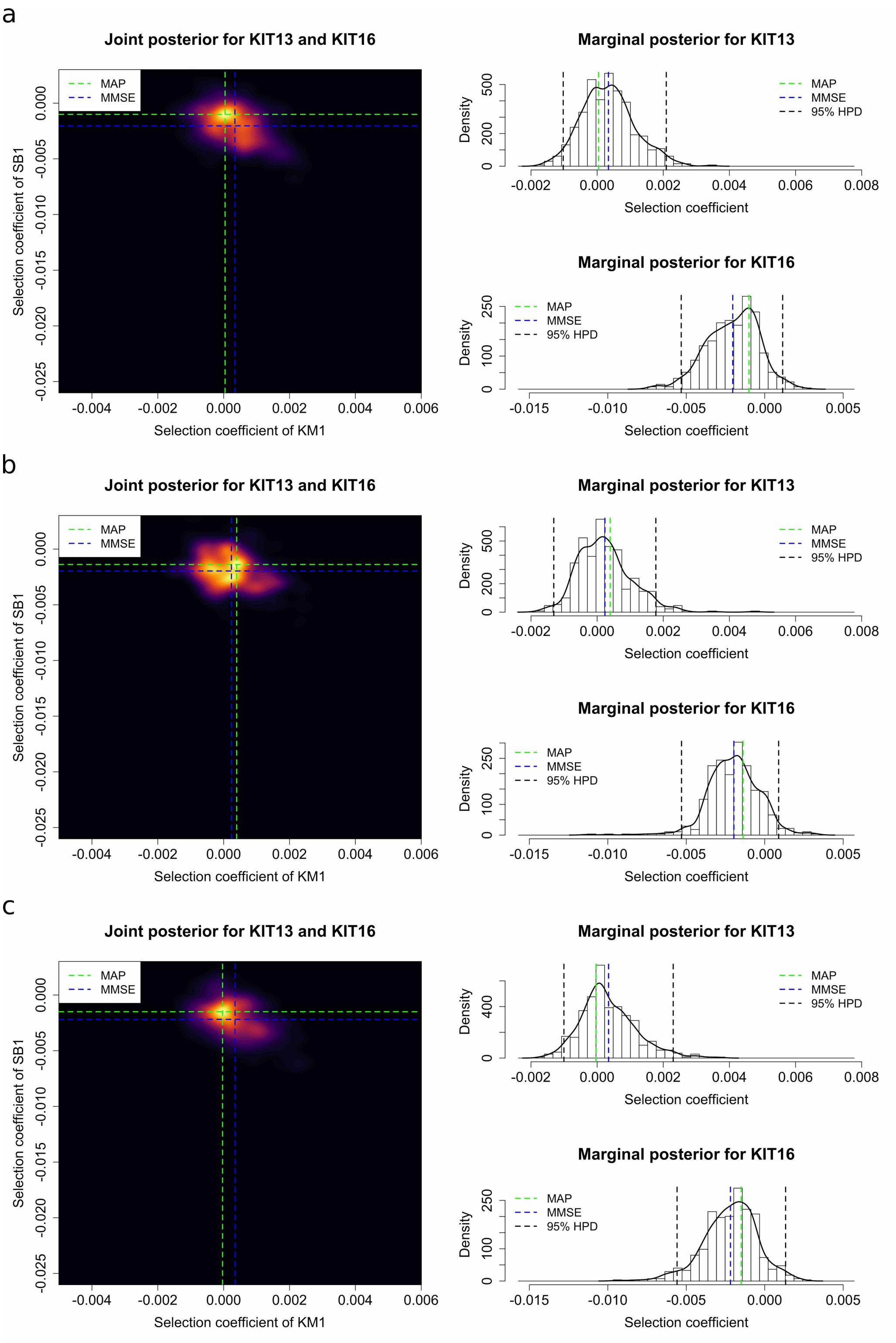


Figure S15: Posterior probability distributions for *KIT13* and *KIT16* obtained by using the two-locus method from the samples dated from 17146 years BP (the first sampling time point) with the population size of 32000 and the average rate of recombination (a) $5 \times 10^{-9}$ crossovers/bp, (b) $1 \times 10^{-8}$ crossovers/bp and (c) $5 \times 10^{-8}$ crossovers/bp.

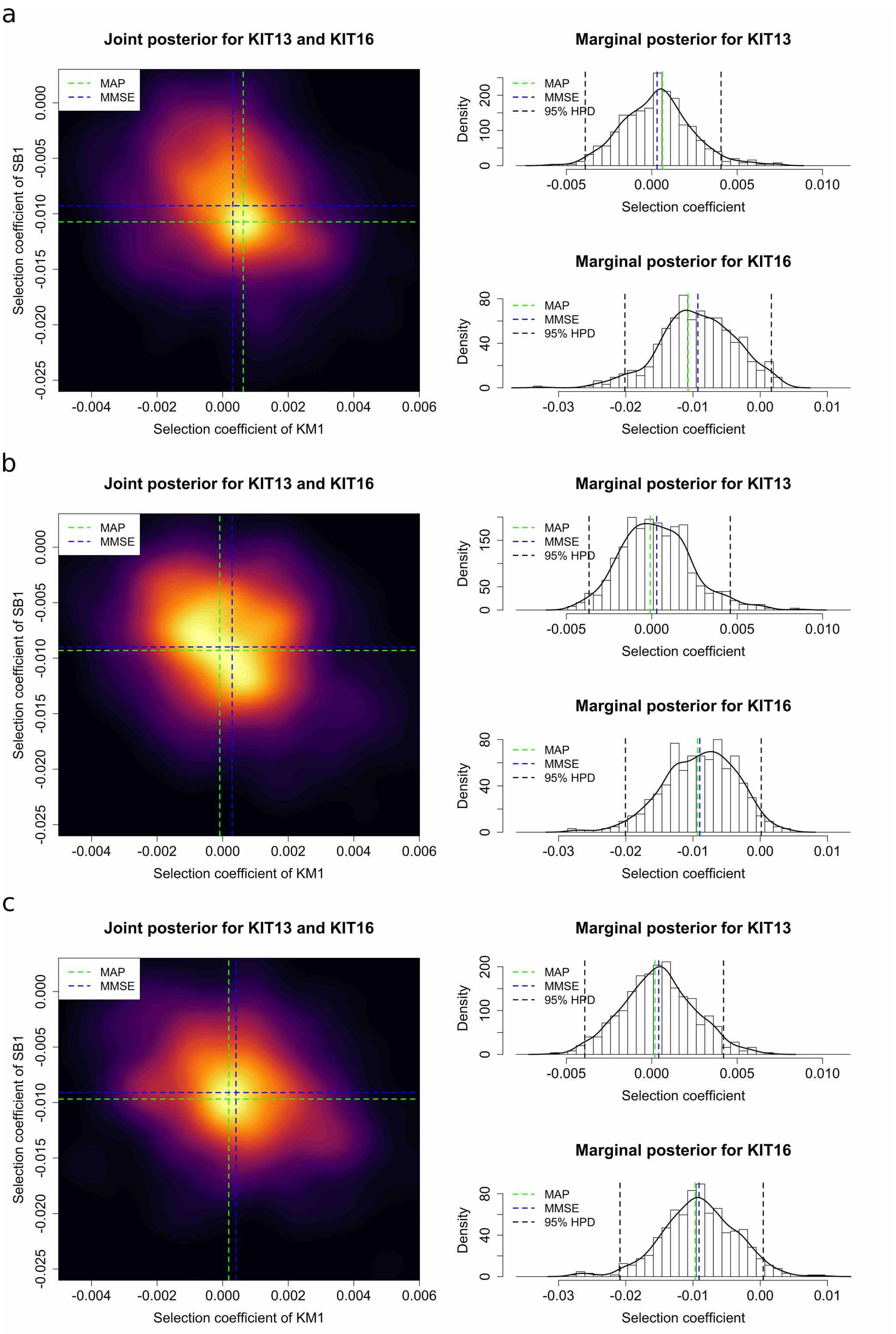


Figure S16: Posterior probability distributions for *KIT13* and *KIT16* obtained by using the two-locus method from the samples dated from 7029 years BP (the second sampling time point) with the population size of 8000 and the average rate of recombination (a) $5 \times 10^{-9}$ crossovers/bp, (b) $1 \times 10^{-8}$ crossovers/bp and (c) $5 \times 10^{-8}$ crossovers/bp.

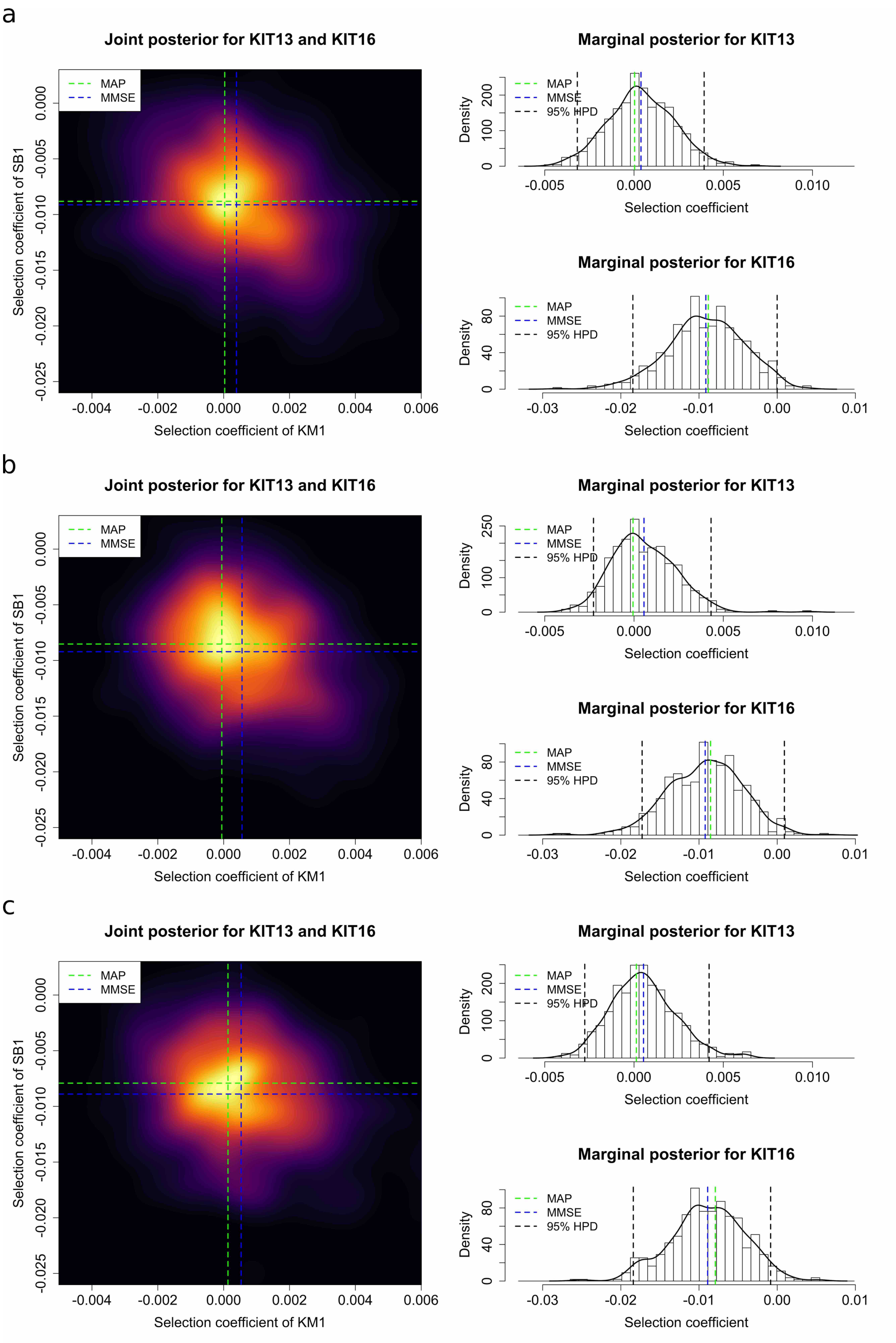


Figure S17: Posterior probability distributions for *KIT13* and *KIT16* obtained by using the two-locus method from the samples dated from 7029 years BP (the second sampling time point) with the population size of 16000 and the average rate of recombination (a) $5 \times 10^{-9}$ crossovers/bp, (b) $1 \times 10^{-8}$ crossovers/bp and (c) $5 \times 10^{-8}$ crossovers/bp.

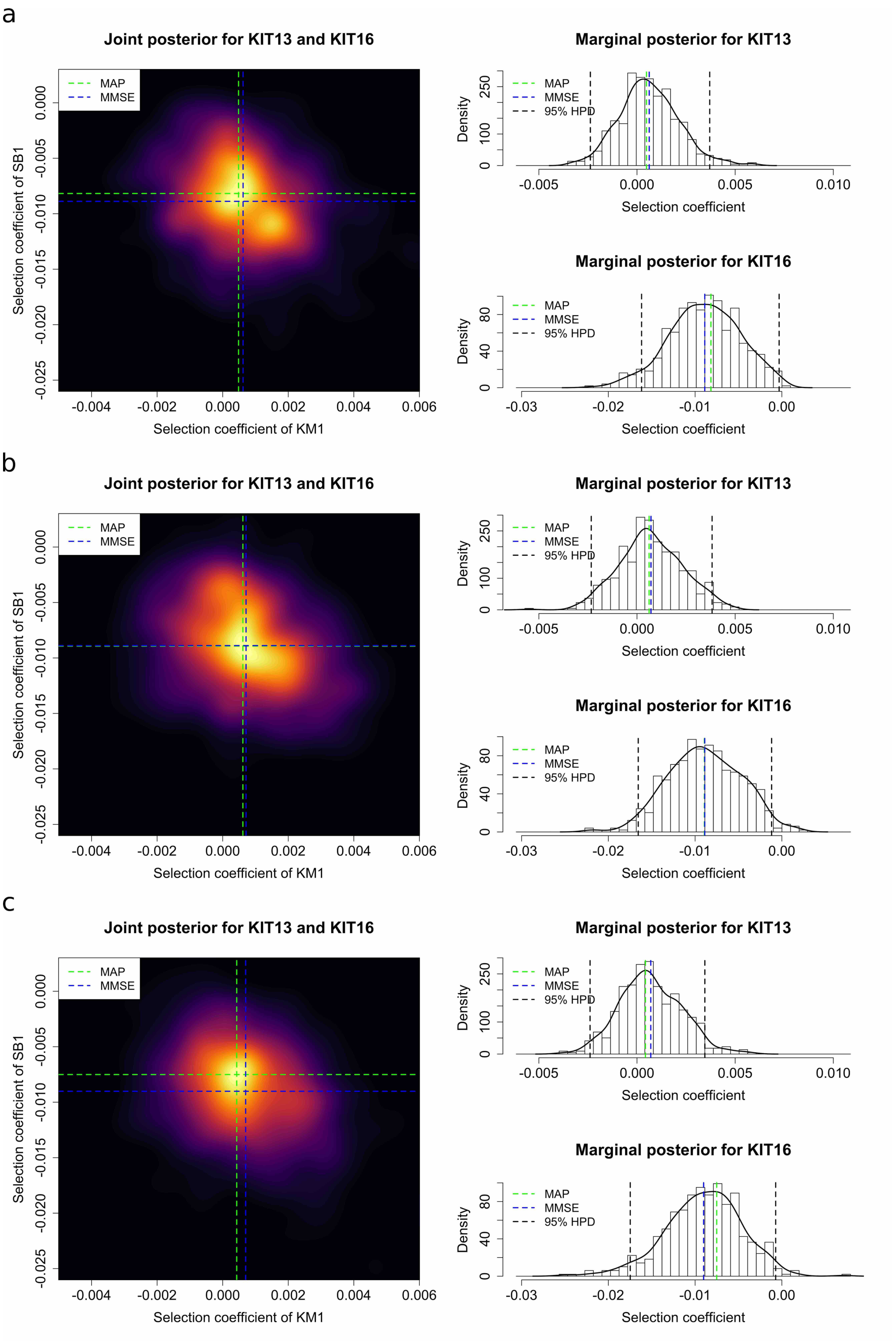


Figure S18: Posterior probability distributions for *KIT13* and *KIT16* obtained by using the two-locus method from the samples dated from 7029 years BP (the second sampling time point) with the population size of 32000 and the average rate of recombination (a) $5 \times 10^{-9}$ crossovers/bp, (b) $1 \times 10^{-8}$ crossovers/bp and (c) $5 \times 10^{-8}$ crossovers/bp.

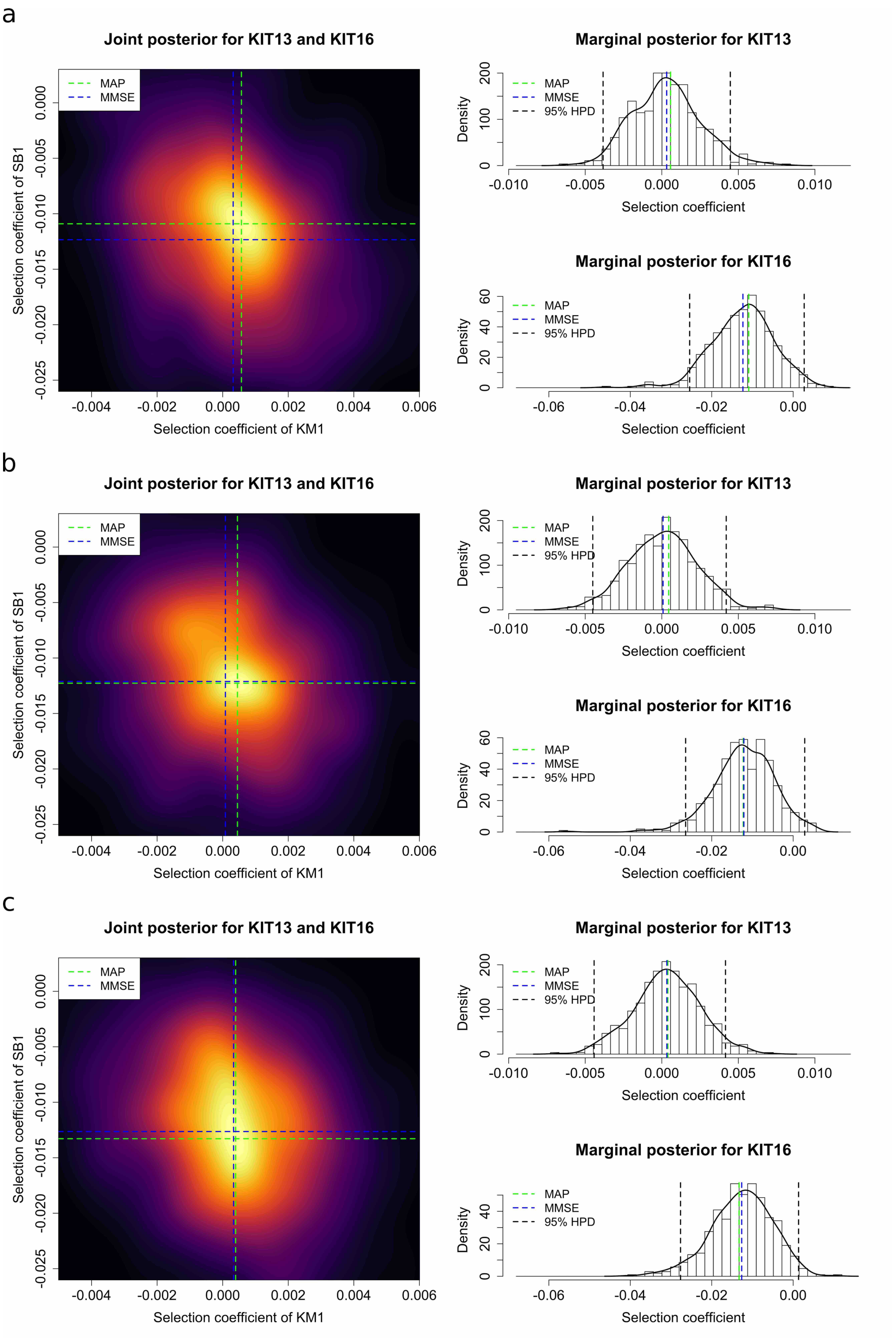


Figure S19: Posterior probability distributions for *KIT13* and *KIT16* obtained by using the two-locus method from the samples dated from 5472 years BP (the third sampling time point) with the population size of 8000 and the average rate of recombination (a) $5 \times 10^{-9}$ crossovers/bp, (b) $1 \times 10^{-8}$ crossovers/bp and (c) $5 \times 10^{-8}$ crossovers/bp.

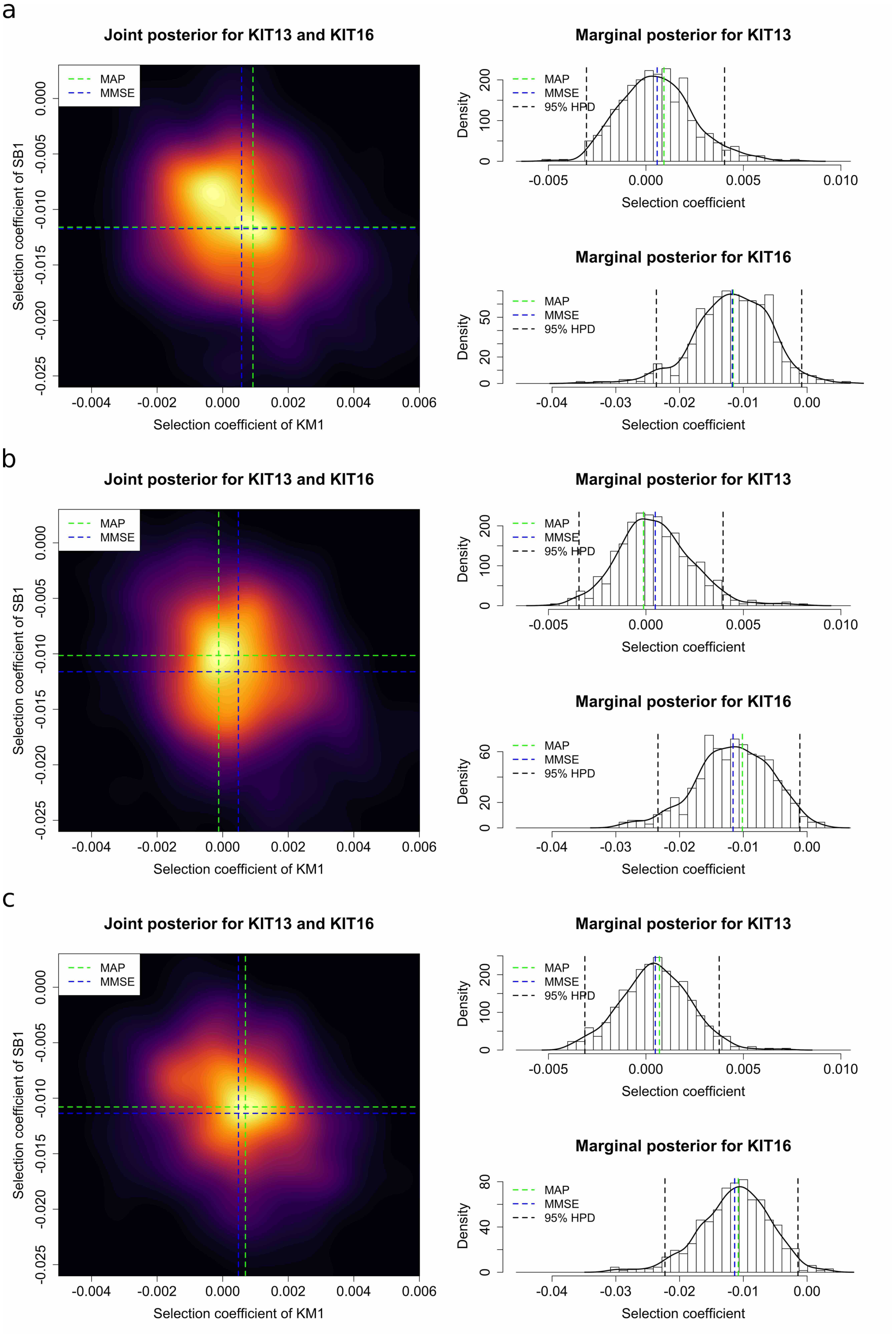


Figure S20: Posterior probability distributions for *KIT13* and *KIT16* obtained by using the two-locus method from the samples dated from 5472 years BP (the third sampling time point) with the population size of 32000 and the average rate of recombination (a) $5 \times 10^{-9}$ crossovers/bp, (b) $1 \times 10^{-8}$ crossovers/bp and (c) $5 \times 10^{-8}$ crossovers/bp.

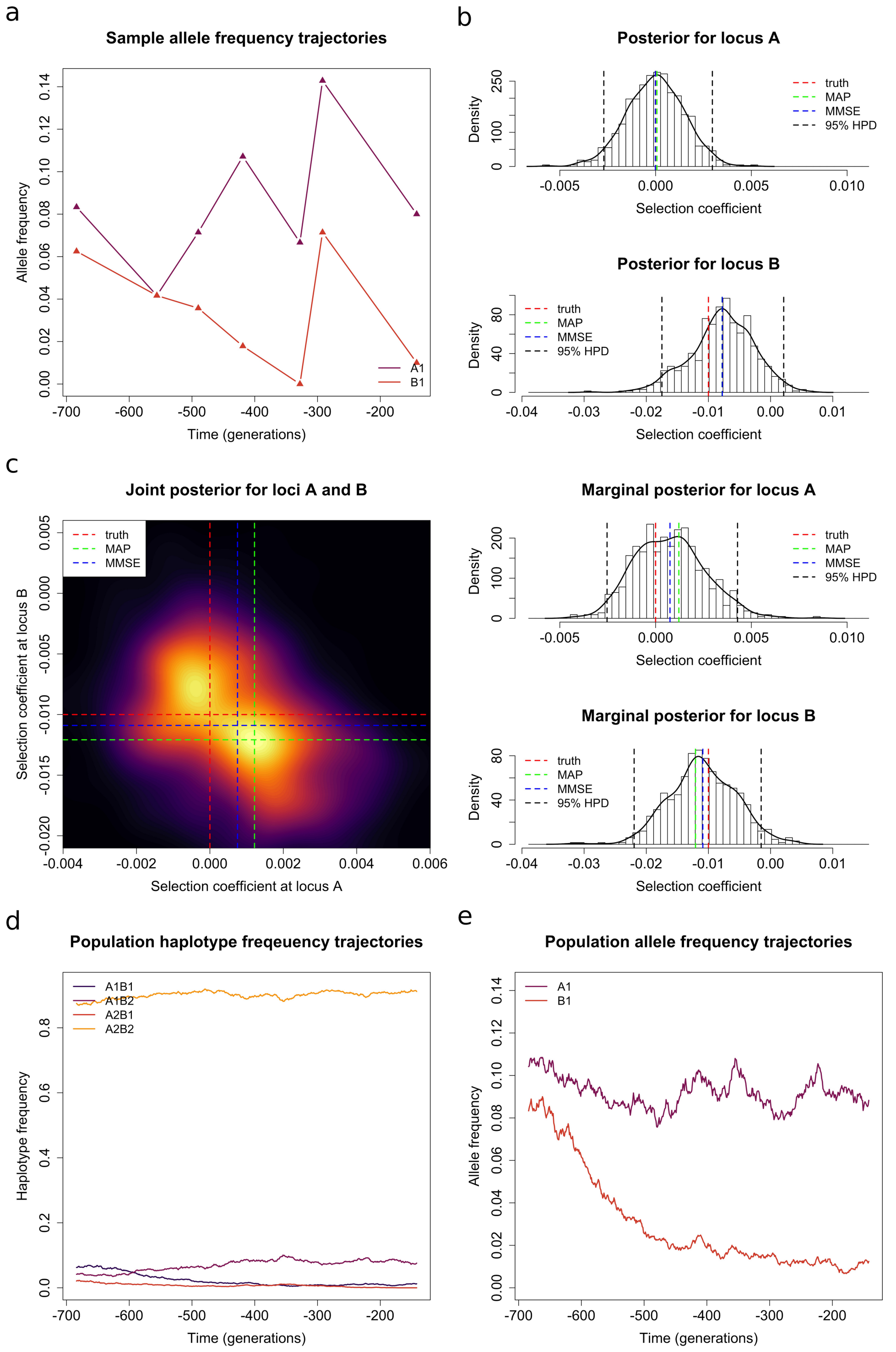


Figure S21: A comparison of the performance differences of the single-locus method and the two-locus method on the simulated dataset of a negatively selected locus tightly linked with a selectively neutral locus. (a) Sample mutant allele frequency trajectories. (b) Posteriors obtained with a single-locus method. (c) Posteriors obtained with a two-locus method. (d) Population mutant allele frequency trajectories. (e) Population haplotype frequency trajectories. It should be noted that in this case we simulate the haplotype frequency trajectories of the underlying population with the initial population haplotype frequencies $\boldsymbol{x}_0 = (1/16, 1/24, 1/48, 7/8)$, where the selection coefficients $s_{\mathcal{A}} = 0$ and $s_{\mathcal{B}} = -0.01$, the dominance parameters $h_{\mathcal{A}} = 0$ and $h_{\mathcal{B}} = 0.5$, the recombination rate $r = 0.469 \times 10^{-4}$ and the population size $N = 16000$. Sampling times and sizes are the same as the *KIT* dataset but without unknown alleles.

| | | single-locus method | two-locus method |
|---|---|---|---|
| selection coefficient $s_{\mathcal{A}}$ | MAP ($\times 10^{-2}$) | 0.007 | 0.122 |
| | MMSE ($\times 10^{-2}$) | $-0.001$ | 0.075 |
| | 95% HPD ($\times 10^{-2}$) | $[-0.270, 0.297]$ | $[-0.253, \quad 0.428]$ |
| selection coefficient $s_{\mathcal{B}}$ | MAP ($\times 10^{-2}$) | $-0.780$ | $-1.208$ |
| | MMSE ($\times 10^{-2}$) | $-0.777$ | $-1.089$ |
| | 95% HPD ($\times 10^{-2}$) | $[-1.748, 0.211]$ | $[-2.195, -0.151]$ |

Table S13: A comparison of the Bayesian estimates obtained by using the single-locus method and the two-locus method from the simulated dataset of a negatively selected locus tightly linked with a selectively neutral locus. It should be noted that in this case we simulate the haplotype frequency trajectories of the underlying population with the initial population haplotype frequencies $\boldsymbol{x}_0 = (1/16, 1/24, 1/48, 7/8)$, where the selection coefficients $s_{\mathcal{A}} = 0$ and $s_{\mathcal{B}} = -0.01$, the dominance parameters $h_{\mathcal{A}} = 0$ and $h_{\mathcal{B}} = 0.5$, the recombination rate $r = 0.469 \times 10^{-4}$ and the population size $N = 16000$. Sampling times and sizes are the same as the *KIT* dataset but without unknown alleles.

| | recombination rate | MAP ($\times 10^{-2}$) | MMSE ($\times 10^{-2}$) | 95% HPD ($\times 10^{-2}$) |
|---|---|---|---|---|
| *KIT13* | $0.234 \times 10^{-4}$ | -0.001 | 0.027 | $[-0.204, \ 0.310]$ |
| | $0.469 \times 10^{-4}$ | 0.025 | 0.020 | $[-0.185, \ 0.244]$ |
| | $2.340 \times 10^{-4}$ | 0.038 | 0.051 | $[-0.211, \ 0.370]$ |
| *KIT16* | $0.234 \times 10^{-4}$ | -0.220 | -0.237 | $[-0.738, \ 0.254]$ |
| | $0.469 \times 10^{-4}$ | -0.254 | -0.208 | $[-0.688, \ 0.287]$ |
| | $2.340 \times 10^{-4}$ | -0.257 | -0.275 | $[-0.911, \ 0.278]$ |

(a) Resulting estimates obtained with the population size of 8000.

| | recombination rate | MAP ($\times 10^{-2}$) | MMSE ($\times 10^{-2}$) | 95% HPD ($\times 10^{-2}$) |
|---|---|---|---|---|
| *KIT13* | $0.234 \times 10^{-4}$ | 0.027 | 0.027 | $[-0.177, \ 0.216]$ |
| | $0.469 \times 10^{-4}$ | 0.000 | 0.026 | $[-0.193, \ 0.213]$ |
| | $2.340 \times 10^{-4}$ | 0.031 | 0.035 | $[-0.171, \ 0.235]$ |
| *KIT16* | $0.234 \times 10^{-4}$ | -0.249 | -0.228 | $[-0.682, \ 0.158]$ |
| | $0.469 \times 10^{-4}$ | -0.112 | -0.229 | $[-0.691, \ 0.191]$ |
| | $2.340 \times 10^{-4}$ | -0.153 | -0.234 | $[-0.694, \ 0.157]$ |

(b) Resulting estimates obtained with the population size of 16000.

| | recombination rate | MAP ($\times 10^{-2}$) | MMSE ($\times 10^{-2}$) | 95% HPD ($\times 10^{-2}$) |
|---|---|---|---|---|
| *KIT13* | $0.234 \times 10^{-4}$ | 0.004 | 0.034 | $[-0.103, \ 0.209]$ |
| | $0.469 \times 10^{-4}$ | 0.040 | 0.024 | $[-0.131, \ 0.178]$ |
| | $2.340 \times 10^{-4}$ | -0.003 | 0.035 | $[-0.100, \ 0.230]$ |
| *KIT16* | $0.234 \times 10^{-4}$ | -0.101 | -0.204 | $[-0.532, \ 0.114]$ |
| | $0.469 \times 10^{-4}$ | -0.138 | -0.197 | $[-0.530, \ 0.089]$ |
| | $2.340 \times 10^{-4}$ | -0.150 | -0.218 | $[-0.558, \ 0.133]$ |

(c) Resulting estimates obtained with the population size of 32000.

Table S14: MAP and MMSE estimates, as well as the 95% HPD intervals, for *KIT13* and *KIT16* obtained from the samples dated from 17146 years BP (the first sampling time point).

| | recombination rate | MAP ($\times 10^{-2}$) | MMSE ($\times 10^{-2}$) | 95% HPD ($\times 10^{-2}$) |
|---|---|---|---|---|
| *KIT13* | $0.234 \times 10^{-4}$ | 0.063 | 0.031 | $[-0.389, \ 0.406]$ |
| | $0.469 \times 10^{-4}$ | -0.009 | 0.029 | $[-0.367, \ 0.460]$ |
| | $2.340 \times 10^{-4}$ | 0.019 | 0.041 | $[-0.392, \ 0.420]$ |
| *KIT16* | $0.234 \times 10^{-4}$ | -1.073 | -0.928 | $[-2.013, \ 0.166]$ |
| | $0.469 \times 10^{-4}$ | -0.931 | -0.899 | $[-2.005, \ 0.015]$ |
| | $2.340 \times 10^{-4}$ | -0.969 | -0.910 | $[-2.086, \ 0.045]$ |

(a) Resulting estimates obtained with the population size of 8000.

| | recombination rate | MAP ($\times 10^{-2}$) | MMSE ($\times 10^{-2}$) | 95% HPD ($\times 10^{-2}$) |
|---|---|---|---|---|
| *KIT13* | $0.234 \times 10^{-4}$ | 0.003 | 0.039 | $[-0.318, \ 0.393]$ |
| | $0.469 \times 10^{-4}$ | -0.006 | 0.056 | $[-0.227, \ 0.432]$ |
| | $2.340 \times 10^{-4}$ | 0.013 | 0.053 | $[-0.276, \ 0.421]$ |
| *KIT16* | $0.234 \times 10^{-4}$ | -0.881 | -0.913 | $[-1.846, -0.001]$ |
| | $0.469 \times 10^{-4}$ | -0.852 | -0.921 | $[-1.728, \ 0.092]$ |
| | $2.340 \times 10^{-4}$ | -0.791 | -0.889 | $[-1.841, -0.084]$ |

(b) Resulting estimates obtained with the population size of 16000.

| | recombination rate | MAP ($\times 10^{-2}$) | MMSE ($\times 10^{-2}$) | 95% HPD ($\times 10^{-2}$) |
|---|---|---|---|---|
| *KIT13* | $0.234 \times 10^{-4}$ | 0.048 | 0.062 | $[-0.238, \ 0.370]$ |
| | $0.469 \times 10^{-4}$ | 0.062 | 0.071 | $[-0.233, \ 0.383]$ |
| | $2.340 \times 10^{-4}$ | 0.043 | 0.070 | $[-0.239, \ 0.346]$ |
| *KIT16* | $0.234 \times 10^{-4}$ | -0.818 | -0.889 | $[-1.617, -0.031]$ |
| | $0.469 \times 10^{-4}$ | -0.893 | -0.889 | $[-1.656, -0.118]$ |
| | $2.340 \times 10^{-4}$ | -0.751 | -0.902 | $[-1.748, -0.071]$ |

(c) Resulting estimates obtained with the population size of 32000.

Table S15: MAP and MMSE estimates, as well as the 95% HPD intervals, for *KIT13* and *KIT16* obtained from the samples dated from 7029 years BP (the second sampling time point).

| | recombination rate | MAP ($\times 10^{-2}$) | MMSE ($\times 10^{-2}$) | 95% HPD ($\times 10^{-2}$) |
|---|---|---|---|---|
| *KIT13* | $0.234 \times 10^{-4}$ | 0.057 | 0.032 | $[-0.384, \ 0.447]$ |
| | $0.469 \times 10^{-4}$ | 0.045 | 0.008 | $[-0.451, \ 0.421]$ |
| | $2.340 \times 10^{-4}$ | 0.040 | 0.033 | $[-0.442, \ 0.416]$ |
| *KIT16* | $0.234 \times 10^{-4}$ | -1.090 | -1.234 | $[-2.542, \ 0.271]$ |
| | $0.469 \times 10^{-4}$ | -1.227 | -1.212 | $[-2.641, \ 0.284]$ |
| | $2.340 \times 10^{-4}$ | -1.329 | -1.264 | $[-2.766, \ 0.133]$ |

(a) Resulting estimates obtained with the population size of 8000.

| | recombination rate | MAP ($\times 10^{-2}$) | MMSE ($\times 10^{-2}$) | 95% HPD ($\times 10^{-2}$) |
|---|---|---|---|---|
| *KIT13* | $0.234 \times 10^{-4}$ | 0.079 | 0.056 | $[-0.268, \ 0.476]$ |
| | $0.469 \times 10^{-4}$ | -0.021 | 0.037 | $[-0.292, \ 0.451]$ |
| | $2.340 \times 10^{-4}$ | 0.036 | 0.040 | $[-0.283, \ 0.447]$ |
| *KIT16* | $0.234 \times 10^{-4}$ | -1.238 | -1.175 | $[-2.316, \ 0.250]$ |
| | $0.469 \times 10^{-4}$ | -1.076 | -1.187 | $[-2.407, \ 0.007]$ |
| | $2.340 \times 10^{-4}$ | -1.001 | -1.152 | $[-2.283, \ 0.002]$ |

(b) Resulting estimates obtained with the population size of 16000.

| | recombination rate | MAP ($\times 10^{-2}$) | MMSE ($\times 10^{-2}$) | 95% HPD ($\times 10^{-2}$) |
|---|---|---|---|---|
| *KIT13* | $0.234 \times 10^{-4}$ | 0.092 | 0.058 | $[-0.305, \ 0.403]$ |
| | $0.469 \times 10^{-4}$ | -0.012 | 0.048 | $[-0.343, \ 0.396]$ |
| | $2.340 \times 10^{-4}$ | 0.069 | 0.048 | $[-0.313, \ 0.376]$ |
| *KIT16* | $0.234 \times 10^{-4}$ | -1.160 | -1.173 | $[-2.364, -0.084]$ |
| | $0.469 \times 10^{-4}$ | -1.015 | -1.161 | $[-2.339, -0.113]$ |
| | $2.340 \times 10^{-4}$ | -1.079 | -1.136 | $[-2.229, -0.143]$ |

(c) Resulting estimates obtained with the population size of 32000.

Table S16: MAP and MMSE estimates, as well as the 95% HPD intervals, for *KIT13* and *KIT16* obtained from the samples dated from 5472 years BP (the third sampling time point).

#### File S3. Detecting and quantifying natural selection at a single locus

We employ a Bayesian statistical framework to infer natural selection at a single locus from allele frequency time series data. Similar to Section 2.2, we use an HMM framework, where the underlying population is assumed to evolve according to the one-locus Wright-Fisher diffusion with selection

$$dX(t) = \alpha X(t)(1 - X(t))\left((1 - h) - (1 - 2h)X(t)\right) \, dt + \sqrt{X(t)(1 - X(t))} \, dW(t) \tag{1}$$

for $t \geq t_0$, with initial condition $X(t_0) = x_0$. In Eq. (1), $X(t)$ is the mutant allele frequency of the underlying population and $W(t)$ is a standard Brownian motion. An excellent theoretical introduction to the Wright-Fisher diffusion with selection can be found in Durrett (2008). The observations are modelled as independent binomial samples drawn from the underlying population at each given time point. We assume that the available data are always sampled from the underlying population at a finite number of distinct time points, say $t_1 < t_2 < \ldots < t_K$, measured in units of $2N$ generations. At the $k$-th sampling time point, there are $u_k$ mutant alleles and $v_k$ ancestral alleles observed in the sample of $n_k$ chromosomes drawn from the underlying population.

The population genetic quantities of interest are the scaled selection coefficient $\alpha$ and the dominance parameter $h$. We let $\boldsymbol{x}_{1:K} = (x_1, x_2, \ldots, x_K)$ represent the mutant allele frequency trajectory of the underlying population at the sampling time points $\boldsymbol{t}_{1:K}$, then the posterior probability distribution for the population genetic quantities of interest can be formulated as

$$p\left(\alpha, h \mid \boldsymbol{u}_{1:K}, \boldsymbol{v}_{1:K}\right) = \int p\left(\alpha, h, \boldsymbol{x}_{1:K} \mid \boldsymbol{u}_{1:K}, \boldsymbol{v}_{1:K}\right) \, d\boldsymbol{x}_{1:K},$$

where

$$p\left(\alpha, h, \boldsymbol{x}_{1:K} \mid \boldsymbol{u}_{1:K}, \boldsymbol{v}_{1:K}\right) \propto p\left(\alpha, h\right) p\left(\boldsymbol{x}_{1:K} \mid \alpha, h\right) p\left(\boldsymbol{u}_{1:K}, \boldsymbol{v}_{1:K} \mid \boldsymbol{x}_{1:K}\right). \tag{2}$$

In Eq. (2), the first term $p(\alpha, h)$ is the prior probability distribution for the population genetic quantities of interest and can be taken to be a uniform prior over the parameter space if their prior knowledge is poor. The second term $p(\boldsymbol{x}_{1:K} \mid \alpha, h)$ is the probability distribution for the mutant allele frequency trajectory of the underlying population at the sampling time points

$\boldsymbol{t}_{1:K}$ and can be written as

$$p\left(\boldsymbol{x}_{1:K} \mid \alpha, h\right) = p\left(x_1 \mid \alpha, h\right) \prod_{k=1}^{K-1} p\left(x_{k+1} \mid x_k; \alpha, h\right),$$

where $p(x_1 \mid \alpha, h)$ is the prior probability distribution for the mutant allele frequency of the underlying population at the initial sampling time point and can be set to be a uniform prior over the state space $[0, 1]$ if its prior knowledge is poor, and $p(x_{k+1} \mid x_k; \alpha, h)$ is the transition probability density function of the Wright-Fisher diffusion between two consecutive sampling time points for $k = 1, 2, \ldots, K-1$. The third term $p(\boldsymbol{u}_{1:K}, \boldsymbol{v}_{1:K} \mid \boldsymbol{x}_{1:K})$ is the conditional probability for the observations at the sampling time points $\boldsymbol{t}_{1:K}$ given the mutant allele frequency trajectory of the underlying population and can be decomposed as

$$p\left(\boldsymbol{u}_{1:K}, \boldsymbol{v}_{1:K} \mid \boldsymbol{x}_{1:K}\right) = \prod_{k=1}^{K} p\left(u_k, v_k \mid x_k\right).$$

Let $\phi$ be the rate of the sampled chromosomes that contain variants with unknown alleles at each locus, and then we have

$$p\left(u_k, v_k \mid x_k\right) = \sum_{z_k=u_k}^{n_k-v_k} p\left(z_k \mid x_k\right) p\left(u_k, v_k \mid z_k\right),$$

where

$$p\left(z_k \mid x_k\right) = \frac{n_k!}{z_k!(n_k - z_k)!} x_k^{z_k} (1 - x_k)^{n_k - z_k}$$

and

$$p\left(u_k, v_k \mid z_k\right) = \frac{z_k!(n_k - z_k)!}{u_k!(z_k - u_k)! v_k!(n_k - z_k - v_k)!} \phi^{n_k - (u_k + v_k)} (1 - \phi)^{u_k + v_k}.$$

To compute the posterior probability distribution $p(\alpha, h \mid \boldsymbol{u}_{1:K}, \boldsymbol{v}_{1:K})$, we employ a special case of the PMMH algorithm, where we do not generate and store the mutant allele frequency trajectory of the underlying population in the state of the Markov chain. More specifically, our Bayesian inference procedure proceeds as follows:

Step 1: Draw initial candidates of the parameters $\alpha^{(1)}, h^{(1)} \sim p(\alpha, h)$.

Step 2: Calculate the marginal likelihood $p(\boldsymbol{u}_{1:K}, \boldsymbol{v}_{1:K} \mid \alpha^{(1)}, h^{(1)})$.

Repeat Step 3 until sufficient samples of the parameters $\alpha$ and $h$ have been obtained:

Step 3.1: Draw new candidates of the parameters $\alpha^{(i+1)}, h^{(i+1)} \sim q(\alpha, h \mid \alpha^{(i)}, h^{(i)})$.

Step 3.2: Calculate the marginal likelihood $p(\boldsymbol{u}_{1:K}, \boldsymbol{v}_{1:K} \mid \alpha^{(i+1)}, h^{(i+1)})$.

Step 3.3: Accept proposed candidates of the parameters $\alpha^{(i+1)}$ and $h^{(i+1)}$ with the Metropolis-Hastings ratio

$$A = \frac{p(\alpha^{(i+1)}, h^{(i+1)})}{p(\alpha^{(i)}, h^{(i)})} \frac{p(\boldsymbol{u}_{1:K}, \boldsymbol{v}_{1:K} \mid \alpha^{(i+1)}, h^{(i+1)})}{p(\boldsymbol{u}_{1:K}, \boldsymbol{v}_{1:K} \mid \alpha^{(i)}, h^{(i)})} \frac{q(\alpha^{(i)}, h^{(i)} \mid \alpha^{(i+1)}, h^{(i+1)})}{q(\alpha^{(i+1)}, h^{(i+1)} \mid \alpha^{(i)}, h^{(i)})}.$$

Given the values of the parameters $\alpha$ and $h$, the marginal likelihood $p(\boldsymbol{u}_{1:K}, \boldsymbol{v}_{1:K} \mid \alpha, h)$ can be obtained by running the bootstrap particle filter introduced by Gordon et al. (1993). More specifically, the marginal likelihood $p(\boldsymbol{u}_{1:K}, \boldsymbol{v}_{1:K} \mid \alpha, h)$ is estimated by SMC, which is equal to the product of average weights of all particles at the sampling time points $\boldsymbol{t}_{1:K}$. In the bootstrap particle filter, we generate particles from the Wright-Fisher diffusion with the Euler-Maruyama scheme. Full details of the PMMH algorithm can be found in Andrieu et al. (2010).

Once we obtain enough samples of the parameters $\alpha$ and $h$, we can get the MMSE estimates for the population genetic quantities of interest. Alternatively, we can approximate the posterior probability distribution $p(\alpha, h \mid \boldsymbol{u}_{1:K}, \boldsymbol{v}_{1:K})$ with the samples of the parameters $\alpha$ and $h$ and achieve the MAP estimates for the population genetic quantities of interest.
